# Capturing the distribution as it shifts: chile pepper (*Capsicum annuum* L.) domestication gradient meets geography

**DOI:** 10.1101/2022.11.29.518324

**Authors:** NE Martínez-Ainsworth, H Scheppler, A Moreno-Letelier, V Bernau, MB Kantar, KL Mercer, L Jardón-Barbolla

## Abstract

**Aim:** Domestication is an ongoing well-described process. However, while many have studied the changes domestication causes in the genetic landscape, few have explored the way domestication changes the geographic landscape in which the plants exist. Therefore, the goal of this study was to understand how the domestication status changed the suitable geographic space of chile pepper in its center of origin.

**Methods:** *Capsicum annuum* is a major crop species globally whose domestication center, Mexico, has been well studied. This provides a unique opportunity to explore the degree to which ranges of different domestication classes diverged and how these ranges might be altered by climate change. To this end, we created ecological niche models for four domestication classes (wild, semiwild, landrace, modern cultivar) based on present climate and future climate scenarios for 2050, 2070, and 2090.

**Results:** Considering present environment, we found substantial overlap in the geographic niches of all the domestication gradient categories. Yet, there were also clear unique environmental and geographic aspects to the current ranges. Wild and commercial varieties were at ease in desert conditions as opposed to landraces. With projections into the future, habitat was lost asymmetrically, with wild, semiwild and landraces at far greater risk than modern cultivars. Further, we identified areas where future suitability overlap between landraces and wilds is expected to decouple.

**Main conclusions:** While range expansion is widely associated with domestication, there is little support of a constant niche expansion (either in environmental or geographical space) throughout the domestication gradient. However, a shift to higher altitudes with cooler climate was identified for landraces. The clear differences in environmental adaptation, such as higher mean diurnal range and precipitation seasonality along the domestication gradient classes and their future potential range shifts show the need to increase conservation efforts, particularly to preserve landraces and semiwild genotypes.

## Introduction

Domestication drastically transformed plant species, including their interactions with the environment and other organisms, as well as their geographic distributions – (Purugganan & Fuller, 2009; Hufford et al., 2019). These evolutionary changes have occurred due to natural and human-mediated evolution, partly stimulated by modifications of their growing environment (Eriksson, 2013; Kantar et al., 2017). The results of this evolution can be observed in comparisons of domesticated cultivars and their wild relatives, which help us acquire a deeper understanding of the mechanisms and outcomes of the domestication process (Swanson-Wagner et al., 2012; Meyer & Purugganan, 2013; Pace et al., 2015; Martín-Robles et al., 2019). While wild populations are expected to be adapted to abiotic and biotic conditions typical of natural ecosystems, cultivated varieties should reflect, in part, adaptations to human intervention (Milla et al., 2015). Nuanced comparisons between wild and semiwild (i.e., weedy) forms of crop progenitors (e.g. (Kane & Rieseberg, 2008)) and between landraces and commercial varieties (e.g.(Mercer & Perales, 2019)) are also helpful to clarify the continuum of differentiation represented within a domestication gradient. Domestication gradients are especially evident in centers of origin where the continual presence of wild populations has enabled constant incorporation of wild individuals to the domestication process (Casas et al., 2007).

It is well understood at a global level that domesticated species are not restricted to the spatial range of their progenitor species. Some domesticated crops such as maize, wheat and common bean have shown extreme geographic range expansions outside their centers of origin (reviewed in Cortinovis et al., 2020). New environmental conditions encountered by maize have promoted particular adaptations (Ducrocq et al., 2008; Hufford et al., 2013; Takuno et al., 2015). However, the effect of domestication gradients on geographic distributions within the center of origin is less studied and highly relevant in terms of comprehending the original adaptation process to human cultivation.

An ideal way to approach the shifts in geographic distribution over the course of domestication is with ecological niche models (ENM). ENMs seek to describe the multidimensional niche space of a species (Soberón & Nakamura, 2009, Peterson & Soberón, 2012). For every point in a multidimensional niche space, there is one or more corresponding points (Colwell & Rangel, 2009) in physical space (Hutchinson, 1957). Such partial reciprocity has allowed for ENMs to be projected onto physical space to create maps, that is, to generate species distribution models (SDM) which have been successfully used to elucidate the geographic distribution of plants in natural (De Jesús Sánchez González et al., 2018; Khoury et al., 2020) and cultivated systems (Ramirez-Villegas et al., 2020).

For a given biological system, the dynamics of environmental differentiation among categories, such as species, subspecies, or even local varieties, may vary. For instance, wild squash species SDMs show that they have colonized wetter environments (Castellanos-Morales et al., 2018). Few ENM/SDM studies have compared both wild and domesticated species, exceptions include the Mesoamerican jocote plum (Miller & Knouft, 2006). While maize wild relatives (teosintes) show stable projections into the past, models for primitive cultivated maize landraces into the present showed expansion (Hufford et al., 2012). Methods developed to compare SDM results among different categories include niche equivalency tests and background similarity tests (Warren et al., 2008). Likewise, ENMs can be compared through visualization of expected environmental clustering with principal component analyses (PCA) (e.g (Nakazato et al., 2010)). These comparisons are useful when there is the possibility of niche divergence propelled by artificial selection.

The use of SDMs can allow for assessment of the future effects of climate change on species distributions (IPCC, 2014; Aguirre-Liguori et al., 2021). Rapid changes in projected distributions of a taxa might indicate high risk of climate stress that could corner their populations to either migrate, adapt to new local conditions, or become extinct (Jump & Peñuelas, 2005; Parmesan, 2006; Aitken et al., 2008). Similar processes may govern crops, though natural gene flow could be complemented by human-assisted migration (Mercer & Perales, 2010; Pironon et al., 2019). For maize landraces in Mexico, reduction of suitable areas under climate change scenarios was less harsh than for their wild relatives, teosinte and *Tripsacum*. Nevertheless, differential responses were identified per maize race, with vulnerable taxa, as well as new suitable areas, being detected (Ureta et al., 2012). Although distributions of distinct species or taxa within species are expected to shift in different ways few studies have tackled this issue along a domestication gradient.

Chile pepper (*Capsicum annuum* L.) is an excellent species to study shifts in environmental niche through the process of domestication in its center of origin. Of the 31 wild chile pepper species, five have been successfully domesticated, with *C. annuum* var. *annuum* domesticated in Mexico (Pickersgill, 1971; Carrizo García et al., 2016)—a center of origin hotspot (Vavlilov, 1926) with a long and rich history of chile pepper use (Aguilar-Rincón et al., 2011). Domesticated chile peppers (*C. annuum* var. *annuum*) diverge from wild forms (*C. annuum* var. *glabriusculum*) in a number of traits, such as loss of fruit abscission at maturity; loss of bird dispersal; fewer, larger and pendant fruits with more synchronous maturation; greater color/shape variability; loss of seed dormancy and nurse plant association; annualization of life cycles; reduction in cross pollination; and reduction in lateral branch number (Paran & Van Der Knaap, 2007; Luna-Ruiz et al., 2018). Archaeological findings date *Capsicum annuum* use to approximately 5,600-6,400 yBP, according to colocalization with dated maize and squash macroremains and dated starch fossils from other *Capsicum* species (Mangelsdorf et al., 1965; C. E. Smith, 1967; Long et al., 1989; B. Smith, 1997; Perry et al., 2007). Using archaeological, geographic, paleobiolinguistic and genetic evidence, Kraft *et al*., (2014) concluded that central east Mexico and/or northeastern Mexico were the most likely centers of origin for *C. annuum*; yet they could not rule out multiple domestication events, as suggested by others (Aguilar-Meléndez et al., 2009).

*C. annuum* in Mexico offers a unique testing ground for the influence of the domestication process on niche evolution, since wild, semiwild (or let-standing), landraces and commercial varieties coexist throughout the country. All four types are actively used by humans. In wild chile peppers, long-distance dispersal is mediated by birds and establishment is associated with nurse-plants’ understory (Carlo & Tewksbury, 2014). Dispersal of chile pepper landraces is expected to be partly dependent on ancient and modern farmers that could and do migrate with their crops or exchange seeds (Orozco-Ramírez et al., 2016) but see (Aguilar-Meléndez et al., 2009). Small-holder management practices are common for landrace survival in tropical regions (Perfecto & Vandermeer, 2008); they include rain-fed fields, milpas and backyards, where also spontaneous wild morphotypes are tolerated (let-standing/semi-wild). In fact, incipient domestication phenomena have been recorded for three other Mesoamerican *in situ* managed plants (Casas et al., 2007). Landrace plant varieties tend to contain high genetic variability, partly because they are the product of evolutionary interactions with agroecosystems, where natural and human-mediated selection are at play (Cortinovis et al., 2020). Commercial chile pepper production in Mexico positions this country as the second largest world producer, only after China (FAO, 2019). Mexico is world leader in chile pepper exports, and throughout the 2003-2016 period chile pepper production is estimated to have accounted for 3.2 million annual tons and 3.5 % of the national agricultural GDP, with five varieties leading national production (SAGARPA, 2017); many of these commercial varieties are grown industrial agricultural settings with access to irrigation.

A better understanding of the shift in geographic distributions and environmental space encompassed by various domestication classes in Mexican chile pepper (*C. annuum*) is lacking. Thus, we employed an environmental niche modeling analysis for the complete *C. annuum* domestication gradient – from wild and semiwild to landrace and commercial cultivars – sampled in Mexico. To this end we described environmental envelopes per category using PCA analyses and employed a correlative maximum entropy niche modeling method (Maxent) to generate and compare their SDMs. We performed these analyses based on current, as well as future climatic scenarios.

In particular, our objectives were to:

1. Discern differences in the geographic distributions of classes along a domestication gradient.
2. Clarify the environmental factors that climatically differentiate these classes.
3. Compare projected niche suitability of geographic distributions under current climatic conditions, as well as multiple near-future climate change scenarios, for each class along the domestication gradient.

We hypothesized that we would see expansion of the environmental niche and geographic distribution along the domestication gradient, indicative of human modification of the niche. We also hypothesized that climate change will be predicted to affect natural populations more severely since they lack the buffering effect derived from human management. These analyses will enhance our ability to understand the ways that the ongoing process of domestication has shaped the ecogeographic qualities of plant species. We were interested in assessing how future climatic scenarios (inherently tied to CO2 emissions scenarios) will be expected to affect chile pepper suitability areas along the domestication gradient. These estimates can help track possible changes in overlapping regions and evaluate germplasm relevance for future climate scenarios.

## Methods

### Occurrence point classification system

Occurrence points for *C. annuum* were classified as wild, semiwild, landrace or commercial subsets. Our wild category included *C. annuum* var. *glabriusculum* and was defined to occur in natural habitats and exhibit wild phenotypic traits without any sign of chile pepper domestication syndrome.

Semiwild (*C. annuum* var. *glabriusculum*, mainly) were characterized as plants morphologically corresponding to *C. annuum* var. *glabriusculum* (perennial plants with small erect fruits, small flowers) that may occur in human-managed spaces such as back-yards, small family plots or *milpa* agroecosystems, where they grow spontaneously as *arvenses* (from latin word *arvum:* ploughed; i.e associated to ploughed land) and are tolerated/consumed yet retain a wild-like phenotype (Aguilar-Meléndez et al., 2009; Pérez-Martínez et al., 2022). Landraces (*C. annuum* var *annuum*, mainly) are identified as such by farmers and communities and are typically grown through traditional agriculture with a tendency to pertain to particular cultural regions in association to specific uses by local communities. Commercials (*C. annuum* var *annuum*) represent cohesive and standardized varieties that are usually grown at large scales through high-input technology-dependent agriculture. ODMAP (Overview, Data, Model, Assessment and Prediction) reporting protocol (Zurell et al., 2020) is available for this study’s distribution models in the data accessibility section. All analyses were performed in R v.4.0.2. (R Foundation for Statistical Computing, 2020) except otherwise described.

### Data collection and sampling

Occurrence points for all four domestication categories were obtained from field trips throughout Mexico (2013-2015 and 2018-2019) on behalf of Mercer (Ohio State University - OSU) and Jardón (National Autonomous University of Mexico - UNAM) labs, comprising a collection of 1154 samples. Wild points were subsequently enriched with occurrences reported for Mexico in (Kraft et al., 2014) and (Khoury et al., 2020) as well as public databases: CONABIO (National Commission for the Knowledge and Use of Biodiversity) through SNIB-REMIB (National Information System on Biodiversity-World Network of Information on Biodiversity) up to 2014. Additional vouchers from MEXU (UNAM) herbarium accessions were also included. Finally, a curated database of wild *C. annuum* var. *glabriusculum* compiled by CONABIO and analyzed by (Goettsch et al., 2021) was added. Semiwild collection points were complemented with coordinates from (Aguilar-Meléndez et al., 2009) and (González-Jara et al., 2011). For cultivated varieties, proxies of occurrence points were estimated taking information from municipalities reported by (SAGARPA, 2018) to produce chile pepper through irrigated and non-irrigated cultivation. These municipalities were cropped to include only agricultural polygons as reported by (INEGI, 2017) soil-use layer series VI. For both irrigated and non-irrigated categories random points were generated within the retrieved polygons and used as coordinate proxies of true occurrences. The landrace subset included collection landrace points plus non-irrigated polygon points. The commercial subset was built with collection commercial points plus both irrigated and non-irrigated polygon points. Finally, two broad domestication categories were conceived, wild *sensu lato* (wild-sl), displayed wild-type phenotypes thus including wild and semiwild subcategories, whereas the cultivated group added landraces and commercial subcategories plus SNIB-REMIB CONABIO accessions for cultivated chile peppers that couldn’t be classified as either landrace or commercial. The complete data comprised 8719 occurrence points (Supp Fig 1, see Supp Table 1 for the breakdown by category).

Occurrence data was processed and formatted for each domestication group. For all data points, we extracted their corresponding values for 19 bioclimatic variables from layers averaged across 1970-2000, retrieved from WorldClim (Fick & Hijmans, 2017) at 2.5 arc-min resolution (approximately 5 km). 17 soil variables were obtained from ISRIC (Hengl et al., 2017) at 2.5arc-min, and slope/aspect values were also obtained from WorldClim.

### Environmental niche analyses through principal components

Principal component analysis (PCA) is commonly used to explore environmental niches, as the components of the environment niche are made up of many different variables (Robertson et al., 2001; Di Cola et al., 2017). We selected a subset of variables as described in Appendix 1. We conducted PCAs with FactoMineR v.2.3 (Husson et al., 2016) on 9 selected bioclim variables for the whole dataset. (BIOS:2, 3, 4, 5, 9, 14, 15, 18 and 19) (Supp Table 2). Pairwise comparisons were then plotted for domestication categories, within broad and stepwise classes, along the first two principal components. Minimum convex hulls were traced for each category (Fig 1, Supp Fig 2); their areas, the overlap of areas and the percentage of overlapping area for each category were calculated (Supp Fig 3, Supp Table 3). To account for data point clustering and uneven sampling, we also calculated the percentage of points from each category found in the overlapping area (Supp Table 3). Violin plots were generated for each variable in each domestication category to compare dispersion features (Fig 2, Supp Fig 4).

**# Figure_1.**
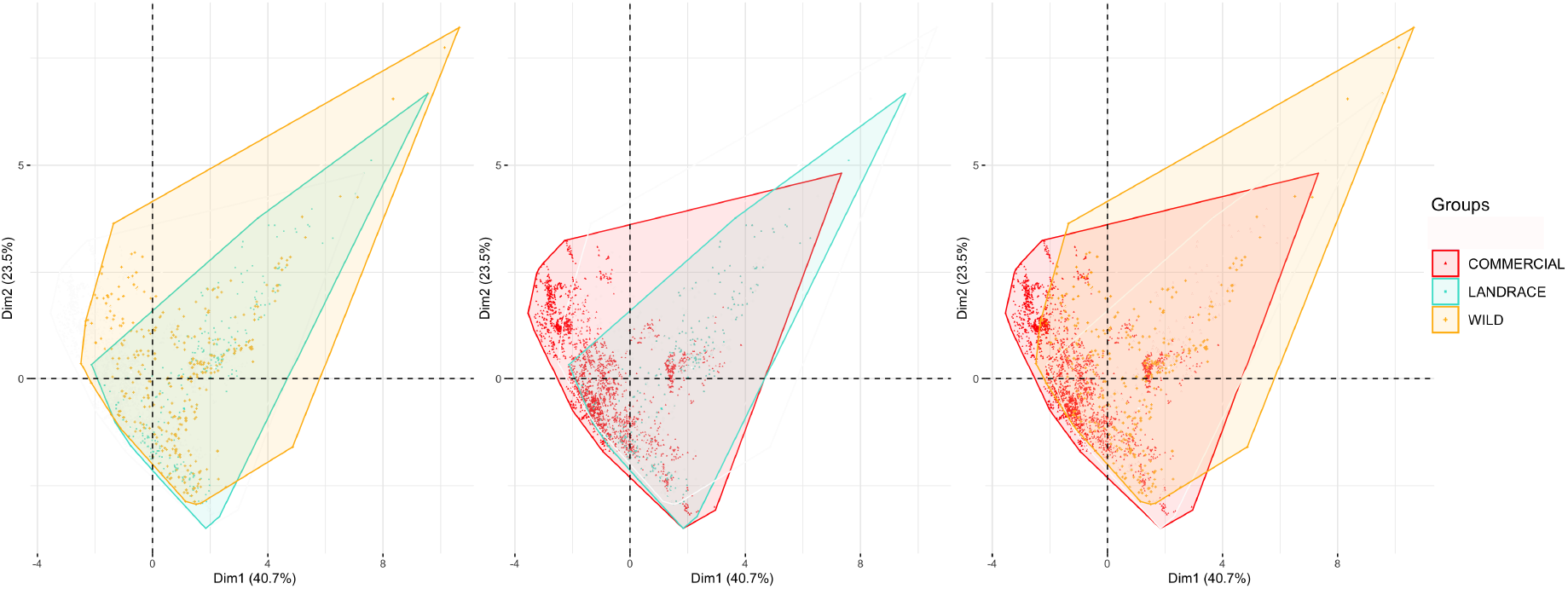
PCA projections on PC1 and PC2 axes for pairwise comparisons of domestication categories. Minimum convex polygons are traced for each category.

**# Figure_2.**
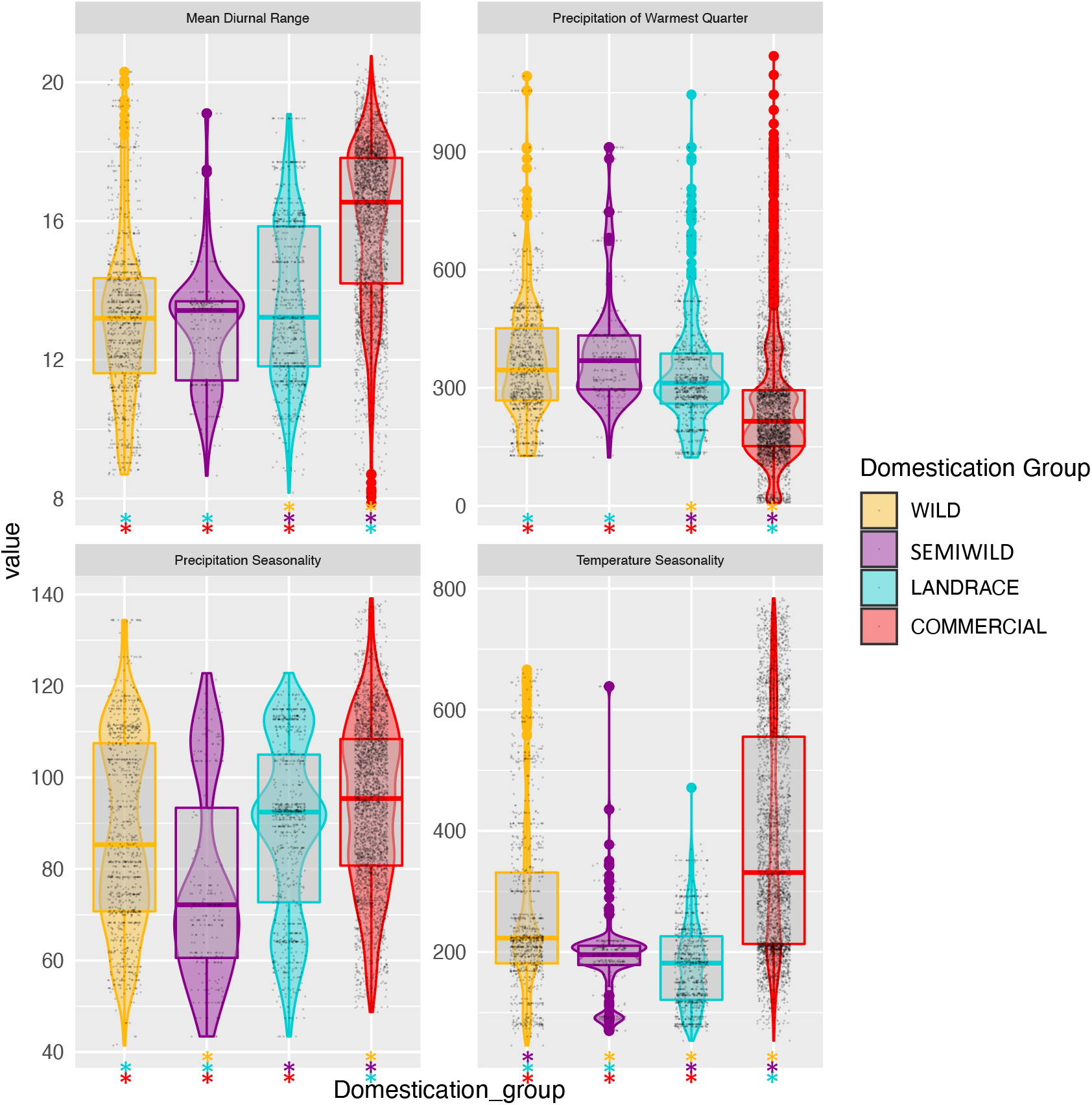
Dispersion of values for four contrasting variables between domestication groups. Boxes within violin plots mark the median and 25/75 percentiles. For mean dirunal range and precipitation of the warmest quarter Wilds are not significantly different from arvenses (bootstrap resampling CI) and for precipitation seasonality Wilds and Landraces have overlapping CI.

### Environmental niche modeling

For each domestication category in *C. annuum*, we modeled the environmental niche and performed geographic projections of niche suitability with the maximum entropy algorithm Maxent (Phillips et al., 2006). We selected Maxent because it is ideal for presence-only data sets and employs a correlative method that has high predictive accuracy (Phillips et al 2006, Elith et al 2011, Merow et al 2013). Environmental layers of the previously selected variables at 2.5 arc-min resolution were employed for training and testing of the models. They were cropped to a minimum convex polygon, comprising all data points together (thinned at 5km), and buffered at three degrees. For further details on data preparation and model tunning see Appendix 1.

MaxEnt runs were performed on thinned datasets with the following specifications. Feature class LQHP was selected for all datasets. Regularization multiplier was set to three for wild, wild *s.l*., semiwild and landrace datasets and one for commercial and cultivated datasets (Supp Table 4 and Supp Fig A3). Clamping was used to account for the artificial sampling barrier we established by including only points within Mexico, a common approach when there are known restrictions on distributions (Stohlgren et al., 2011). We used 10,000 background points and 30% random test points, with 500 maximum iterations. Ten replicates were run using cross validation, and Jackknife tests were requested. Logistic output format projections for the custom-box clipped layers were calculated and the median of the ten replicates were plotted (Supp Fig 5a). Background predictions were logged for downstream True Skill Statistic (TSS) (Allouche et al., 2006) calculations for each mean model. Mean AUC and ROC curves were inspected for the ten-replicate averaged models to ensure consistency among replicates. We compared MaxEnt’s output tables for average permutation importance of variables among domestication categories through stacked barplots (Supp Fig 6). Although variable percent contribution explains how much a variable contributed to a model run, permutation importance measures how changing the variable’s values affects AUC, and thus its importance for the final model (Songer et al., 2012).

We used ENMTools v.1.0.5 (Warren et al., 2010) to calculate three pairwise niche similarity tests between domestication categories: Schoener’s D, Hellinger’s I (normalized by Warren) and Spearman rank correlation. These statistics are intended to quantify niche overlap and are calculated from probability distributions assigned to reticulate geographic space. Schoener’s D assumes that the probability of occurrence in a given cell is related to resource use and thus species local density, whereas Helinger’s I is free from such assumption (Warren et al., 2008). We assessed spatial bias of our different sampling strategies by generating a spatial bias file (Fourcade et al., 2014) buffered at 1 degree for each dataset, and cropped each environmental raster accordingly. MaxEnt models using these files produced over-projections, indicating decently spread sampling effort throughout study areas. To further complement the approach, and in view of our sampling biases among datasets, we decided to measure ecological niche overlap between wild-landrace and wild-cultivated pairs in environmental space (Broennimann et al., 2012), taking random distributions of variables from the predicted distribution to calculate D and I. To better compare the resulting MaxEnt suitability matrices on geographic space, we transformed each median map into binary color codes by calculating a specific threshold for each data set. See Appendix 1 for details.

### Niche projection for future climate scenarios

We were interested in projecting each domestication category’s niche under future climate scenarios to suggest how these might be differently affected and how potential geographical shifts could modify the amount and location of overlapping regions among categories.

Each of the ten replicate models obtained per dataset was independently projected onto future climates (Fick and Hijmans, 2017) CMIP6 (Coordinated Model Intercomparison Project Phase 6) (Eyring et al., 2016) using terminal commands (using the same layers with the same extent and resolution as present-day models; Appendix 1). The future layers included scenario averages for temporal intervals of years 2041-2060, 2061-2080 and 2081-2100, under two shared socioeconomic pathways (SSPs): 2 (“middle of the road”) with representative concentration pathway (RCP) of 45, and 5 (“fossil-fuled development”) with RCP of 85. For each of the afore-mentioned combinations, the eight general circulation models (GCMs) that were available for CMIP6 were run (BCC-CSM2-MR, CNRM-CM6-1, CNRM-ESM2-1, CanESM5, IPSL-CM6A-LR, MIROC-ES2L, MIROC6 and MRI-ESM2-0). GCMs simulate the relationships between atmosphere, oceans, land, and ice, to try to understand how CO2 would impact temperature and how temperature affects the other climatic variables (precipitation, could cover, wind etc.). Mean, median and standard deviation ASCII maps were obtained for each set of ten replicate future projections per configuration in R (data not shown).

Binary maps were obtained for projections of niche suitability under future climate scenarios to allow an easier comparison of differences and overlaps. Thresholds on future projections per domestication category were obtained from the present time models of each dataset and applied to each year-SSP-GCM model median map. Within each domestication category, pixel overlap metrics (loss, gain and shift) were calculated for each future projection as compared to their present projections, for each GCM as well as their intersection (Fig 4, Supp Fig 7, Supp Table 5). In addition, equivalent calculations were obtained for pairwise domestication classes overlaps (Supp Table 6). To better visualize future scenarios results, we used the intersection of the 8 GCMs and compared this to present-day projections and future overlaps between domestication categories as described in Appendix 1.

ENM sources of uncertainty are manyfold, and literature revising niche model projections’ transferability in time mention type of future scenario selected and migration potential, among the main issues (Sequeira et al., 2018; Yates et al., 2018), with caution being summoned when non-analog combinations of conditions are expected (Fitzpatrick & Hargrove, 2009; Zurell et al., 2012). For this reason, (Qiao et al., 2019) suggests environmental similarity be calculated to assess uncertainty in model transferability. Thus, we calculated Multivariate Environmental Similarity Surfaces (MESS) (Elith et al., 2011) between our present and future projections and their layers.

## Results

### Environmental niche analyses through principal components

We built a final Mexican chile pepper database comprising 8519 occurrence points classified into 593 wild, 96 semiwild, 298 landrace and 3816 commercial occurrences. In the PCA analyses on the four fine-grained categories (wilds, semiwild, landraces and commercials), first and second principal components (PC1 and PC2) accounted for 40.7% and 23.5% of the variance respectively (Fig 1, Supp Fig 2). Concentrating on the quadrants where pairwise minimum convex polygons mainly differed, we found that PC1 loadings displayed strong negative correlation with the mean diurnal range and temperature seasonality whereas PC2 showed high positive correlation with temperature seasonality and precipitation of the coldest quarter (Supp Table 2). Landrace PCA polygon area was mostly contained within the wild polygon (Fig 1-a); wild-landrace polygons overlapped by 76%, with 88% of landrace area shared with wild and 67% of the wild area shared with landraces (rounded figures). Hull point density overlap estimates were 99 and 89% for landrace and wilds, respectively (Supp Table 3). The wild data points that fell outside the landrace polygon corresponded to a few hot and dry northern regions (Supp Fig 3-a). For the landrace-commercial pair, we observed that very few landrace data points from very humid regions were not encompassed by commercial varieties’ hull envelope. Areas exclusive of commercials expanded in the negative PC1 and positive PC2 quadrant, corresponding to some of the country’s northern desert regions (77% PC1-PC2 area overlap and 72% PC1-PC2 point density overlap, Fig 1-b, Supp Fig 3-b). When comparing the extremes of the domestication gradient (wilds vs commercials), we found that commercial varieties have greatly re-colonized wild hot and dry conditions. Moreover, commercial varieties have managed to withstand even hotter and dryer values found in the inland northwestern deserts, (Fig 1-c, Supp, Supp Fig 3-c).

We then inquired about the distribution of the most prominent environmental variables that contrasted for each fine-grain domestication category’s occurrence points (Supp Fig 4). For example, the average precipitation of the warmest quarter gradually diminished from wild to commercials, passing through landraces (Fig. 2), whereas mean diurnal range increased. Precipitation seasonality also behaved incrementally from wilds to commercials. Commercial varieties, however, encountered more seasonality in rainfall and temperature than either of the other classes (Fig. 2). Landraces were particularly sensitive to temperature seasonality and were restricted to the lowest ranges, while commercials showed considerable dispersion with the highest median value (Fig. 2).

### Environmental niche modeling

Our first maxent model projection compared wild-sl with cultivated broad categories. From the threshold-cropped binary map, it was evident that each grouping differed geographically and was somewhat complementary, although there was 52.9% of overlap between wild-sl and cultivated (Fig. 3-a). Suitable areas for Cultivated chiles extended into dry climate and desert regions whilst wild-sl suitability could be found in highly humid regions as well as along most coasts. At fine grain categories, landraces mainly overlapped with the wild suitable areas from central to southeastern Mexico except for swamp-marsh sections of Tabasco state and southern Caribbean coast, where only wilds found suitable conditions (Fig. 6-a). Noteworthy were two exclusive regions for landraces: the Trans-Mexican Volcanic Belt (TMVB) and Guatemalan Sierras, both distinguished by their high altitude. Landraces did not have larger distributions with respect to wilds. When comparing landrace and commercial classes, commercials found high suitability in northern desert regions where landraces lacked suitable areas (Fig. 3-b). However, tropical coasts of Guerrero, Oaxaca and Quintana Roo, as well as some humid regions of Chiapas and Tabasco states, were assigned exclusively to landraces and tended to overlap with wilds (Fig. 6-a).

**# Figure_3.**
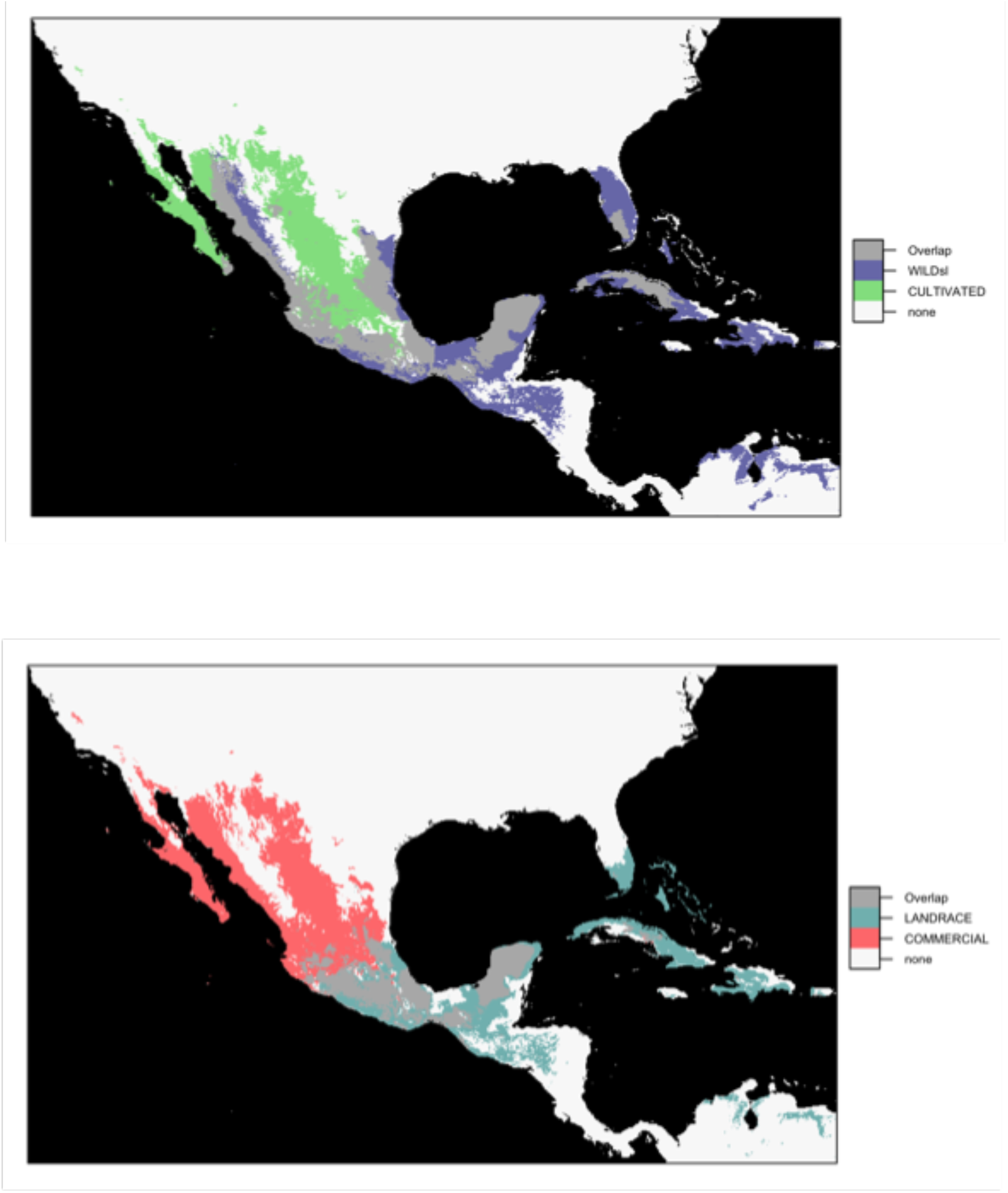
**A.** Overlap of suitability maps for *C. annuum* Wild *sensu lato* (Wilds + Arvenses) and Cultivated (Local varieties + Commercial varieties + varieties that are cultivated but couldn’t be assigned to Local or Commercial subgroupings) niche projections. Median from 10 replicates of logistic *maxent* outputs. Colored areas correspond to pixels above the 10^th^ percentile threshold of training points for each domestication category. Gray areas show projection overlap between niches. **B.** Overlap of suitability maps for *C. annuum* Landrace and Commercial niche projections. Median from 10 replicates of logistic *maxent* outputs. Colored areas correspond to pixels above the 10^th^ percentile threshold of training points for each domestication category. Gray areas show projection overlap between niches.

**# Figure_4.**
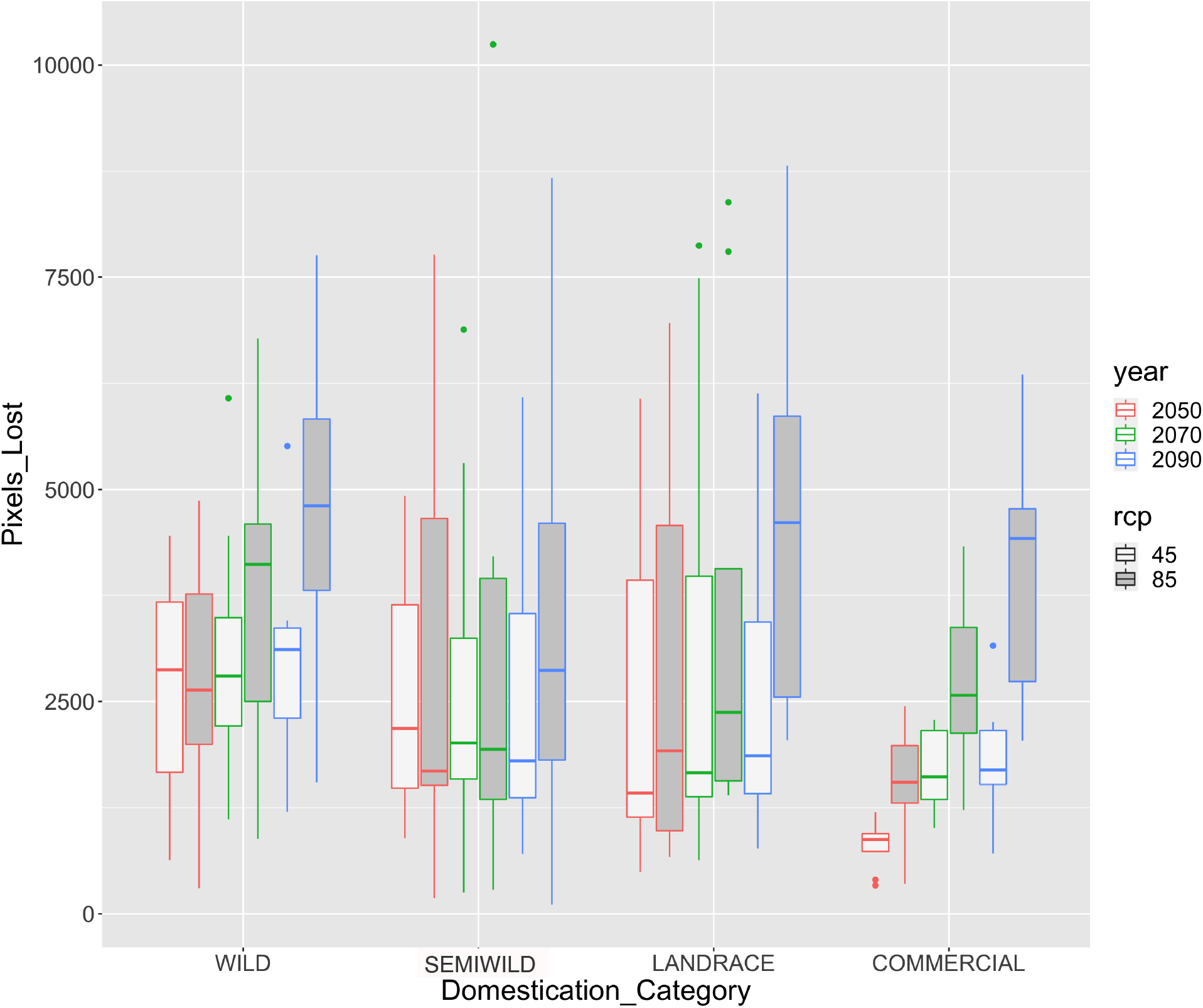
Lost area by future niche projections within each domestication group with respect to their niche projections at present. Boxplots represent dispersion in the amount of area lost according to each of the eight general circulation models employed. Boxes are outlined with color codes for future projection years and filled according to SSP 2-4.5 and SSP 5-8.5 (rcp 45 and 85 respectively). The same 10^th^ percentile threshold of training points at present was used to plot binary maps for each domestication category’s future projections and calculate overlap.

**# Figure_5.**
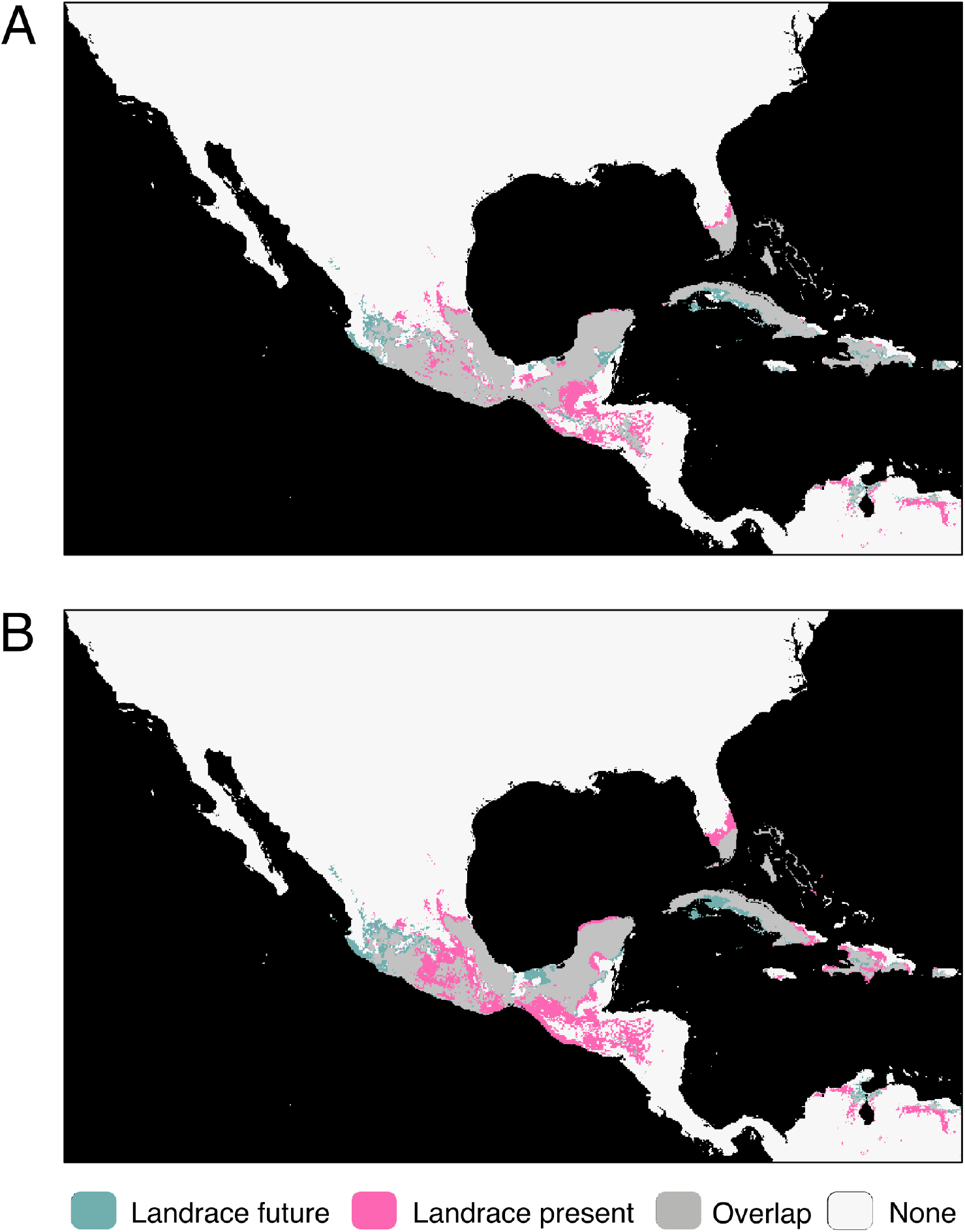
Overlap between present and future projections (year 2090) for Landrace niche under (A) SSP 2-4.5 and (B) SSP 5-8.5. Future area was obtained as the shared area among all eight GCM models, with the same 10^th^ percentile threshold from training points of the present niche model used to plot binary maps each future projection and calculate overlap among GCMs.

**# Figure_6.**
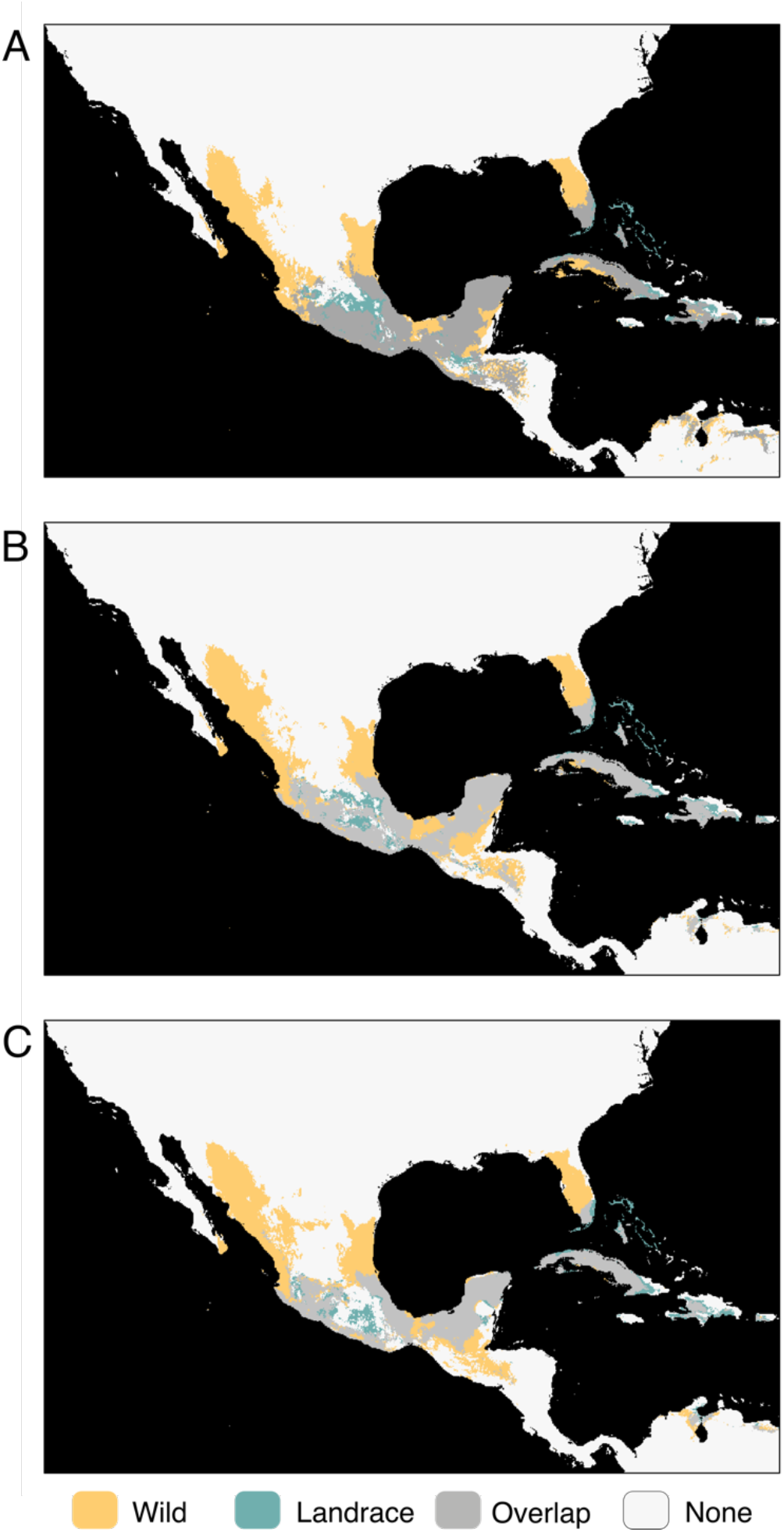
Overlap of suitability maps for *C. annuum* Wild and Landrace niche projections for present conditions (A), future 2090 conditions under SSP 2-4.5 (B) and future 2090 conditions under SSP 5-8.5. Median from 10 replicates of logistic *maxent* outputs. Colored areas correspond to pixels above the 10^th^ percentile threshold of training points for each domestication category. Gray areas show projection overlap between niches. For each domestication category, future area was obtained as the shared area among all eight GCM models, with the same 10^th^ percentile threshold from training points of the present niche model used to plot binary maps each future projection and calculate overlap among GCMs.

When we analyzed variable permutation importance, we observed that wild and wild-sl categories were strongly influenced by the precipitation of the warmest quarter, whereas semiwild and landraces were dominated by temperature seasonality (Supp Fig 6). In these four cases the described variables also showed a high contribution to each model through the jackknife test, as did temperature seasonality for wilds and wild-sl (Supp Table 7). Precipitation of the driest month showed considerable permutation importance for landraces. With the exception of mean diurnal range, the aforementioned variables had strong contributions and loadings in the PCA (Supp Table 2).

Pairwise niche similarity tests indicated that among the four fine grained category pairs, landrace and semiwild were the most similar (D=0.79, I=0.95) and commercial and landrace were the least similar (D=0 .49, I=0.75) (Supp Table8). Landraces and wilds displayed intermediate values of similarity (D=0 .68, I=0.89) that were higher than the commercials and wilds comparison. Further analyses through background tests on wild-landrace and wild-commercial pairs clarified that the landrace environmental niche was indeed nested within the wild niche but not vice versa, and that the commercial and wild niches are less similar than random expectations, thus significantly divergent (Supp Fig8).

### Niche projection for future climate scenarios

For conservation purposes and maintaining the evolutionary processes ongoing in chile pepper, we decided to examine the niche projections of each domestication category under a spectrum of future climate scenarios (i.e., different mitigation strategies and GCMs). Within each domestication class, we visualized the amount of lost area per year-SSP combinations (Fig 4). Although we are aware of the dispersion generated by the eight GCMs, we observed that the amount of original retained area diminished over time and at a faster rate for SSP 5_85, for all categories except for semiwild. For a given year, results by SSP scenario showed variation in area loss among GCMs, yet as a general trend, the Wild category reported the highest area loss (Fig 4).

To identify geographic areas under potential climate change threat, we used the intersection of all eight GCMs per year-SSP scenario and calculated the area overlap with present projections. We drew particular attention to landraces, which for year 2090 under SSP 2_45 suggested a loss of suitability in Guatemalan regions, and under SSP 5_85, an increased vulnerability is foreseen in the highlands (both Trans-Mexican Volcanic Belt--TMVB and Guatemalan sierras), as well as the pacific coast of the Isthmus of Tehuantepec (Figs 5-a and 5-b; proliferation of pink area).

We took notice of potentially vulnerable landraces that were found in the aforementioned “lost” areas (Supp Table 9). Landraces at risk included, Maax-ik from backyards and Tabaquero from milpas in Campeche; Chile de Árbol, Chilito and Miraparriba backyard landraces in Chiapas; and Bolita, Paradito, Chilgole, Guajillo, Jalapeño, Mirasol, Pijita, and Tusta backyard landraces, as well as milpa-grown Huacle, Taviche and Tusta and plantation-grown Costeño Rojo and Chile de Agua in Oaxaca. Additionally, locally unique landraces at risk included Querétaro’s backyard Chile Criollo, Tabasco’s forest-thriving Garbanzo and the Yucatán milpa’s Chile Dulce and Xcatic. When assessing the change in area of overlap between wilds and landraces at present vs future (e.g., 2090) projections, we found that the sierras of Guerrero and Oaxaca were particularly prone to loss of overlap due to reductions in both wild and landrace suitability (Fig. 6; loss of gray area). Overlap in Guatemala was also suggested to decline in the future due only to reductions in landrace suitability.

We worked on temporal transfer of model projections, so it was important for us to assess if the resulting vulnerable areas corresponded to non-analog climate combinations. To this end, we contrasted our maps of future area loss and future overlap loss among classes with MESS maps generated for each year-SSP-GCM combination (Supp Fig 9; novel combinations: red shading). For the MESS projections to 2090, novel climate combinations were scarce (SSP 5_85) or generally absent (SSP 2_45) in the regions with landrace area loss or overlap loss between landraces and wilds. When MESS was estimated for landrace niche model for 2090 only, novel combinations included GCM models IPSL-CM6A-LR and MRI-ESM2-0, for SSP 5_85, and only MRI-ESM2-0 for SSP 2_45 (Supp Fig 10-b). The same was true for wilds with the addition of two small patches in Tabasco and Michoacán states for 5_85 BCC-CSM2-MR.

## Discussion

Humans have interacted with crop plants for many millennia. These interactions are often portrayed as discrete in time and space, yet the reality is much more complex. There has been continuous cultivation of chile pepper for thousands of years. However, it is hard to discern the environments in which that cultivation has occurred. Archaeological evidence provides clues but is incomplete since plant remains are only recoverable under strict conditions (Zohary et al., 2012;Cabanes & Shahack-Gross, 2015). Clarifying the climate envelope within which different domestication classes currently reside helps provide clues on expansion or contraction of the ecological niche of chile pepper as it was domesticated, spread, and improved. With our extensive sampling across the center of origin, we explored niche changes across the domestication gradient (a proxy for the historical change), but also how the distribution of domestication classes may change in the future using the latest IPCC climate models.

### Human management has shaped the ecological niche of cultivated pepper

Literature has suggested that small-holder practices – especially polyculture – may stabilize the environmental conditions experienced by crops (Altieri et al., 2015). However, landraces were distinguished by requiring a certain degree of climatic stability (lowest temperature seasonality and highest isothermality values) (Fig 2, Supp Fig 4). Commercials, by contrast, were exposed to high precipitation seasonality. Precipitation of the driest month had high permutation importance for landraces, presumably attributable to the fact that many landrace points came from rain-fed management regimes. Thus, we suggest that management practices typical of landraces exert milder environmental buffering than those of high input agriculture. The presence of commercial chile peppers in dry areas on the suitability maps (Fig 3-b, Supp Fig 5) and their low PCA values for the precipitation of the warmest quarter (Fig 2), could be explained by artificial irrigation. However, we cannot rule out that these populations might carry drought-adapted genetic variants.

Commercial varieties and wilds constitute extreme ends of the domestication gradient. Thus, we expected that their environmental niche models and distributions would show the strongest divergence. However, our niche similarity measures were lowest for the landrace-commercial pair (D=0.49, I=0.75, Supp Table 8), both of which are cultivated types. This surprising result may be due to divergence in their histories. Landraces are a product of a long cultural process of domestication, spread, and diversification within a labor-intensive management system, while commercials result from formal breeding for success within high input, technologically advanced agricultural systems. This suggests that the mildly divergent footprint from wilds to landraces was ultimately propelled with agronomic breeding efforts that allowed for the expansion of the commercial niche by reclaiming areas typical of wilds, but not exploited by landraces (presumably by means of artificial irrigation) (Figs 1, Supp Fig 5). Nevertheless, our background tests indicated that, although commercials and wilds both occupy dry areas, their suitable conditions are in fact highly differentiated, while landraces were effectively nested within commercials (Supp Fig 8).

Some species, such as maize and plum, show a gradual expansion of their environmental niche and its geographic projections along their domestication gradient (Miller & Knouft, 2006; Calfee et al., 2021; Locqueville et al., 2022; Purugganan, 2022). By contrast, our results indicate that chile pepper domestication may have involved environmental subsampling of the wild climatic and geographic distributions such that landrace distribution is now mostly nested within the wild one. Commercials, however, expanded beyond landraces and partially returned to wild areas. Had we run our analyses on a worldwide sample of *C. annuum*, we might have expected that the expansion of commercial varieties would be further exacerbated. Diffuse or multiple domestication events may complicate the gradual expansion hypothesis. Studies to date have not compellingly concluded a single chile pepper domestication event (Aguilar-Meléndez et al., 2009; Kraft et al., 2014) and wild chile peppers are distributed throughout the country (Khoury et al., 2020). Thus, the lack of environmental and geographical expansion along *C. annuum’s* domestication gradient may be due to a nonunique domestication origin.

### Domestication allowed for expansion into higher elevations

Traditional agriculture, greatly responsible for landrace preservation and continued evolution (Altieri et al., 2015; Jardón Barbolla & Benítez Kienrad, 2016) does not seem to have drastically augmented the amount of chile pepper landraces’ suitable area. Landrace geographic projections overlap with wilds projections in highly humid regions (Fig 6-a) where initial domestication likely occurred; however, they do not overlap in the highlands (e.g., the Trans-Mexican Volcanic Belt ‘TMVB’ and

Guatemalan sierras) (Fig 6-a). In other emblematic crops such as maize the acquisition of additional adaptations following their initial domestication was a fundamental factor that allowed them to colonize higher altitudes (Calfee et al., 2021). In maize, this was achieved through adaptive introgression with their high-altitude dwelling wild relative (Hufford et al., 2013). In chile pepper, besides high-altitude landraces described as old cultivars (Cao et al., 2022) we lack knowledge about this highland migration and its required genetic adaptations. However, the ecogeographic shift into high-altitude terrain was clearly a key event in chile pepper domestication. It is unlikely that the two wild pepper species found at high altitudes, *C. lanceolatum* and *C. rhomboideum* (Guatemala and Chiapas highlands) contributed necessary adaptations through inbreeding since each has 13 chromosomes (Scaldaferro et al., 2013; Tong & Bosland, 2003) not 12 like *C. annuum*.

Commercials’ considerable geographic expansion relative to other domestication classes contrasts with its exclusion from certain highly humid lowland areas (e.g., Lower Huasteca in the state of Veracruz, the Riviera Maya in the state of Quintana Roo, and the state of Chiapas) where wild and landrace suitable areas overlap (Figs 3-b and 6-a). These regions are notably dominated by subsistence production systems (De Clerck & Negreros-Castillo, 2000) farmed by small-holders who have maintained genetic diversity in landraces (Altieri et al., 2015; Dobler-Morales et al., 2020). There may be biological and social reasons for industrial production being excluded from these areas. First, these humid regions are prone to pests such as the silverleaf whitefly (*Bemisia tabaci*, Aleyrodidae), shows some resistance to the natural chemical defenses of cultivated peppers (Ballina-Gomez et al., 2013; Latournerie-Moreno et al., 2015), and a number of fungal pathogens which are often associated with monocultures in humid zones. Finally, these regions may not have the social infrastructure or land requirements to support industrial farming operations, limiting their expansion.

### Climate change may threaten unique landraces and reduce the adaptive capacity of crops

Under “business as usual” 5_85 SSP scenario with no CO2 emissions mitigation policies, all fine grain domestication categories except semiwild saw a reduction in the amount of suitable area that overlaps present-day coverage as climate change progressed over time (Fig 4). While commercials and wilds distributions were both large, the shift in the wilds’ future suitable area was stronger (Supp Fig 7-b). Yet we are wary that such strong migrations will in fact transpire because for wilds they imply fast migration and establishment in available natural settings and for cultivated varieties farming systems that would need to migrate under nationwide coordination and public policies. For these reasons, we focus the rest of our discussion of future scenarios on areas at risk.

Landraces often are locally adapted to specific environmental/field conditions and are conserved *in situ* (Galluzzi et al., 2010; Mercer & Perales, 2010), in part for the substantial adaptive genetic material they possess (Chen et al., 2017). They can act as genetic reservoirs, providing opportunities for adaptive gene flow (Jump & Peñuelas, 2005), as decribed with improved cultivars in Cucurbita (Martínez-González et al., 2021). For *C. annuum* landraces in 2090, we found a strong impact of suitable area loss in Guatemala (Fig 5). Additionally, the non-mitigated scenario (SSP=5_85) also foresaw an important rise in landrace vulnerability for highlands and the Isthmus Pacific coast (Fig 5). ENM projections in future climate scenarios benefit from considering the effects of unequal shifts of biotic interaction partners (e.g. (Mehrabi et al., 2019; Carrasco et al., 2020)). Adding biotic partner layers could deepen our results.

Landraces currently grown in the areas that we predicted will cease to be suitable, might need special attention in the forthcoming years. We found 21 such landraces (Supp Table 9). The strong representation of these in the state of Oaxaca may be due to the location of the expected shift and/or reflect denser sampling. Fortunately, vulnerable landraces in each state were accompanied by alternative populations of the same landrace that were maintained. Among vulnerable landraces detected, Chile Criollo, Tabaquero, Garbanzo and Costeño Rojo were the earliest to display signals of potential threat (from 2050). The first three have restricted distributions, whereas Costeño Rojo is popular in the larger Costa Chica region (Oaxaca and Guerrero coast) where it is grown in milpas or plantations and used by Mixtec and afro-descendant peoples (Muñoz-Zurita, 2015;*pers. obs*). The wild–resembling Garbanzo found in forests might be a valuable gene flow bridge between wild and cultivated germplasm. The historical and cultural value of some of these landraces is linked with native peoples, such as Oaxaca’s Taviche endemic to the Mixtepec region and Tusta’s association with the Zapotec and Chatino peoples from the Loixcha region; highlighting the importance of *in situ* conservation strategies within its historical cultural context (Santiago-Luna et al., 2015; Sánchez-Cortés, 2021).

Our models suggest that climate change can be expected to diminish the geographical overlap between wild and cultivated peppers. We found that wilds may be lost in the mountains of Guerrero and Oaxaca, whereas landraces may withdraw from areas in Guatemala where there is currently overlap; both wilds and landraces may retract from Puebla and southern Chiapas (Fig 6). These losses of overlap may have consequences for the evolutionary processes ongoing in this center of origin. Many crops have been reported to evolve adaptations to new climates through wild to landrace introgression (reviewed in (Janzen et al., 2019)), a process that may be hampered if a crop population’s geographical overlap is strongly diminished (Zhang et al., 2014). In chile pepper, cultivated and wild forms of *C. annuum* can easily hybridize (Eshbaugh, 2012; Pérez-Martínez et al., 2022). Often wild chile peppers establish in milpas and backyards, providing opportunities for wild to landrace gene flow (Guzmán et al., 2005; González-Jara et al., 2011; Pérez-Martínez et al., 2022). Small-holder plots that steward this evolutionary process *in situ* should be encouraged in regions where wild and landrace distributions overlap.

## Acknowledgments

The authors would like to thank all people involved in the construction, curation and maintenance of our collection data base. We thank Ana Laura Pérez-Martínez, Gabriela Martínez-Andrade, Niza Gámez, Luis Eguiarte, Leah McHale, Esther van der Knaap, and Jack McCoy for their help and assistance during the field collection. We thank MEXU National Herbarium of the National Autonomous University of Mexico as well as CONABIO for making available high quality chile pepper occurrence descriptions. We deeply thank all the Mexican farmers and native communities for the rich heritage and ongoing preservation of chile pepper germplasm and its evolutionary dynamics.

This research was funded through the USDA-AFRI, Physiology of Agricultural Plants section for support under the grant number 2017-06351, “Genetic structure and mechanisms of drought adaptation in *Capsicum”*. N.M-A thanks her postdoctoral fellowship granted by CONACyT. Additional field trips were funded by the PAPIIT grant IA-202515 (UNAM-DGAPA) and CEIICH-UNAM annual budgets. Salary and research support for KLM was also provided by state and federal funds appropriated to the Ohio Agricultural Research and Development Center; additional funding came from the Center for Applied Plant Sciences at Ohio State University.

## Data Accessibility Statement

Occurrence points by domestication category, the complete ODMAP protocol report and R-markdown code for all analyses herein included are available at the following Figshare link: Add link and Github link: Add link.

## Supporting information

Additional Supporting Information may be found online in the supporting information section at the end of this article.

## Author Contributions

NEMA: conceptualization, formal analysis, drafting, writing/revising, HS: formal analysis, writing/revising, AML: conceptualization, formal analysis, writing/revising, VB: conceptualization, writing/revising, MBK: conceptualization, writing/revising, LJB: conceptualization, writing/revising, funding acquisition, KLM: conceptualization, writing/revising, funding acquisition.

