## Supplementary_Figures for "Capturing the distribution as it shifts: chile pepper (*Capsicum annuum* L.) domestication gradient meets geography"

##########################
##### Supplementary Figures ###
##########################
**# SuppFig_1**
Maps of Mexico for each domestication category showing the data point occurrences included in this study

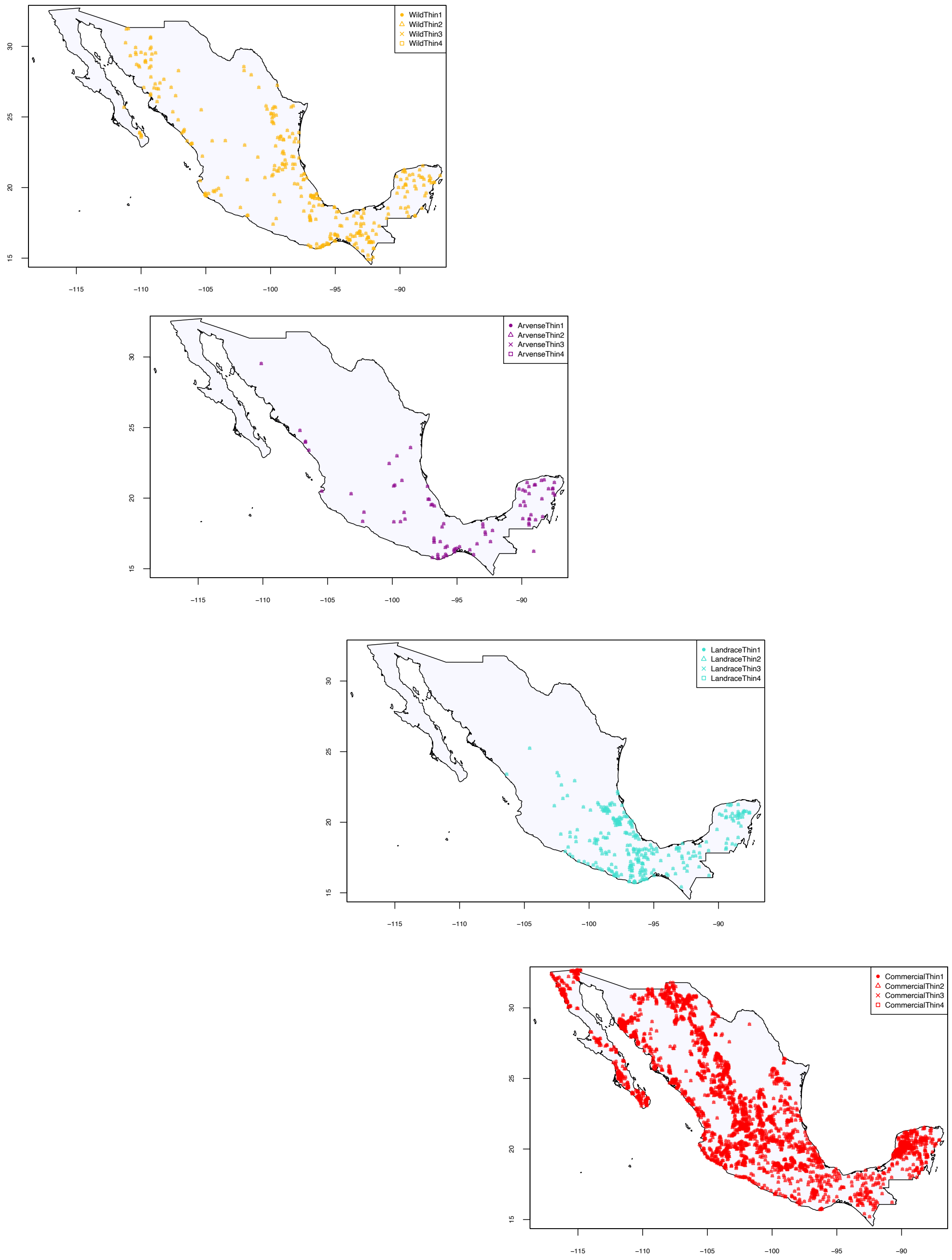

### SuppFig_2
PCA projection plots on first two principal components, showing pairwise combinations of stepwise domestication classes.

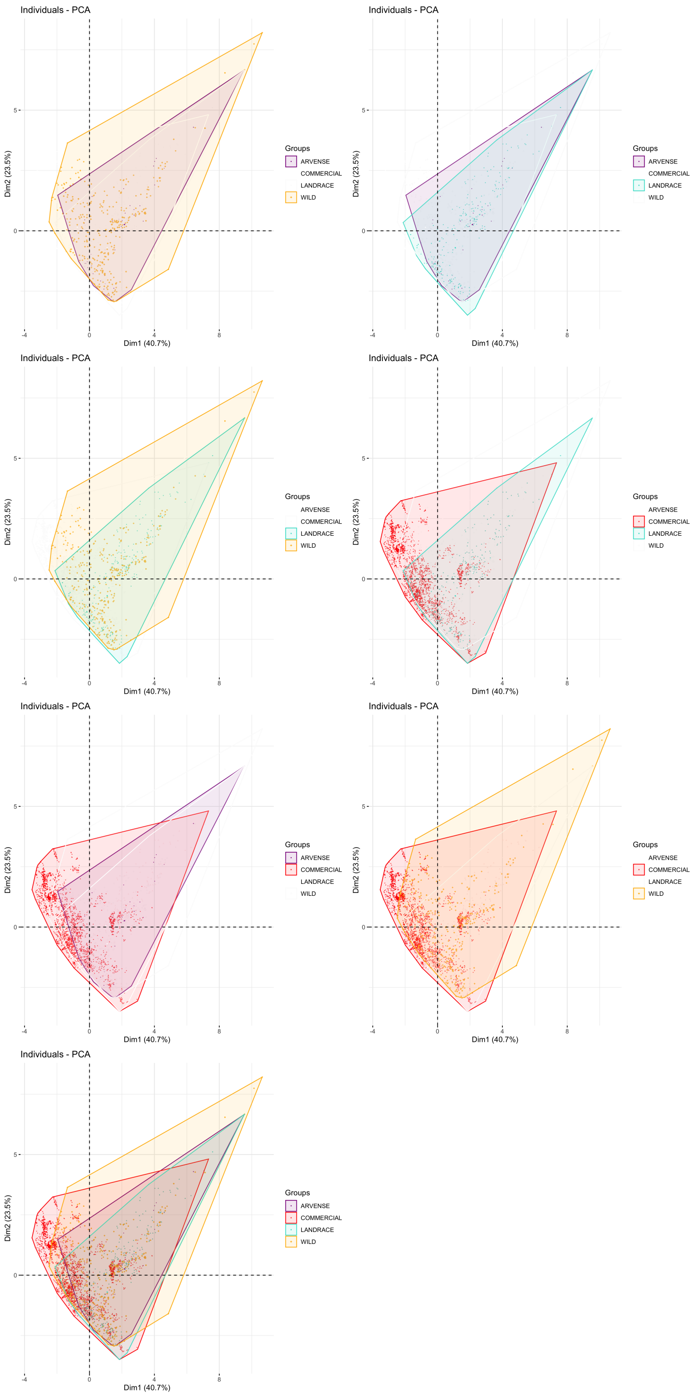

### # SuppFig_3

Maps for pairwise combinations of stepwise domestication classes depicting data points exclusive and shared to each class according to the convex hull areas plotted in the corresponding PCAs. Empty circles indicate occurrences found only in one of the classes' hull, filled circles indicate occurrences in the overlapping portion of the hulls.

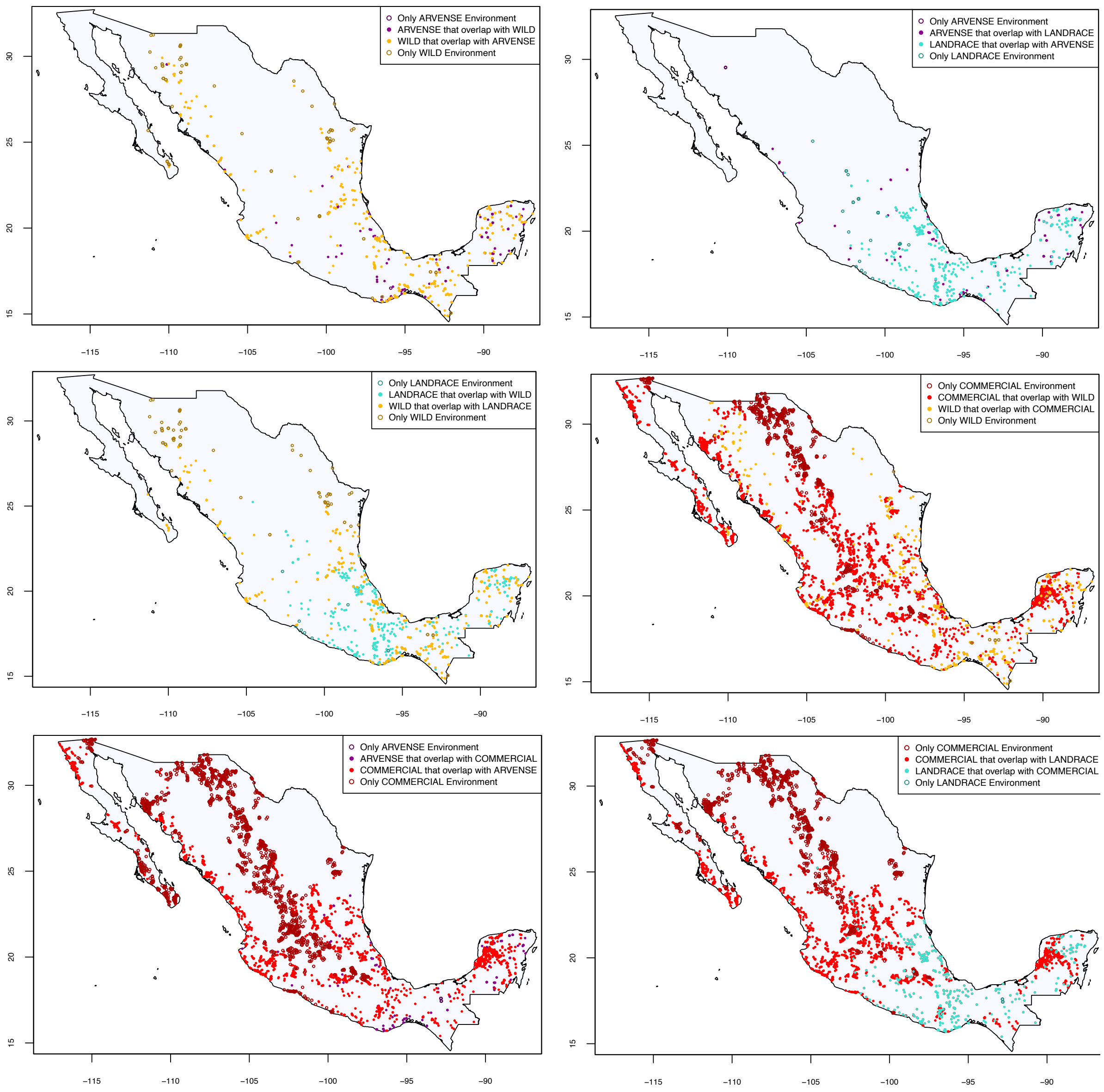

### SuppFig_4
Violin plots of the dispersal for each variable included in PCAs by domestication class.

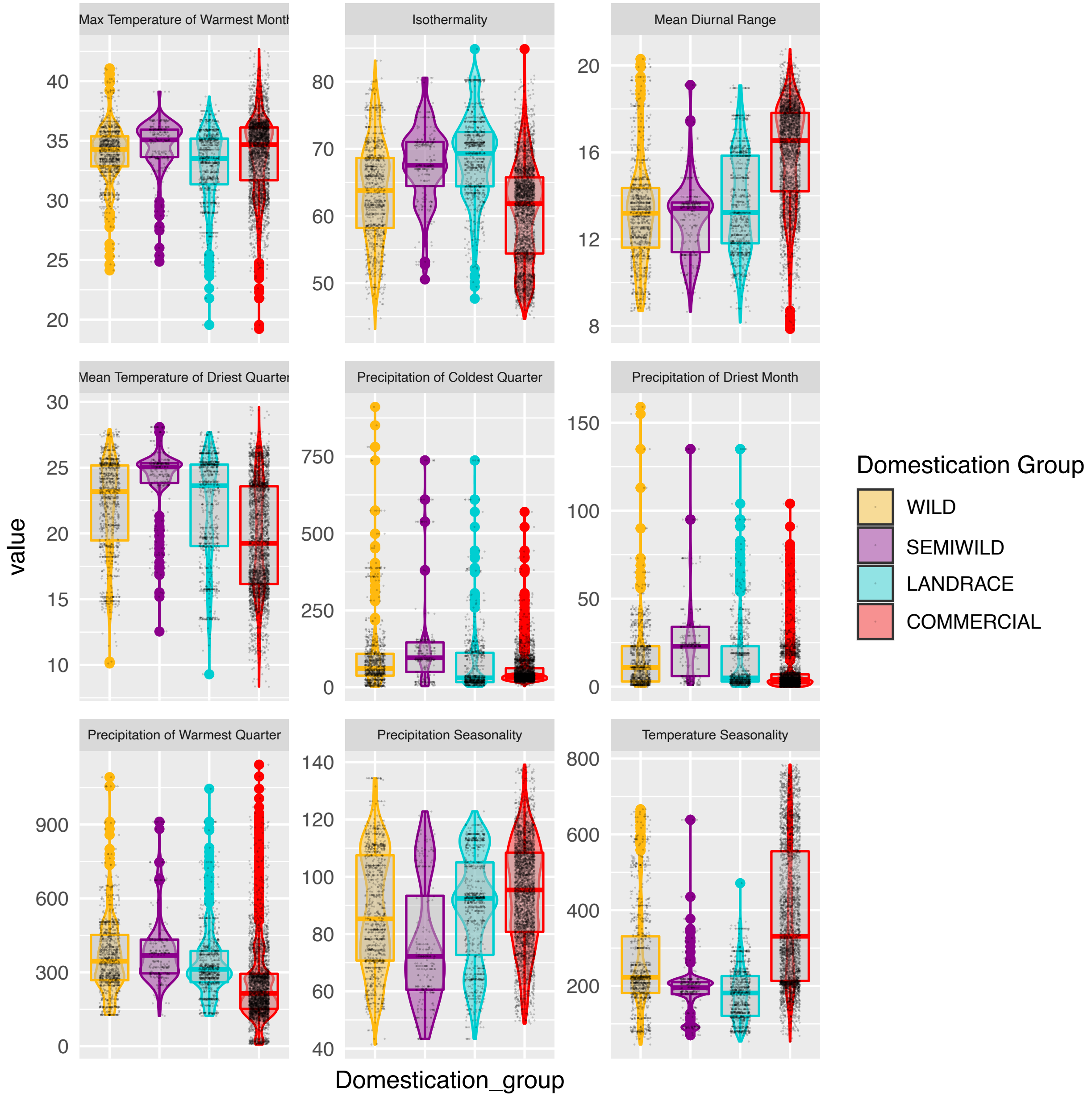

**# SuppFig_5**
A. Median Maxent logistic projection output over 10 replicate runs per domestication class. B. Binary maps with unique thresholds for each domestication class. C. Paired domestication classes binary plots showing overlap area.
##########################
##### Supplementary Figures ###
##########################
**# SuppFig_1**
Maps of Mexico for each domestication category showing the data point occurrences included in this study

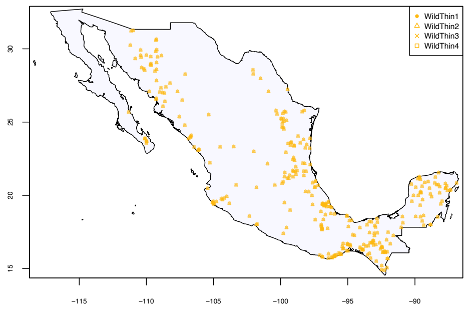
WildThin1
WildThin2
WildThin3
WildThin4
30
25
20
15
−115
−110
−105
−100
−95
−90

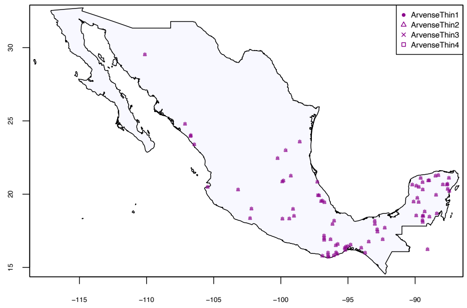
ArvenseThin1
ArvenseThin2
ArvenseThin3
ArvenseThin4
30
25
20
15
−115
−110
−105
−100
−95
−90

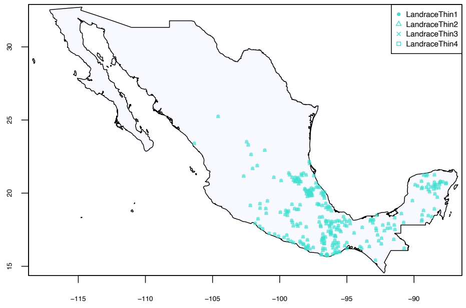
LandraceThin1
LandraceThin2
LandraceThin3
LandraceThin4
30
25
20
15
−115
−110
−105
−100
−95
−90

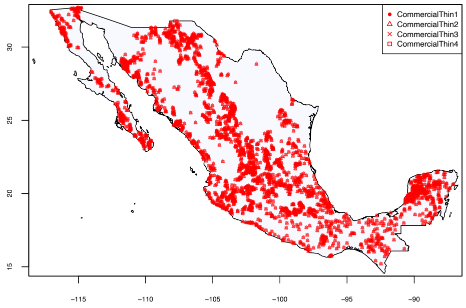
CommercialThin1
CommercialThin2
CommercialThin3
CommercialThin4
30
25
20
15
−115
−110
−105
−100
−95
−90

### SuppFig_2

PCA projection plots on first two principal components, showing pairwise combinations of stepwise domestication classes.

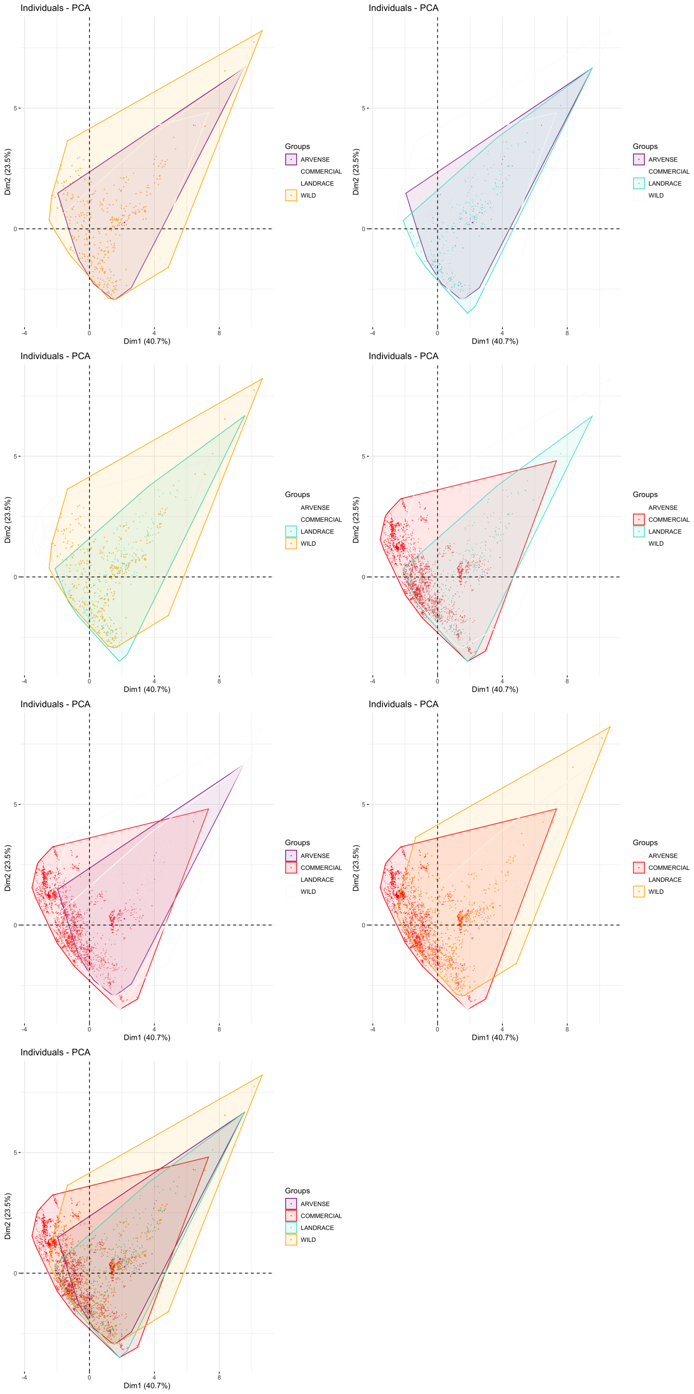

### SuppFig_3

Maps for pairwise combinations of stepwise domestication classes depicting data points exclusive and shared to each class according to the convex hull areas plotted in the corresponding PCAs. Empty circles indicate occurrences found only in one of the classes' hull, filled circles indicate occurrences in the overlapping portion of the hulls.

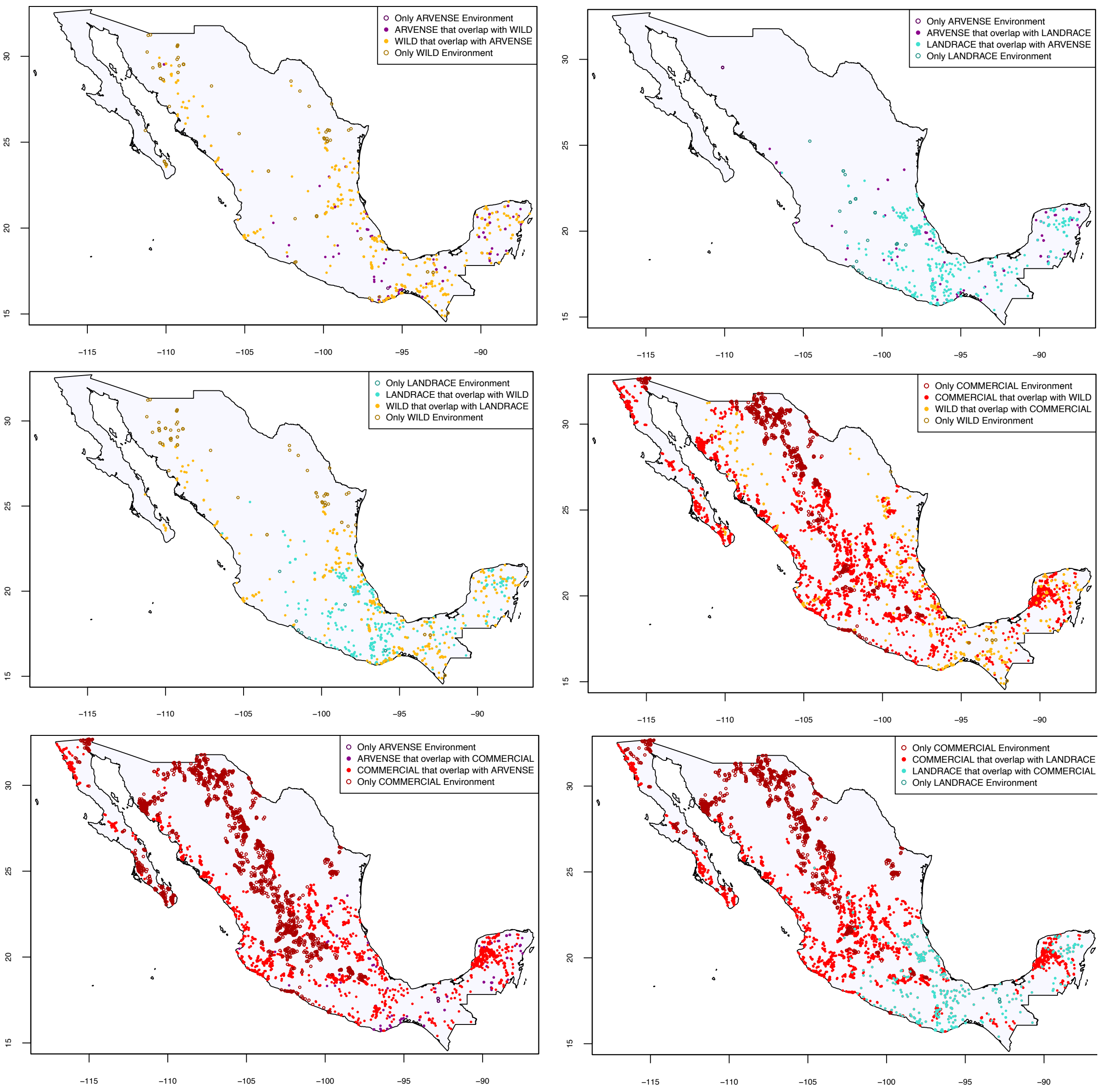

### SuppFig_4
Violin plots of the dispersal for each variable included in PCAs by domestication class.

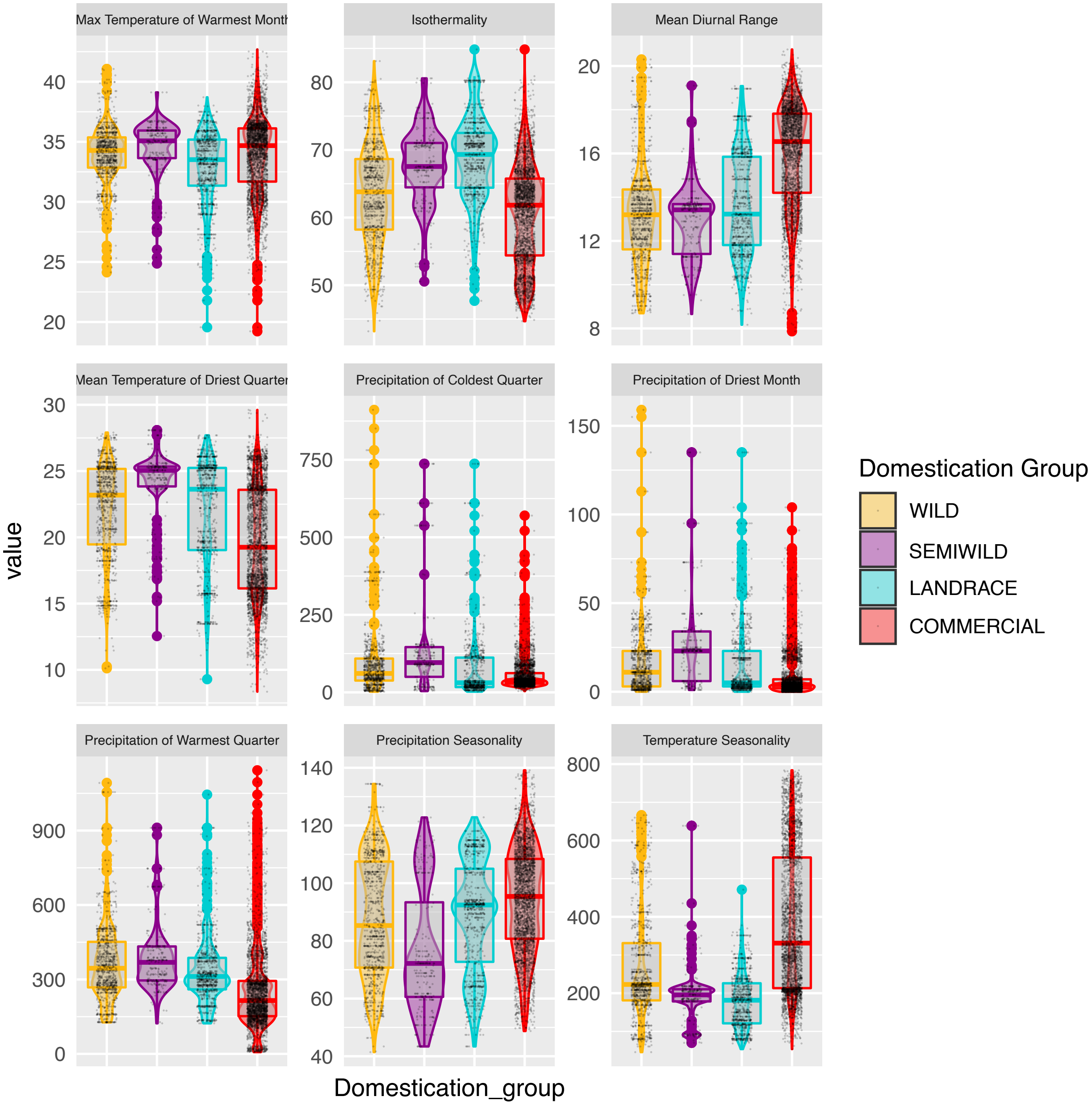

### **SuppFig_5**
A. Median Maxent logistic projection output over 10 replicate runs per domestication class. B. Binary maps with unique thresholds for each domestication class. C. Paired domestication classes binary plots showing overlap area.

A

median

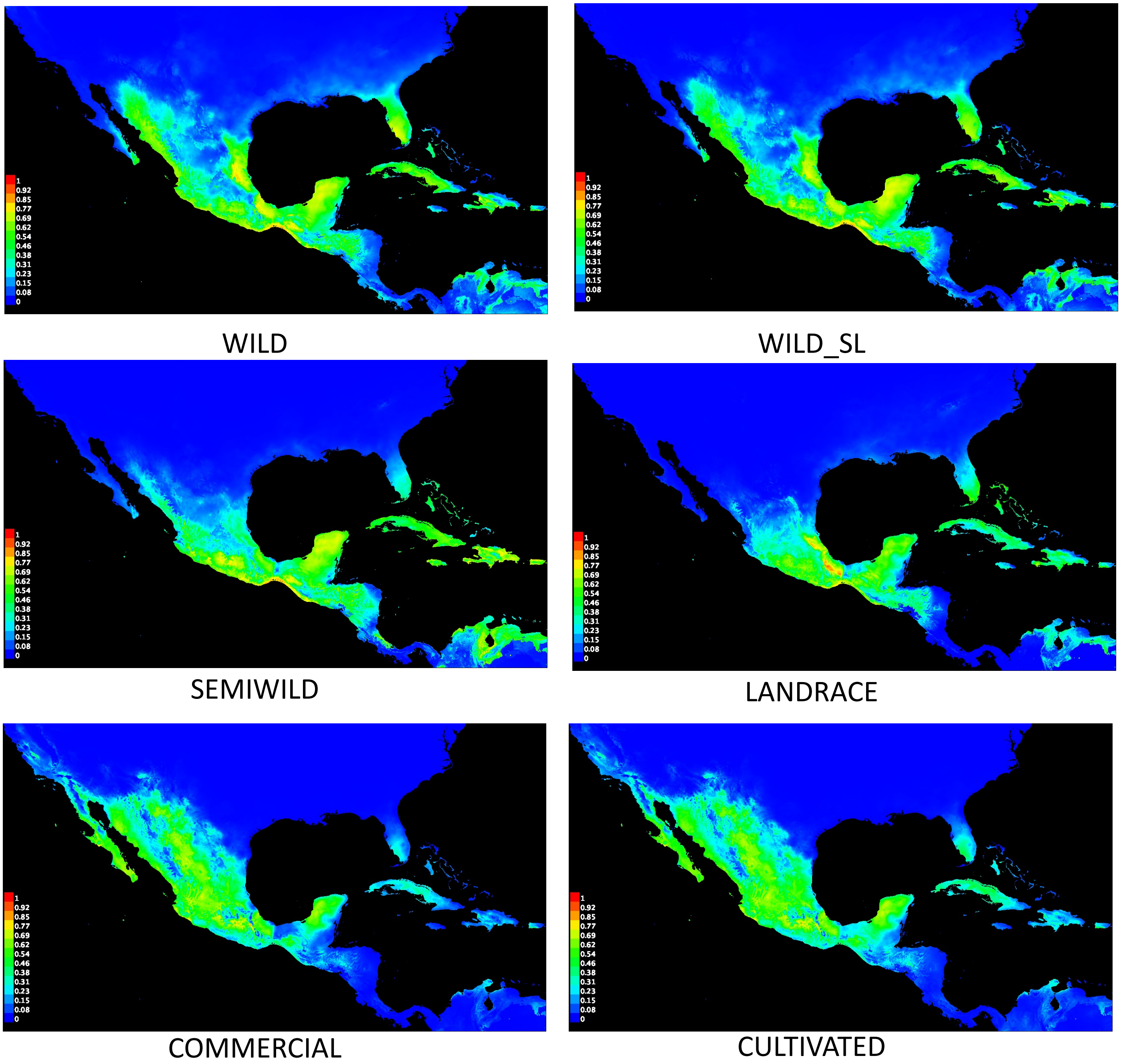

1
0.92
0.85
0.77
0.69
0.62
0.54
0.46
0.38
0.31
0.23
0.15
0.08
0
WILD
WILD_SL
SEMIWILD
LANDRACE
COMMERCIAL
CULTIVATED

B

binary

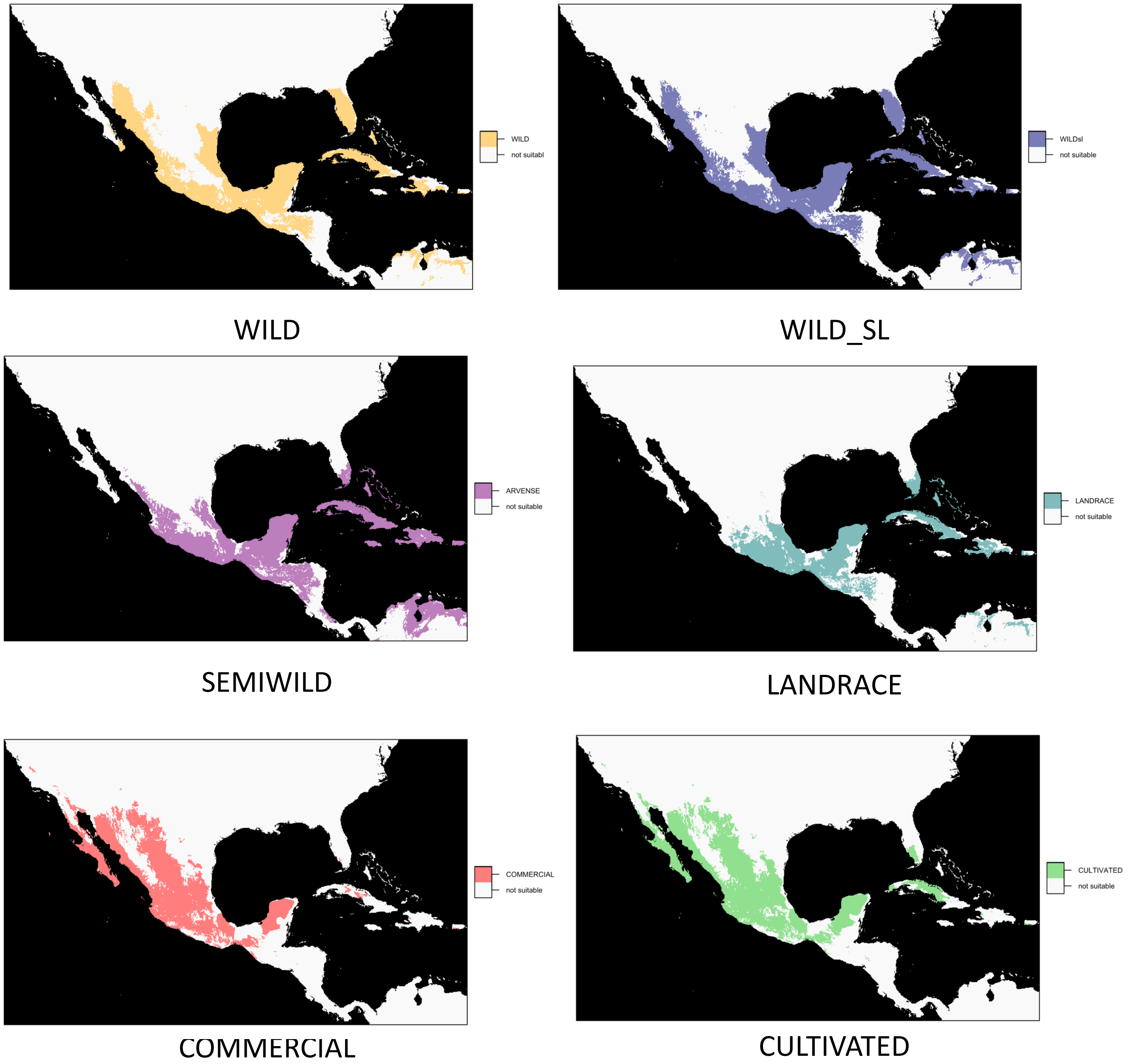
WILD
not suitabl
WILD
WILDsl
not suitable
WILD_SL
ARVENSE
not suitable
SEMIWILD
LANDRACE
not suitable
LANDRACE
COMMERCIAL
not suitable
COMMERCIAL
CULTIVATED
not suitable
CULTIVATED

# C

binary paired

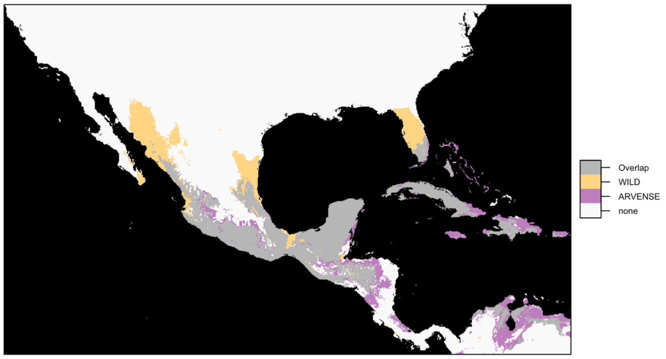


WILD-SEMIWILD

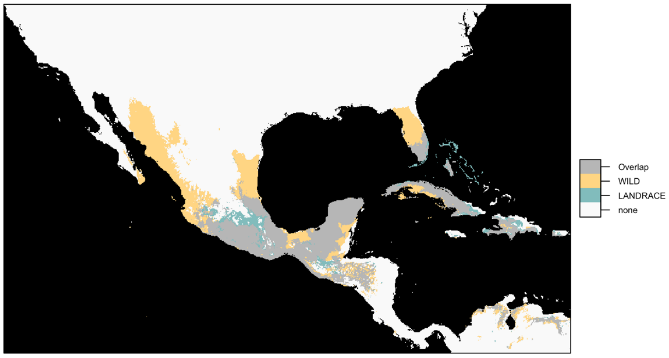


WILD-LANDRACE

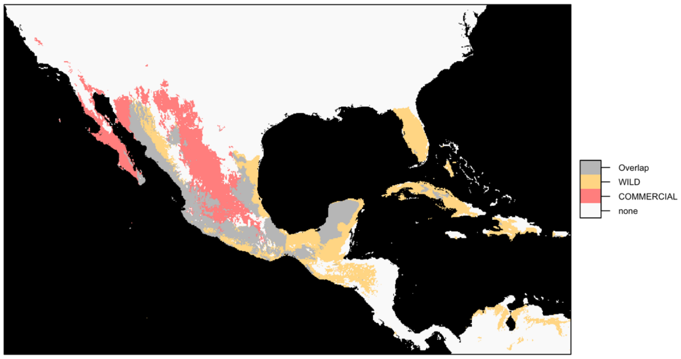


WILD-COMMERCIAL

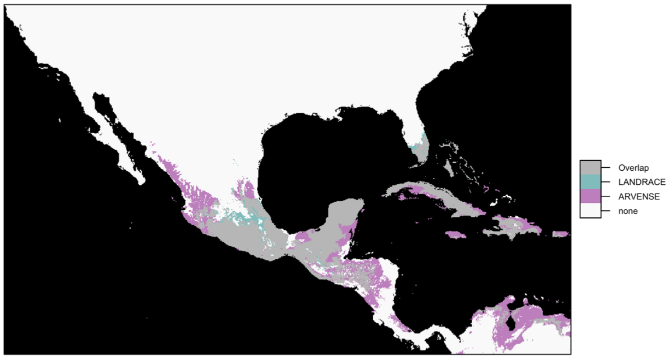


SEMIWILD-LANDRACE

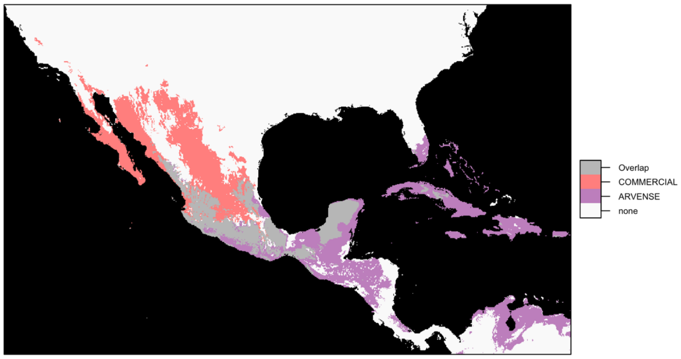


SEMIWILD-COMMERCIAL

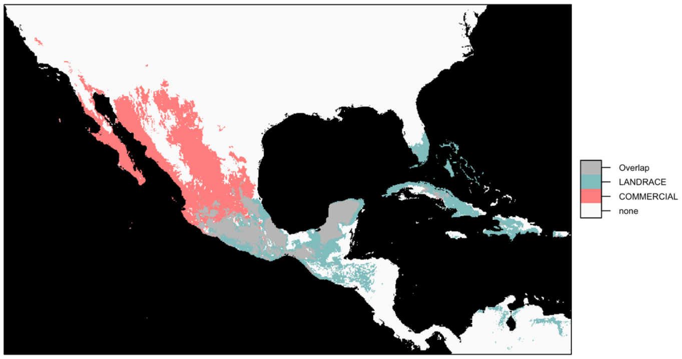


LANDRACE-COMMERICAL

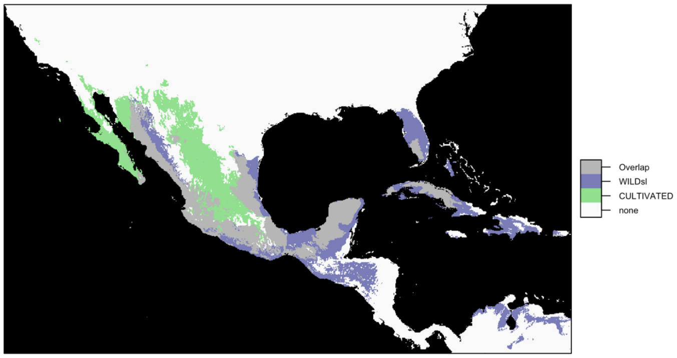


WILD_SL-CULTIVATED

### SuppFig_6

Stacked bar plots showing permutation importance of each variable included in the maxent niche modeling per domestication class.

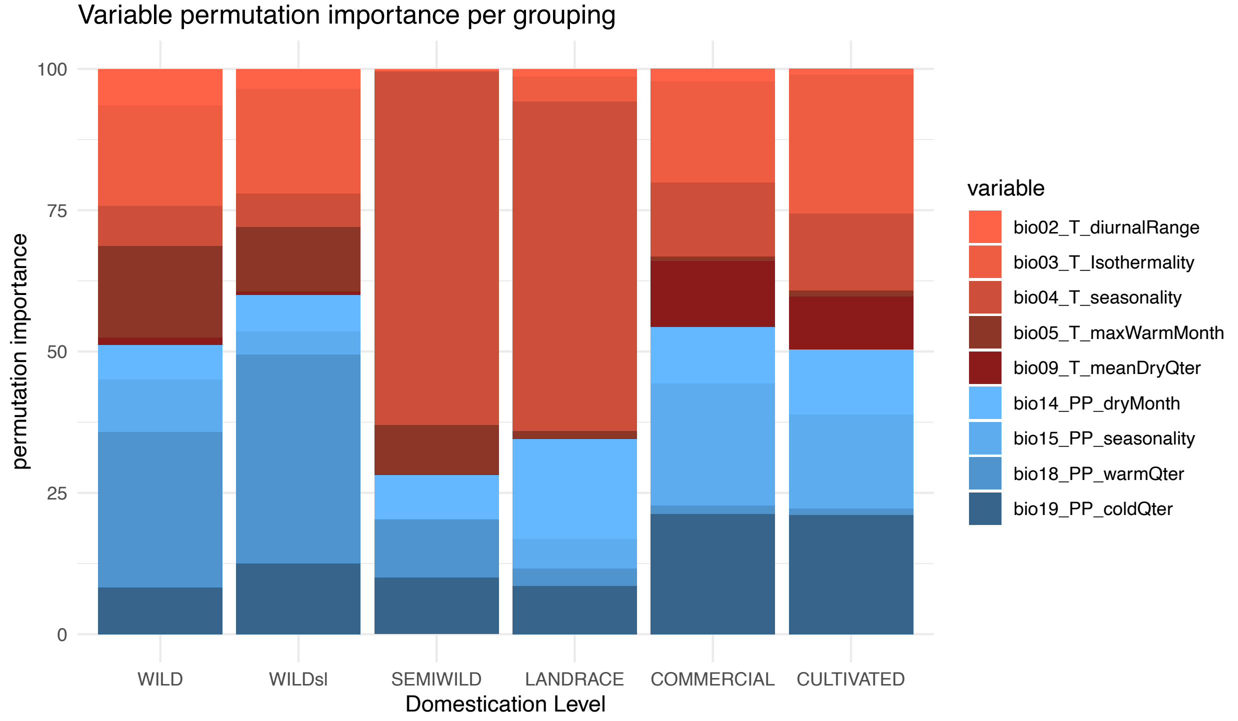

### SuppFig_7

Boxplots depicting pixels lost (A), kept (B) and new (C) for each domestication class, according to area differences between present maxent niche model geographical projections and their future projections under three years and two SSP pathways associated to rcp 45 and 85.

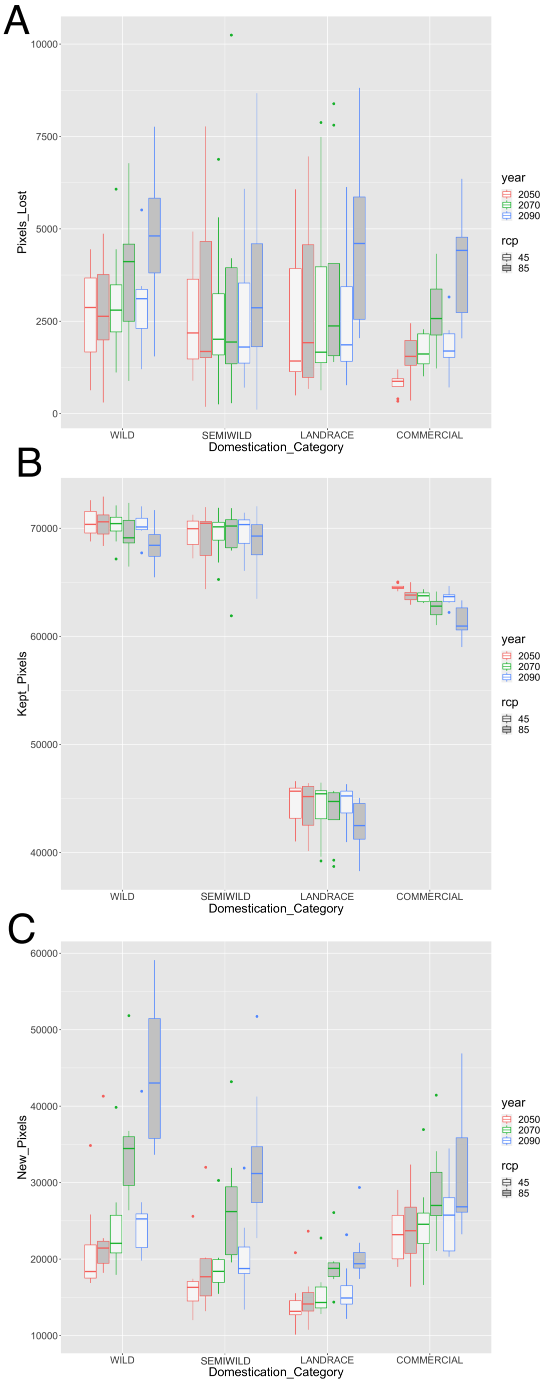

### SuppFig_8

Background test histograms comparing niche similarity values for D and I in comparison pairs of wilds vs commercials and wilds vs landraces.

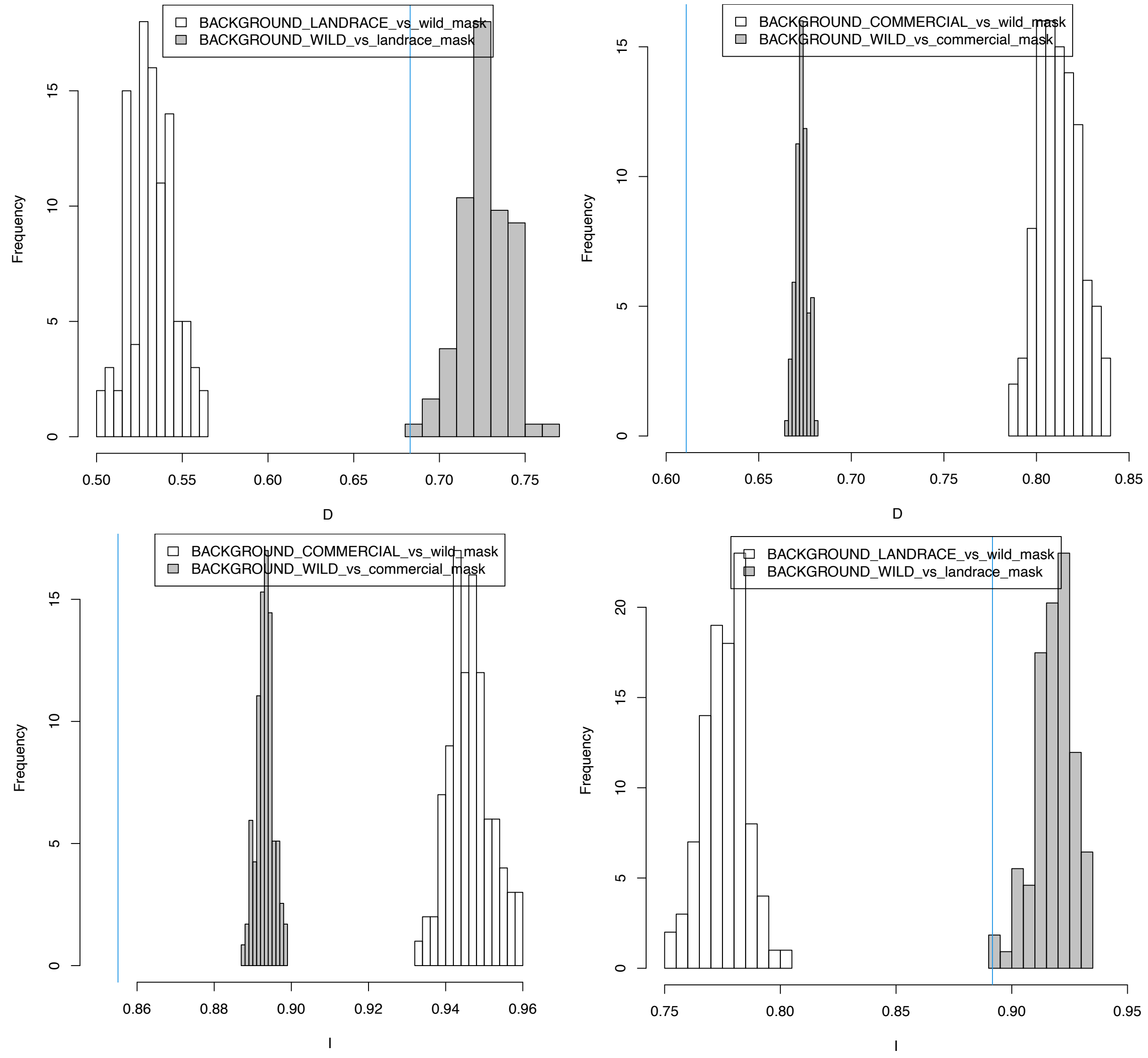

**# SuppFig_9**
Multivariate Environmental Similarity Surfaces for each year-SSP-GCM combinations.

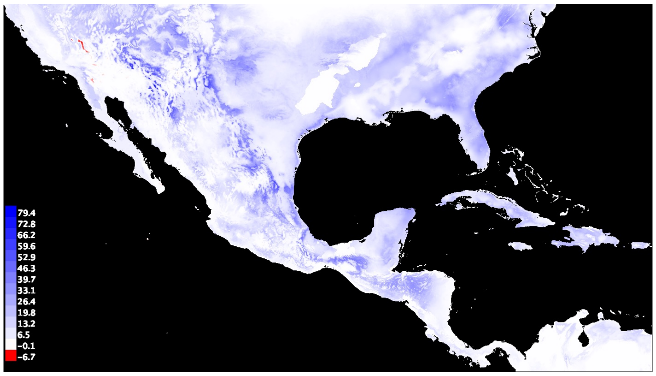


BCC-CSM2-MR

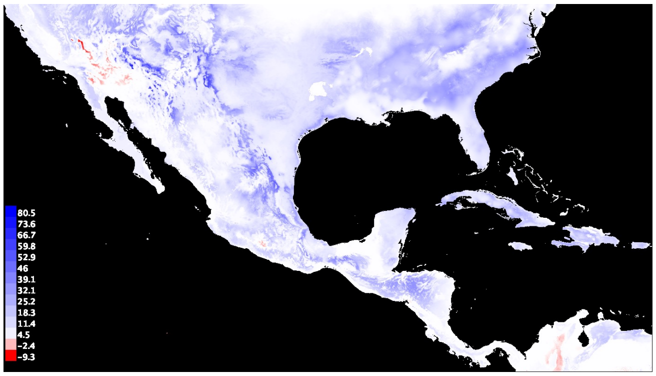


CanESM5

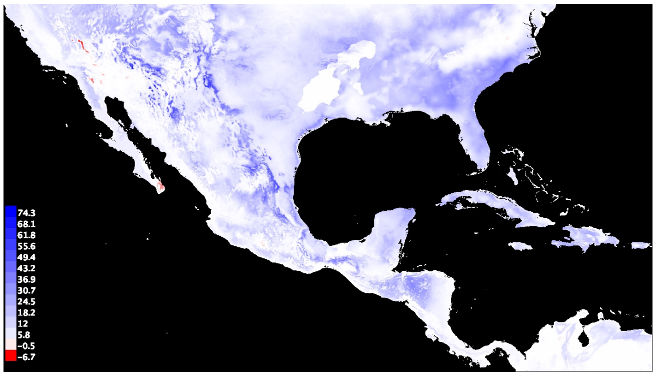


CNRM-CM6-1

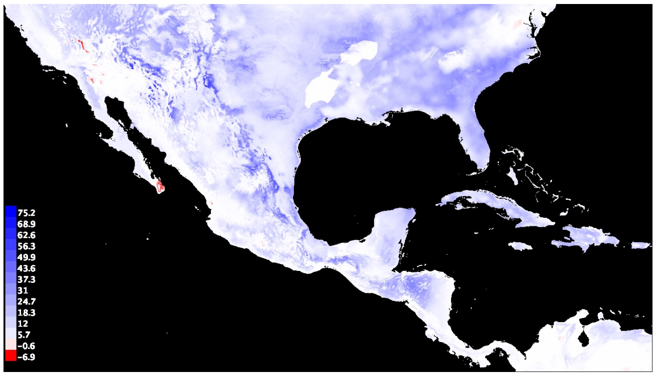


CNRM-ESM2-1

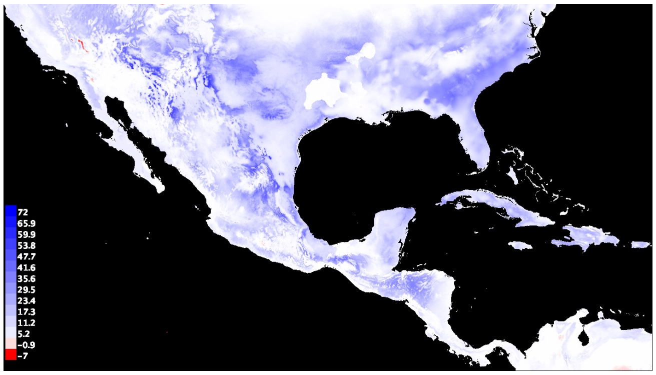


IPSL-CM6A-LR

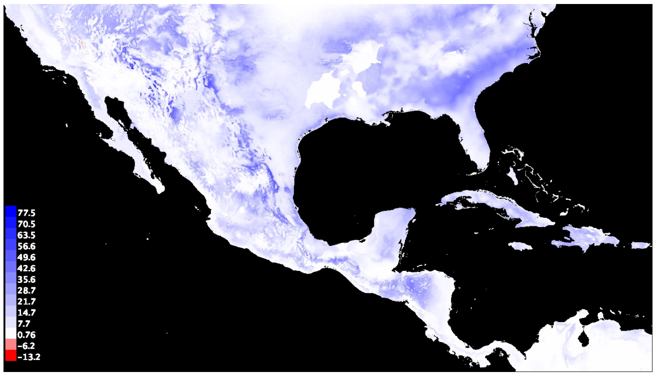


MIROC-ES2L

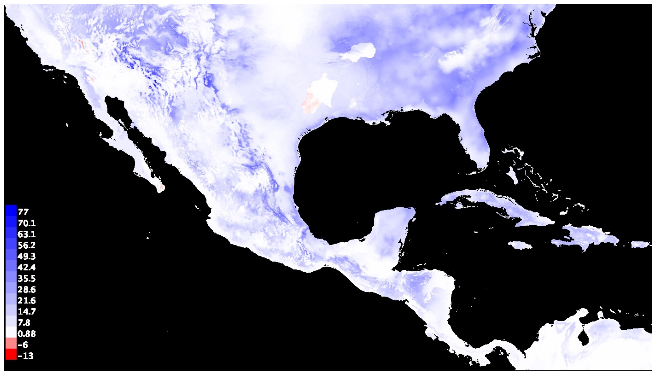


MIROC6

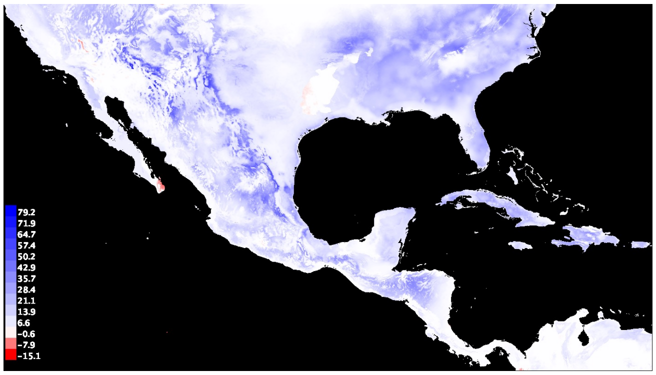


MRI-ESM2-0

**2050** SSP=2_45

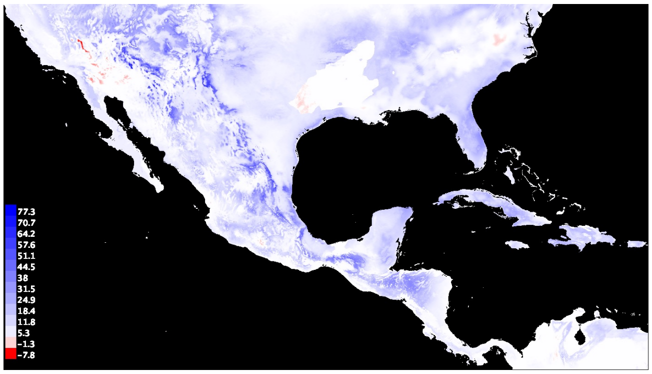


BCC-CSM2-MR

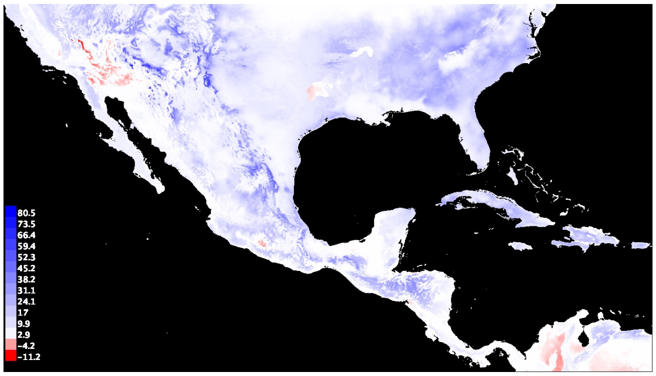


CanESM5

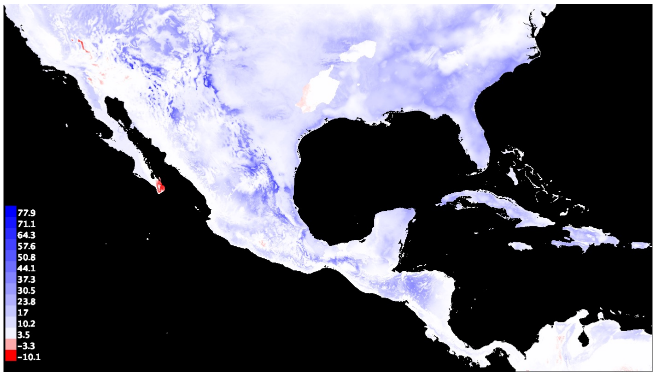


CNRM-CM6-1

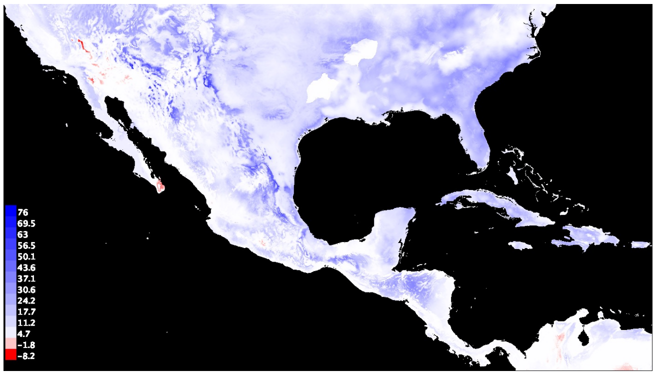


CNRM-ESM2-1

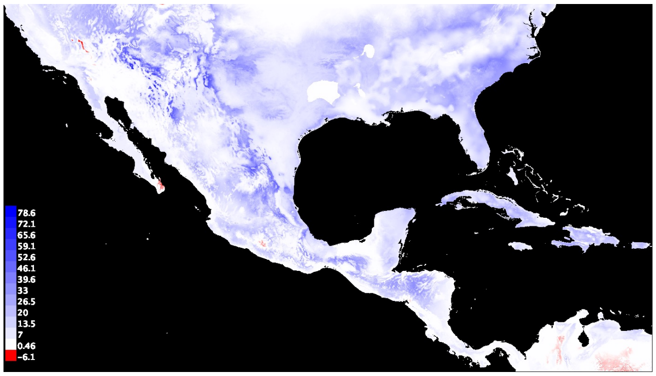


IPSL-CM6A-LR

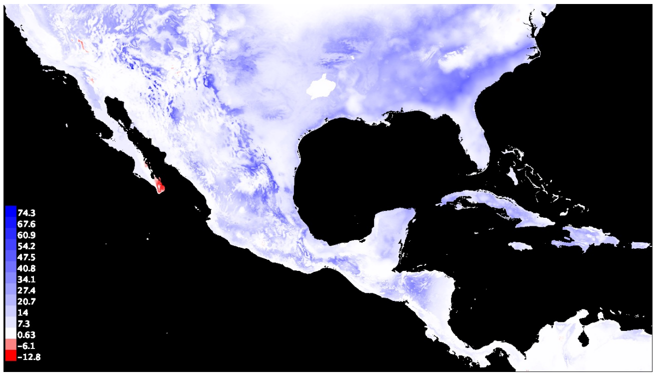


MIROC-ES2L

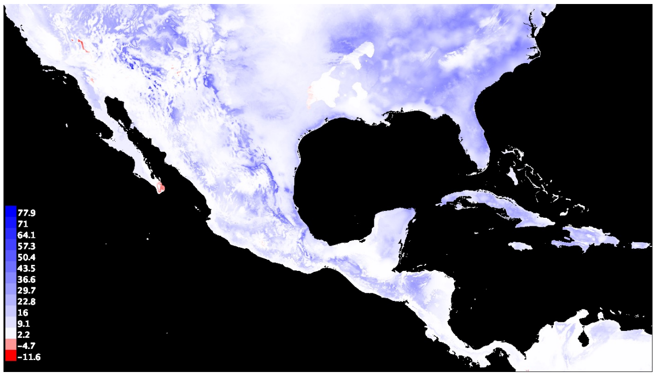


MIROC6

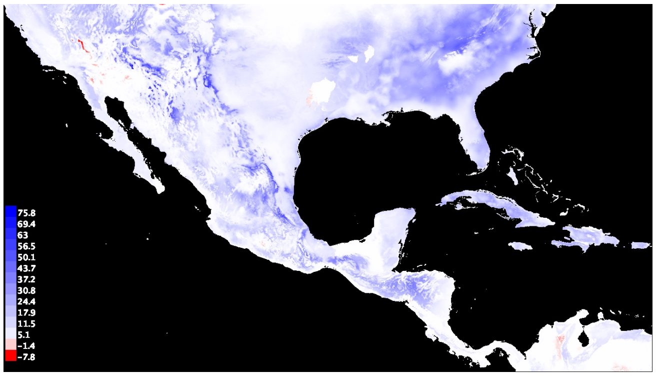


MRI-ESM2-0

**2050** SSP=5_85

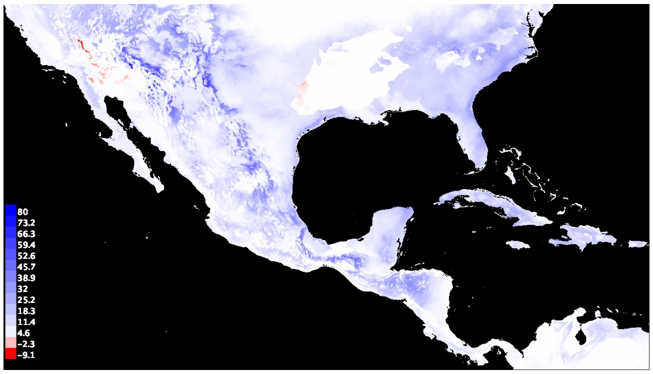


BCC-CSM2-MR

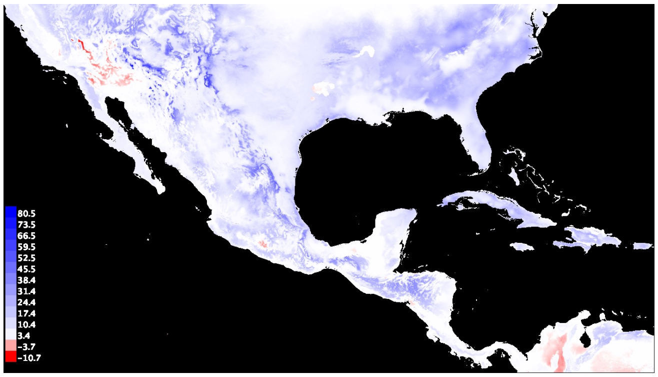


CanESM5

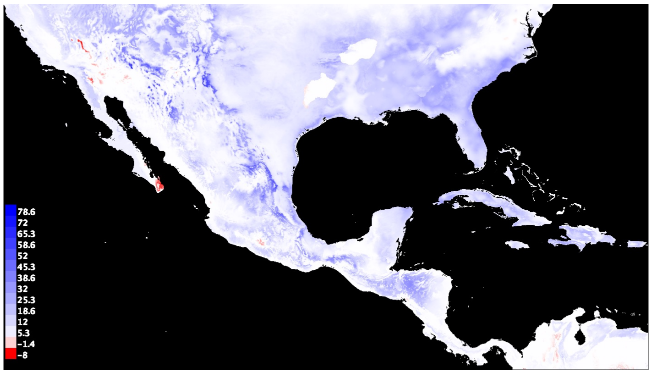


CNRM-CM6-1

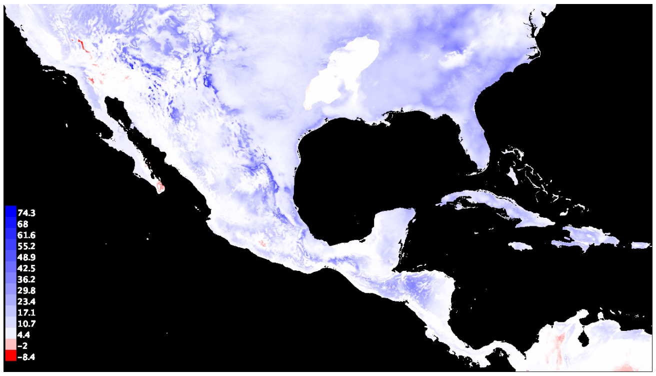


CNRM-ESM2-1

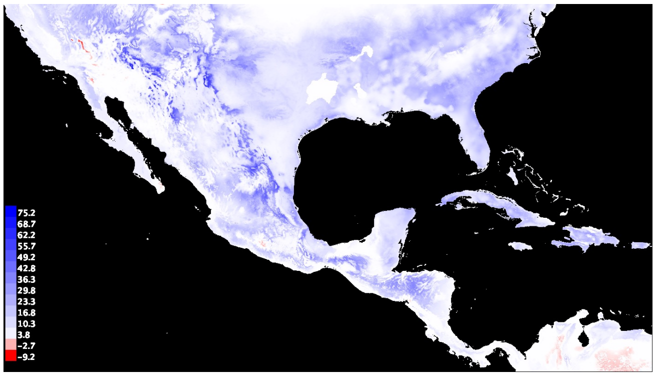


IPSL-CM6A-LR

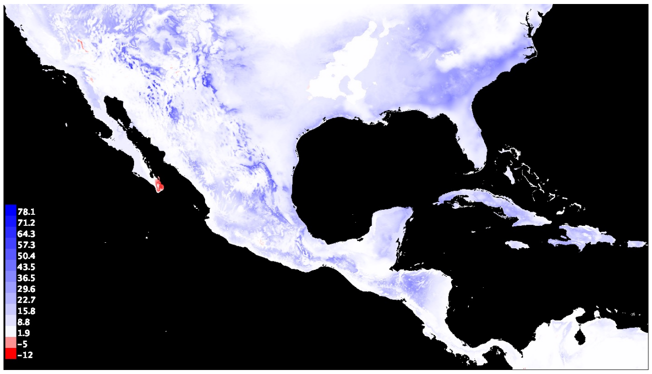


MIROC-ES2L

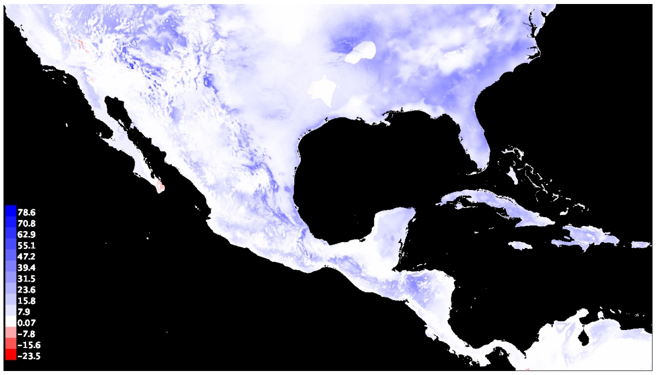


MIROC6

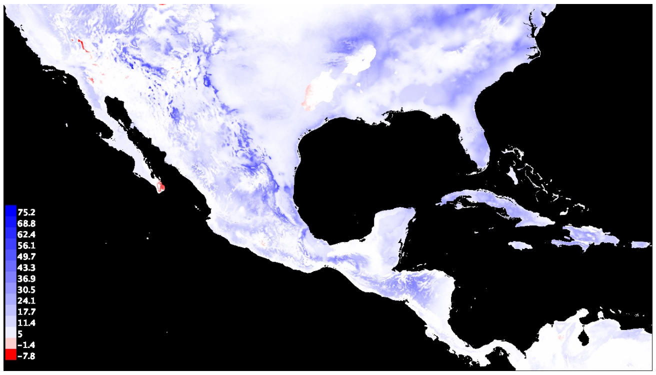


MRI-ESM2-0

**2070** SSP=2_45

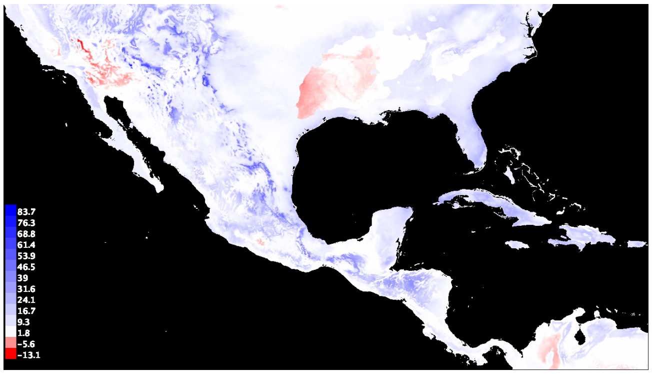


BCC-CSM2-MR

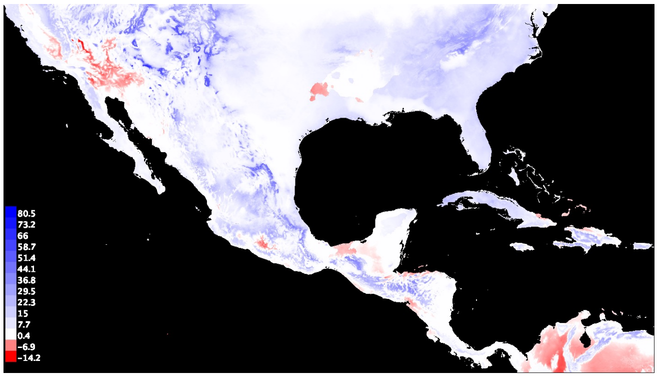


CanESM5

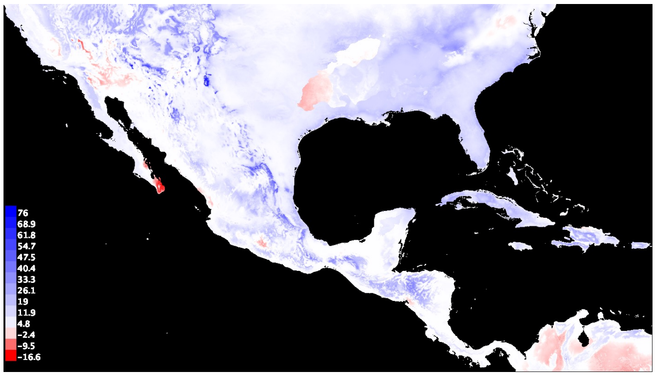


CNRM-CM6-1

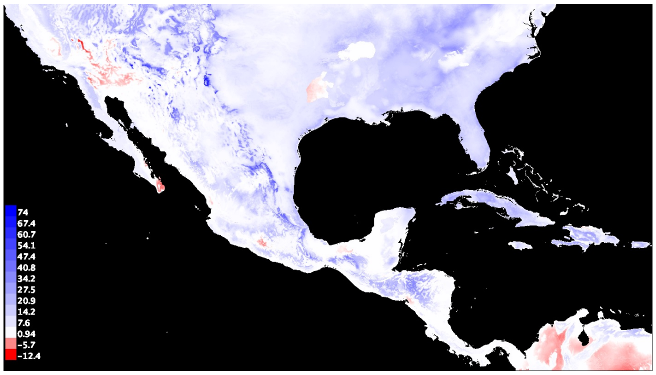


CNRM-ESM2-1

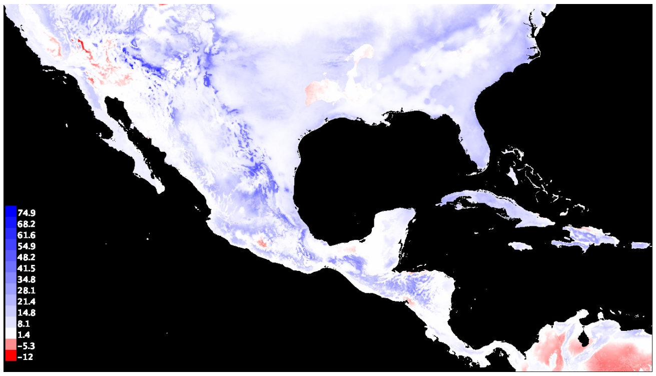


IPSL-CM6A-LR

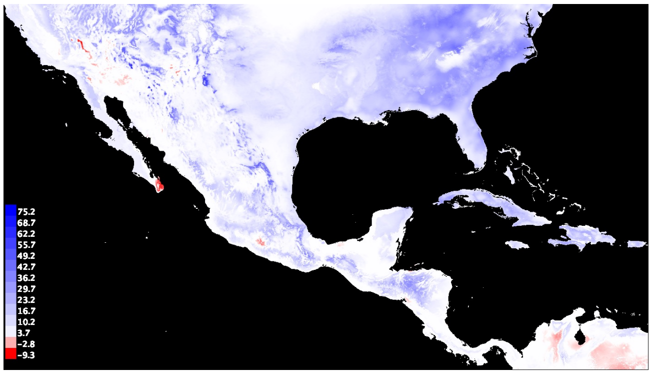


MIROC-ES2L

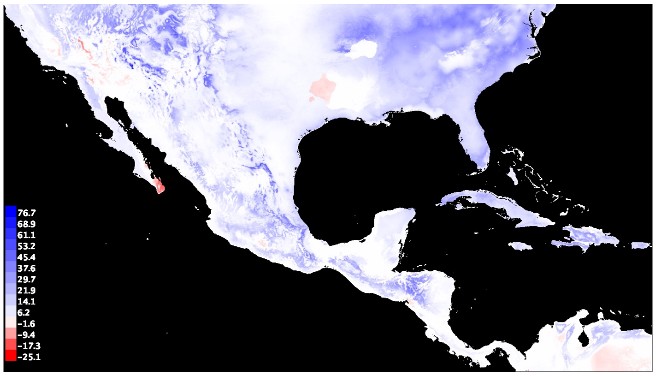


MIROC6

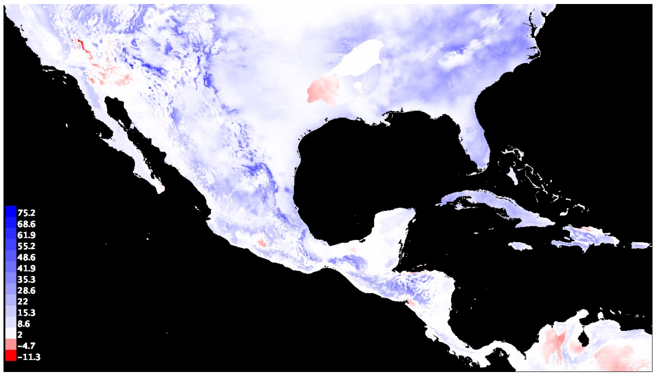


MRI-ESM2-0

**2070** SSP=5_85

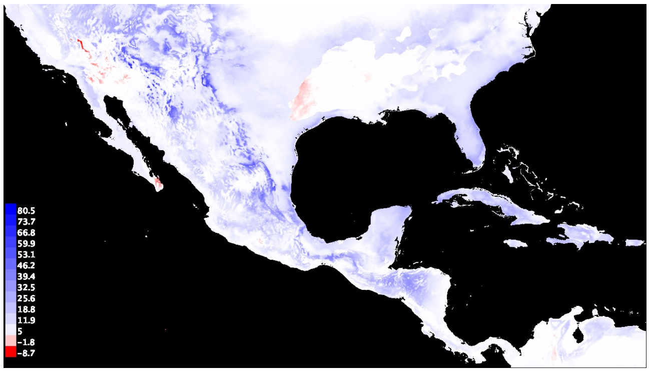


BCC-CSM2-MR

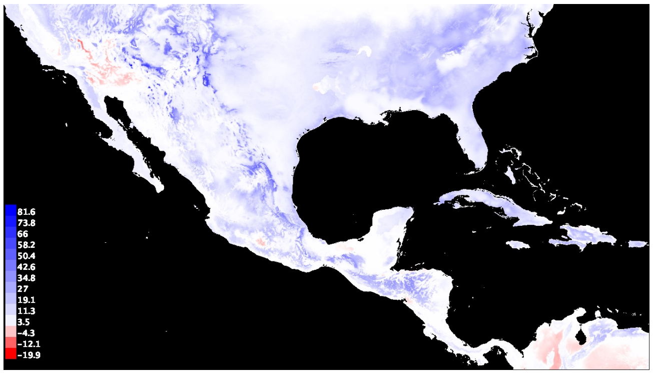


CanESM5

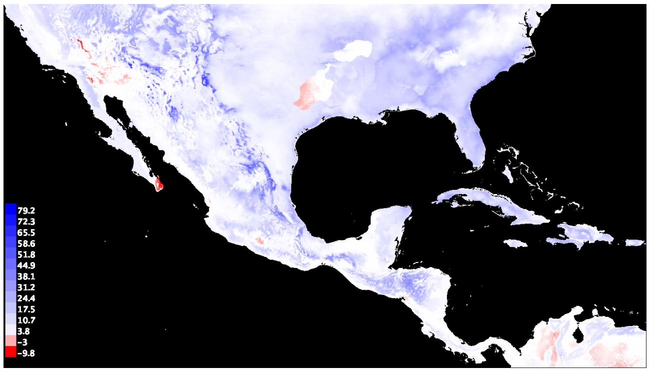


CNRM-CM6-1

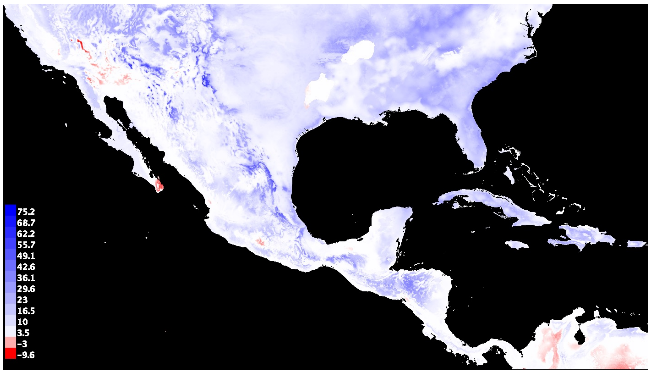


CNRM-ESM2-1

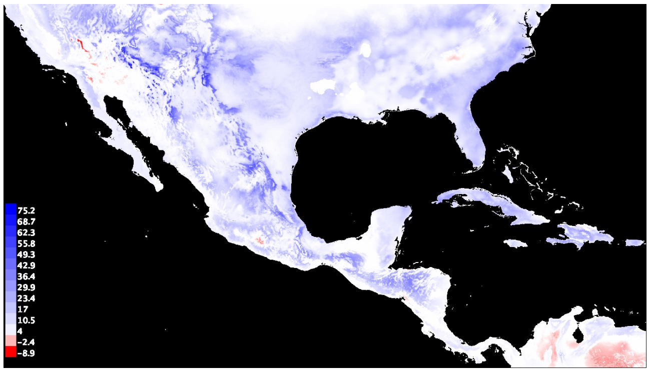


IPSL-CM6A-LR

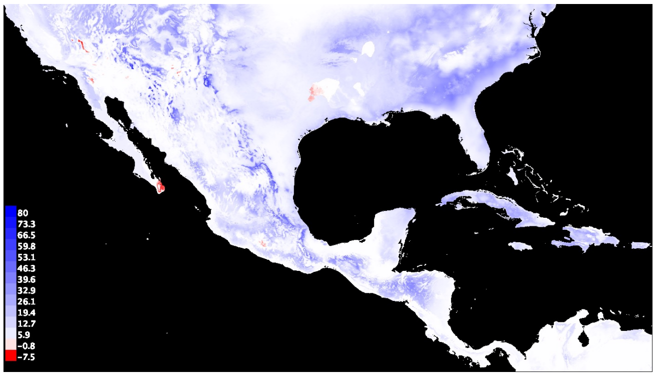


MIROC-ES2L

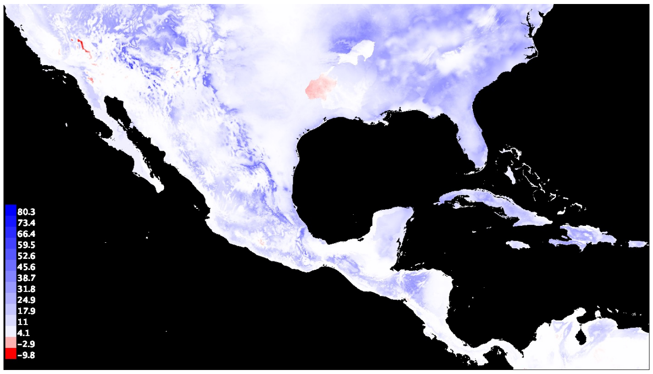


MIROC6

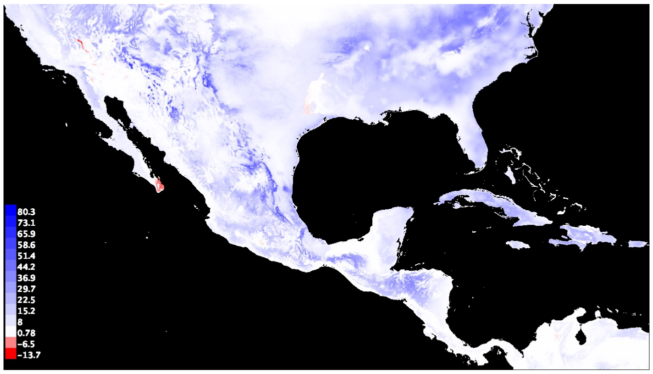


MRI-ESM2-0

**2090** SSP=2_45

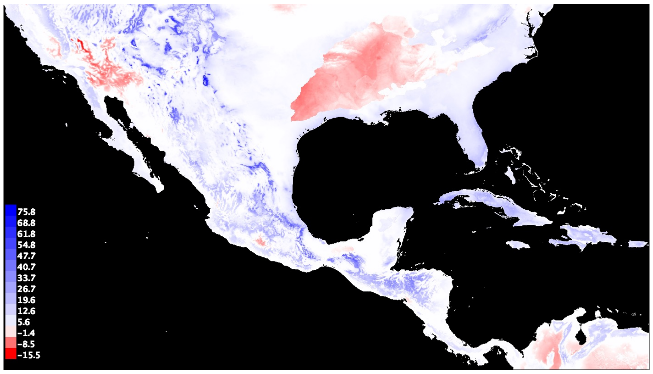


BCC-CSM2-MR

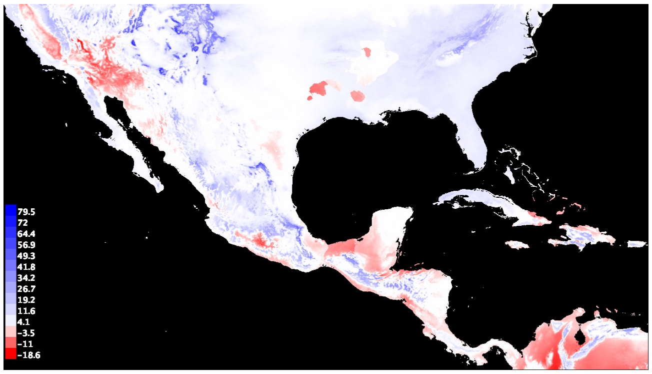


CanESM5

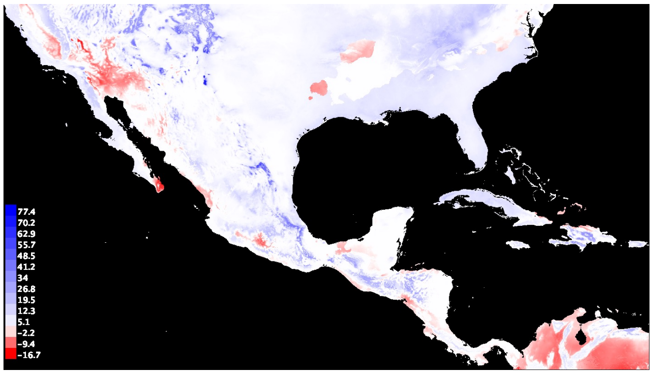


CNRM-CM6-1

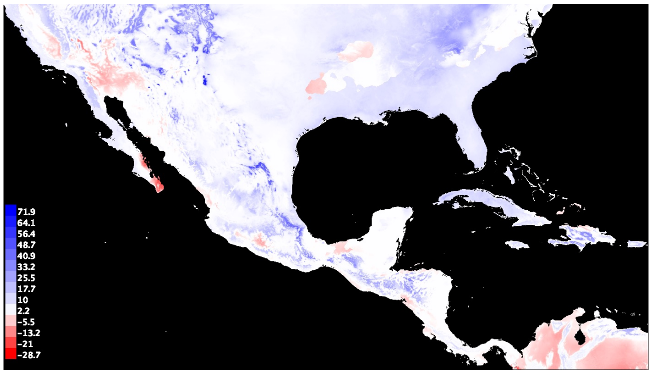


CNRM-ESM2-1

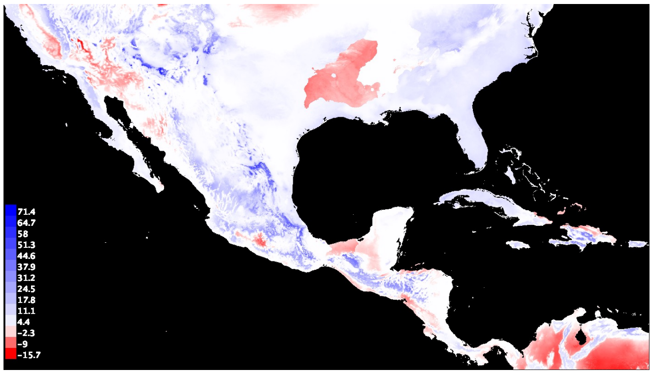


IPSL-CM6A-LR

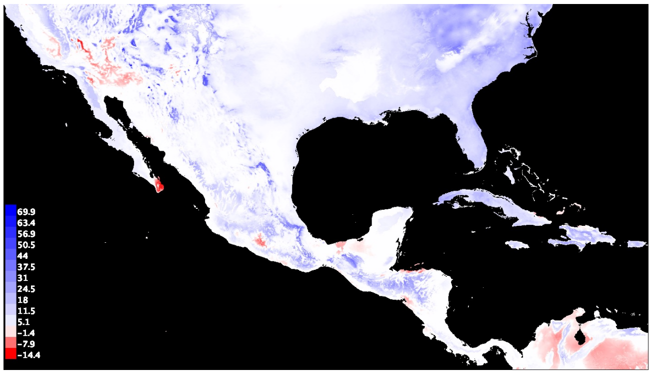


MIROC-ES2L

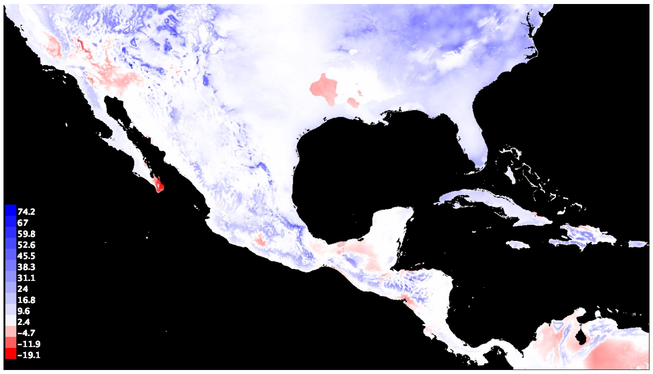


MIROC6

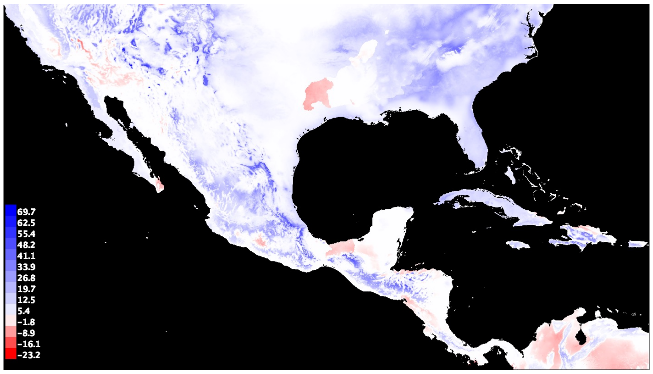


MRI-ESM2-0

**2090** SSP=5_85

### # SuppFig_10

Multivariate Environmental Similarity Surfaces found to diverge in landraces.

LANDRACES

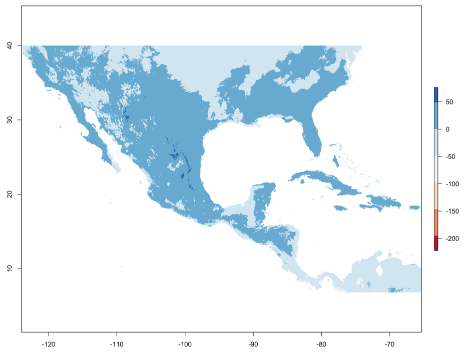


MRI-ESM2-0

**2090** SSP=2_45

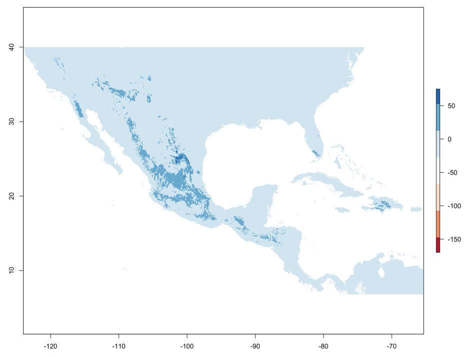


IPSL-CM6A-LR

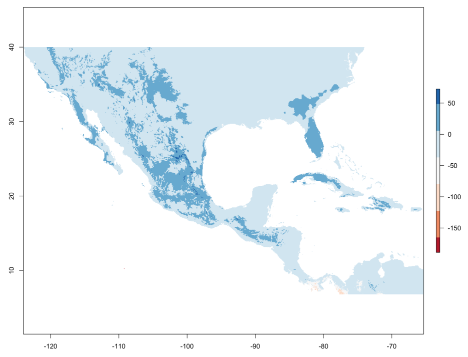


MRI-ESM2-0

**2090** SSP=5_85

### Supp Fig_A1

Factor map for PCA analyses of all samples, soil and climate variables marking associated uncertainty per variable due to missing values imputation.

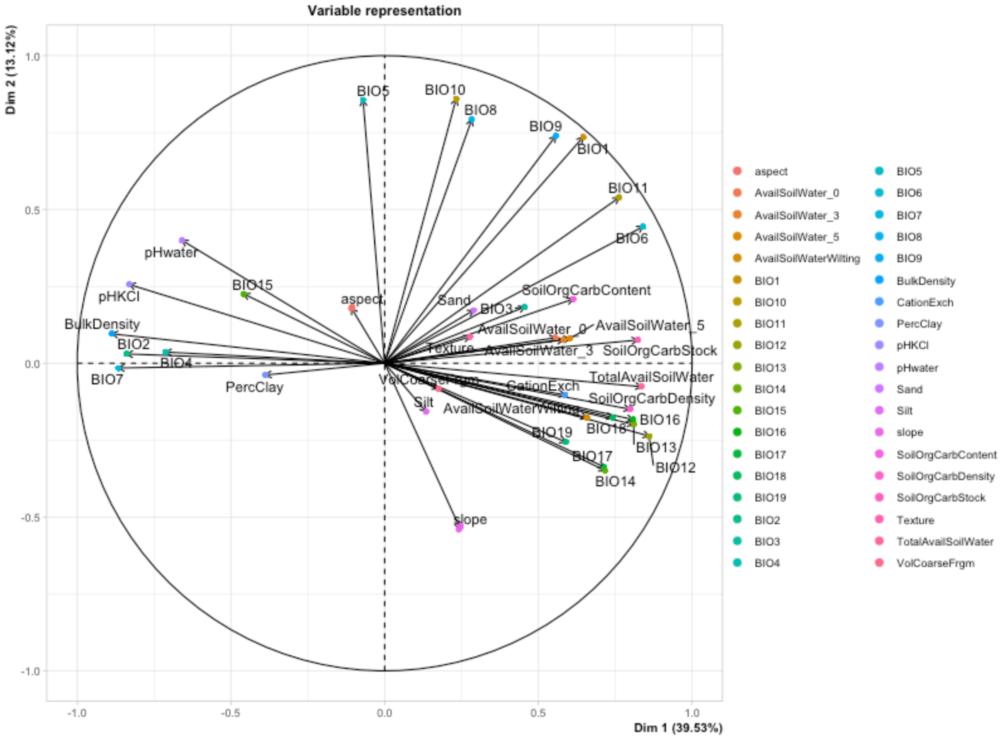

### **SuppFig_A2**

Correlation plots for climatic variables calculated for the whole dataset, broad domestication categories (wild *sl* and cultivated) and stepwise domestication categories (Wild, Arvense, Landrace and Commercial).

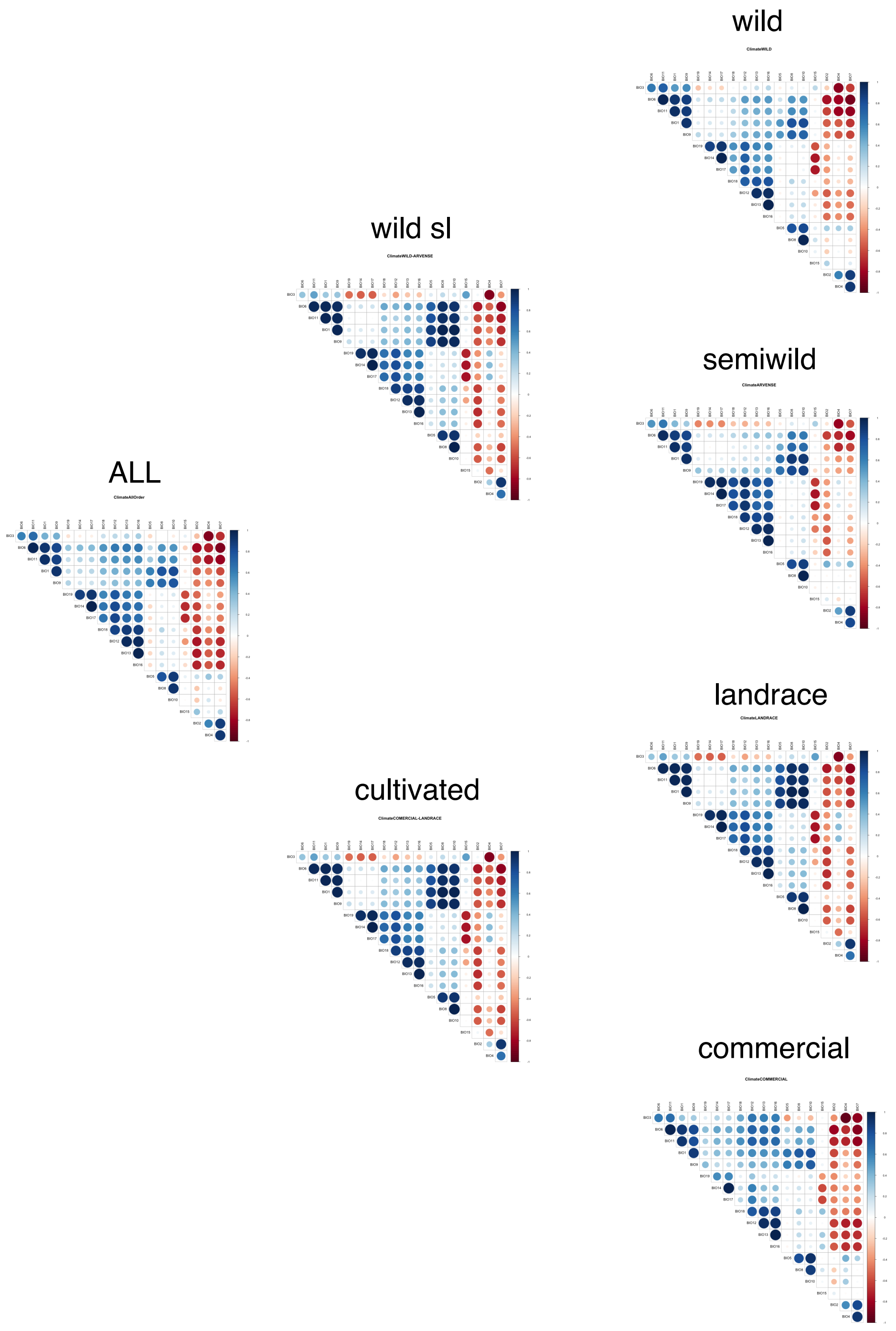

### # SuppFig_A3

Effect of varying combinations of feature class and regularization multiplier values on average test AUC, AICc and OR for maxent modeling tunning

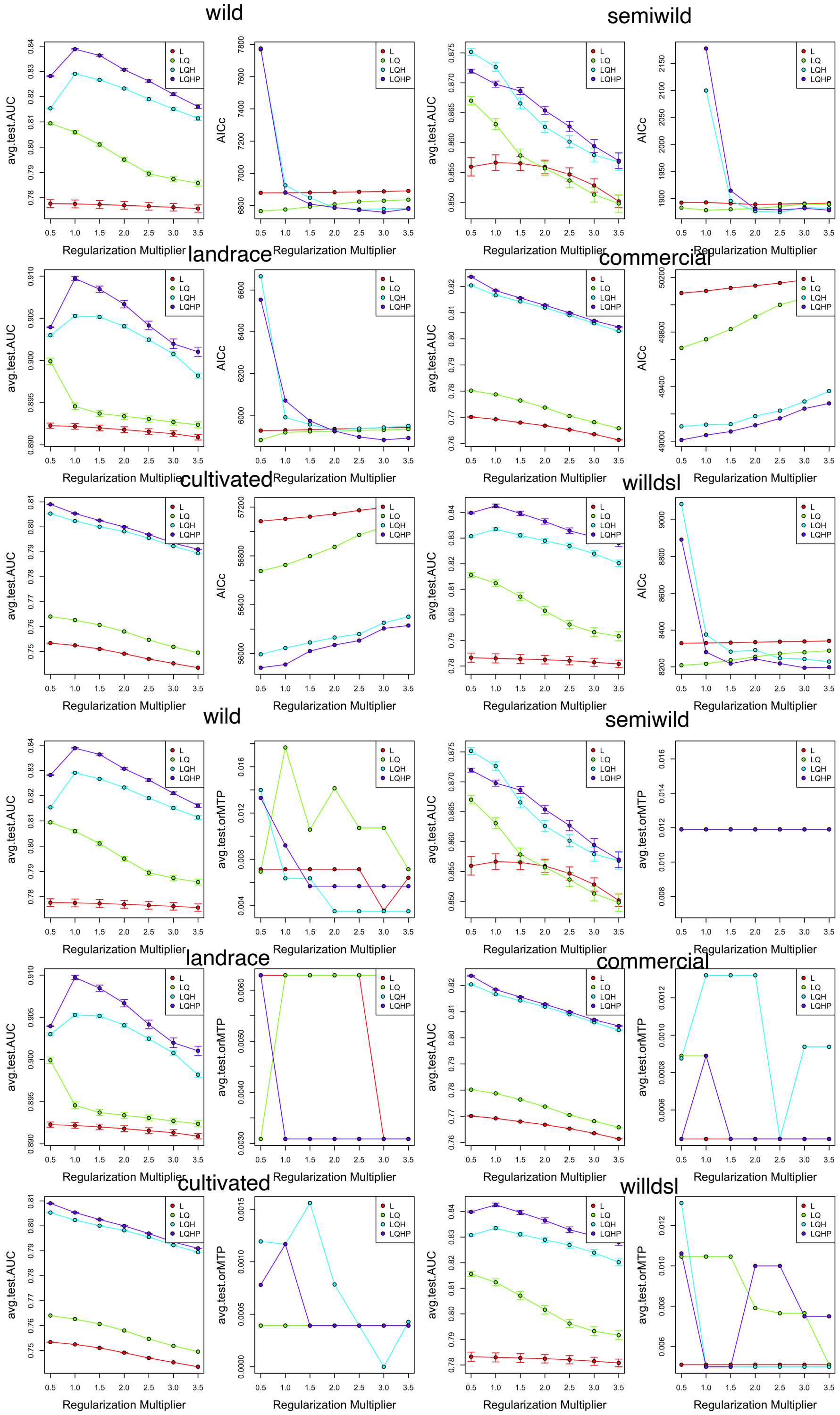

### SuppFigs_A4
Future projections of niche modeling per domestication class. For each class and year-ssp combination, all GCMs are plotted showing gradual shading when overlap occurs.

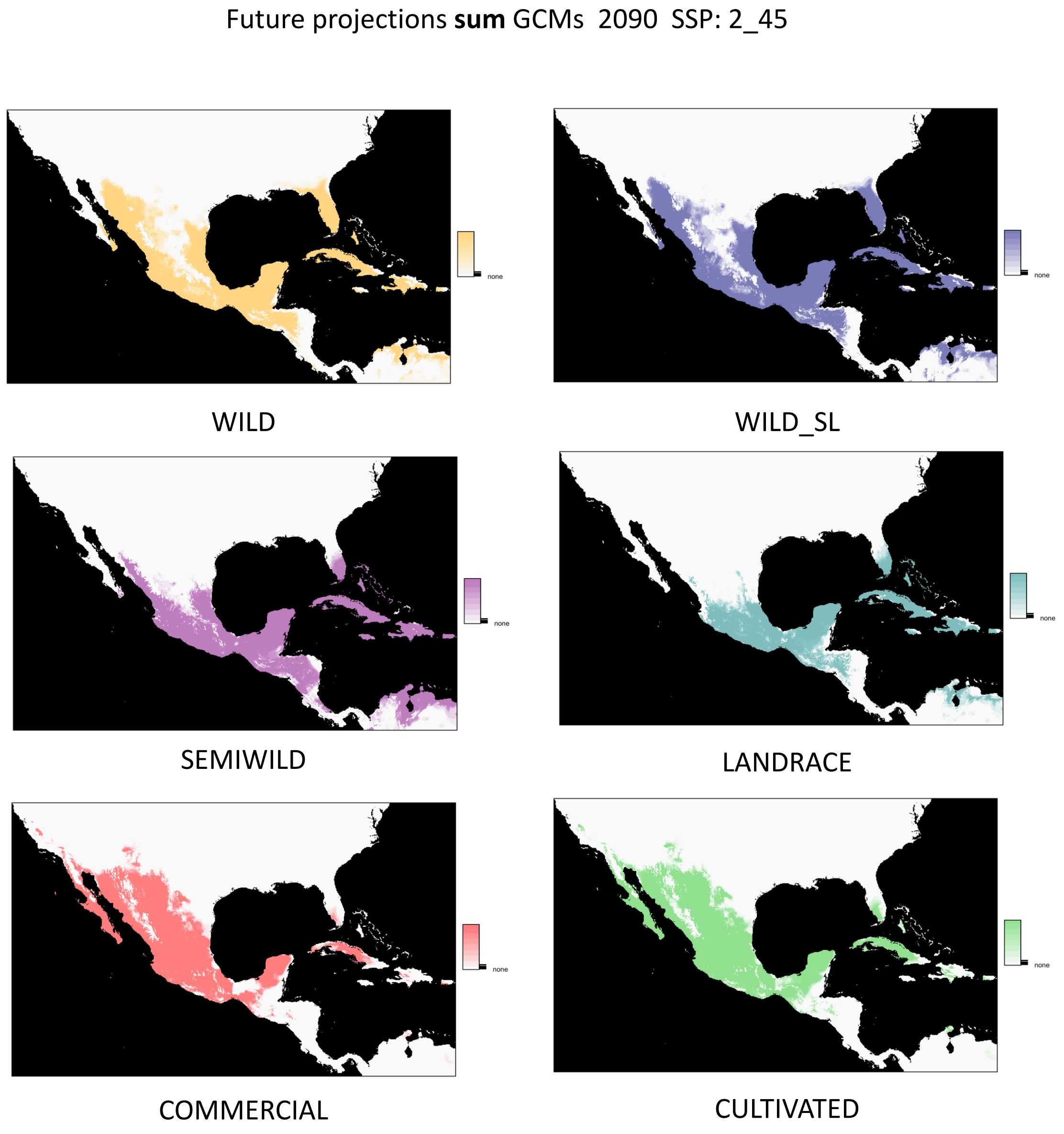

### **SuppFig_A5**
Future projections of niche modeling per domestication class displaying the overlap of each class with respect to its present-day projection.

### **SuppFig_A6**
Future projections of niche modeling per domestication class displaying the overlap among pairwise classes.
