## Supplementary_Tables for "Capturing the distribution as it shifts: chile pepper (*Capsicum annuum* L.) domestication gradient meets geography"

### Supplementary table 2

**Variables Contribution**

| | Dim.1 | Dim.2 | Dim.3 | Dim.4 | Dim.5 |
|---|---|---|---|---|---|
| BIO2 | 18.8 | 0.8 | 0.1 | 0.6 | 65.8 |
| BIO3 | 7.4 | 26.8 | 0.3 | 8.8 | 16.2 |
| BIO4 | 13.6 | 19.1 | 2.1 | 3.8 | 0.6 |
| BIO5 | 0.0 | 5.0 | 53.7 | 2.7 | 2.8 |
| BIO9 | 10.9 | 0.8 | 33.6 | 3.4 | 0.6 |
| BIO14 | 16.3 | 13.8 | 2.6 | 0.0 | 2.4 |
| BIO15 | 4.8 | 16.8 | 6.7 | 36.8 | 0.0 |
| BIO18 | 14.8 | 0.2 | 0.3 | 41.8 | 6.8 |
| BIO19 | 13.3 | 16.7 | 0.6 | 2.1 | 4.8 |

**Variables Coordinates**

| | Dim.1 | Dim.2 | Dim.3 | Dim.4 | Dim.5 |
|---|---|---|---|---|---|
| BIO2 | -0.83 | 0.13 | -0.03 | -0.07 | 0.53 |
| BIO3 | 0.52 | -0.75 | -0.06 | -0.27 | 0.26 |
| BIO4 | -0.71 | 0.63 | 0.18 | 0.17 | -0.05 |
| BIO5 | -0.02 | 0.32 | 0.91 | -0.15 | 0.11 |
| BIO9 | 0.63 | -0.13 | 0.72 | -0.17 | -0.05 |
| BIO14 | 0.77 | 0.54 | -0.20 | 0.02 | 0.10 |
| BIO15 | -0.42 | -0.60 | 0.32 | 0.54 | 0.01 |
| BIO18 | 0.74 | -0.07 | 0.07 | 0.58 | 0.17 |
| BIO19 | 0.70 | 0.59 | -0.10 | 0.13 | 0.14 |

**Variables Loadings**

| | Dim.1 | Dim.2 | Dim.3 | Dim.4 | Dim.5 |
|---|---|---|---|---|---|
| BIO2 | -0.43 | 0.09 | -0.03 | -0.08 | 0.81 |
| BIO3 | 0.27 | -0.52 | -0.05 | -0.30 | 0.40 |
| BIO4 | -0.37 | 0.44 | 0.15 | 0.19 | -0.08 |
| BIO5 | -0.01 | 0.22 | 0.73 | -0.16 | 0.17 |
| BIO9 | 0.33 | -0.09 | 0.58 | -0.18 | -0.08 |
| BIO14 | 0.40 | 0.37 | -0.16 | 0.02 | 0.15 |
| BIO15 | -0.22 | -0.41 | 0.26 | 0.61 | 0.02 |
| BIO18 | 0.39 | -0.04 | 0.06 | 0.65 | 0.26 |
| BIO19 | 0.37 | 0.41 | -0.08 | 0.14 | 0.22 |

**Supplementary table 3**

| | |
|---|---|
| **WILD-SEMIWILD** | |
| percentage points overlap | 0.91 |
| SEMIWILD percentage points overlap | 0.99 |
| WILD percentage points overlap | 0.89 |
| **WILD-LANDRACE** | |
| percentage points overlap | 0.93 |
| LANDRACE percentage points overlap | 0.99 |
| WILD percentage points overlap | 0.89 |
| **WILD-COMMERCIAL** | |
| percentage points overlap | 0.82 |
| COMMERCIAL percentage points overlap | 0.78 |
| WILD percentage points overlap | 0.98 |
| **LANDRACE-COMMERCIAL** | |
| percentage points overlap | 0.72 |
| COMMERCIAL percentage points overlap | 0.65 |
| LANDRACE percentage points overlap | 0.99 |
| **SEMIWILD-LANDRACE** | |
| percentage points overlap | 0.90 |
| SEMIWILD percentage points overlap | 0.98 |
| LANDRACE percentage points overlap | 0.88 |
| **SEMIWILD-COMMERCIAL** | |
| percentage points overlap | 0.52 |
| SEMIWILD percentage points overlap | 0.96 |
| COMMERCIAL percentage points overlap | 0.48 |

| | |
|---|---|
| **WILD-SEMIWILD** | |
| SEMIWILD hull area | 142.94 |
| WILD hull area | 214.65 |
| intersection of hulls area | 142.08 |
| SEMIWILD hull volume | 104.92 |
| WILD hull volume | 207.45 |
| intersection of hulls volume | 104.39 |
| percentage area overlap | 0.79 |
| SEMIWILDpercentage area overlap | 0.99 |
| WILDpercentage area overlap | 0.66 |

**WILD-LANDRACE**

| | |
|---|---|
| LANDRACE hull area | 163.81 |
| WILD hull area | 214.65 |
| intersection of hulls area | 144.31 |
| LANDRACE hull volume | 121.77 |
| WILD hull volume | 207.45 |
| intersection of hulls volume | 109.21 |
| percentage area overlap | 0.76 |
| LANDRACEpercentage area overlap | 0.88 |
| WILDpercentage area overlap | 0.67 |

**WILD-COMMERCIAL**

| | |
|---|---|
| COMMERCIAL hull area | 202.49 |
| WILD hull area | 214.65 |
| intersection of hulls area | 162.46 |
| COMMERCIAL hull volume | 204.22 |
| WILD hull volume | 207.45 |
| intersection of hulls volume | 154.42 |
| percentage area overlap | 0.78 |
| COMMERCIALpercentage area overlap | 0.80 |
| WILDpercentage area overlap | 0.76 |

**LANDRACE-COMMERCIAL**

| | |
|---|---|
| COMMERCIAL hull area | 202.49 |
| LANDRACE hull area | 163.81 |
| intersection of hulls area | 141.98 |
| COMMERCIAL hull volume | 204.22 |
| LANDRACE hull volume | 121.77 |
| intersection of hulls volume | 111.24 |
| percentage area overlap | 0.78 |
| COMMERCIALpercentage area overlap | 0.70 |
| LANDRACEpercentage area overlap | 0.87 |

**SEMIWILD-LANDRACE**

| | |
|---|---|
| SEMIWILD hull area | 142.94 |
| LANDRACE hull area | 163.81 |
| intersection of hulls area | 127.08 |
| SEMIWILD hull volume | 104.92 |
| LANDRACE hull volume | 121.77 |
| intersection of hulls volume | 90.18 |
| percentage area overlap | 0.83 |

| | |
|---|---|
| SEMIWILDpercentage area overlap | 0.89 |
| LANDRACEpercentage area overlap | 0.78 |

**SEMIWILD-COMMERCIAL**

| | |
|---|---|
| SEMIWILD hull area | 142.94 |
| COMMERCIAL hull area | 202.49 |
| intersection of hulls area | 128.01 |
| SEMIWILD hull volume | 104.92 |
| COMMERCIAL hull volume | 204.22 |
| intersection of hulls volume | 96.87 |
| percentage area overlap | 0.74 |
| SEMIWILDpercentage area overlap | 0.90 |
| COMMERCIALpercentage area overlap | 0.63 |

### Supplementary table 4

| Type | settings | train.AUC | avg.test.AUC | var.test.AUC | avg.diff.AUC | var.diff.AUC | avg.test.orMTP | var.test.orMTP | avg.test.or10pct | var.test.or10pct | AICc | delta.AICc | w.AIC |
|---|---|---|---|---|---|---|---|---|---|---|---|---|---|
| WILD | L_0.5 | 0.786 | 0.778 | 0.002 | 0.018 | 0.001 | 0.007 | 0.000 | 0.123 | 0.010 | 6878.700 | 120.234 | 0.000 |
| WILD | LQ_0.5 | 0.822 | 0.809 | 0.000 | 0.015 | 0.001 | 0.007 | 0.000 | 0.124 | 0.003 | 6765.157 | 6.691 | 0.034 |
| WILD | LQH_0.5 | 0.860 | 0.815 | 0.000 | 0.047 | 0.000 | 0.014 | 0.000 | 0.146 | 0.002 | 7769.080 | 1010.614 | 0.000 |
| WILD | LQHP_0.5 | 0.865 | 0.828 | 0.000 | 0.038 | 0.001 | 0.013 | 0.000 | 0.147 | 0.002 | 7774.796 | 1016.330 | 0.000 |
| WILD | L_1 | 0.786 | 0.778 | 0.002 | 0.017 | 0.001 | 0.007 | 0.000 | 0.119 | 0.009 | 6879.521 | 121.055 | 0.000 |
| WILD | LQ_1 | 0.819 | 0.806 | 0.001 | 0.016 | 0.001 | 0.018 | 0.000 | 0.125 | 0.005 | 6775.268 | 16.802 | 0.000 |
| WILD | LQH_1 | 0.854 | 0.829 | 0.000 | 0.027 | 0.000 | 0.006 | 0.000 | 0.116 | 0.003 | 6925.169 | 166.703 | 0.000 |
| WILD | LQHP_1 | 0.863 | 0.839 | 0.000 | 0.027 | 0.000 | 0.009 | 0.000 | 0.142 | 0.002 | 6881.592 | 123.126 | 0.000 |
| WILD | L_1.5 | 0.786 | 0.777 | 0.002 | 0.017 | 0.001 | 0.007 | 0.000 | 0.122 | 0.009 | 6880.794 | 122.327 | 0.000 |
| WILD | LQ_1.5 | 0.814 | 0.801 | 0.001 | 0.016 | 0.001 | 0.011 | 0.000 | 0.122 | 0.004 | 6793.051 | 34.585 | 0.000 |
| WILD | LQH_1.5 | 0.848 | 0.827 | 0.000 | 0.023 | 0.000 | 0.006 | 0.000 | 0.115 | 0.003 | 6848.421 | 89.955 | 0.000 |
| WILD | LQHP_1.5 | 0.858 | 0.836 | 0.000 | 0.023 | 0.000 | 0.006 | 0.000 | 0.120 | 0.001 | 6809.180 | 50.713 | 0.000 |
| WILD | L_2 | 0.786 | 0.777 | 0.002 | 0.017 | 0.001 | 0.007 | 0.000 | 0.122 | 0.009 | 6882.549 | 124.083 | 0.000 |
| WILD | LQ_2 | 0.809 | 0.795 | 0.001 | 0.017 | 0.001 | 0.014 | 0.000 | 0.126 | 0.006 | 6807.005 | 48.539 | 0.000 |
| WILD | LQH_2 | 0.844 | 0.823 | 0.000 | 0.020 | 0.000 | 0.004 | 0.000 | 0.119 | 0.003 | 6785.576 | 27.110 | 0.000 |
| WILD | LQHP_2 | 0.852 | 0.831 | 0.000 | 0.020 | 0.001 | 0.006 | 0.000 | 0.114 | 0.002 | 6787.745 | 29.279 | 0.000 |
| WILD | L_2.5 | 0.785 | 0.777 | 0.002 | 0.017 | 0.001 | 0.007 | 0.000 | 0.122 | 0.009 | 6884.960 | 126.494 | 0.000 |
| WILD | LQ_2.5 | 0.802 | 0.790 | 0.001 | 0.017 | 0.001 | 0.011 | 0.000 | 0.133 | 0.006 | 6824.220 | 65.753 | 0.000 |
| WILD | LQH_2.5 | 0.839 | 0.819 | 0.000 | 0.018 | 0.001 | 0.004 | 0.000 | 0.110 | 0.003 | 6776.380 | 17.914 | 0.000 |
| WILD | LQHP_2.5 | 0.846 | 0.826 | 0.000 | 0.017 | 0.001 | 0.006 | 0.000 | 0.114 | 0.002 | 6772.671 | 14.204 | 0.001 |
| WILD | L_3 | 0.784 | 0.776 | 0.001 | 0.017 | 0.001 | 0.004 | 0.000 | 0.135 | 0.010 | 6887.629 | 129.163 | 0.000 |
| WILD | LQ_3 | 0.799 | 0.787 | 0.001 | 0.017 | 0.001 | 0.011 | 0.000 | 0.123 | 0.004 | 6829.948 | 71.482 | 0.000 |
| WILD | LQH_3 | 0.833 | 0.815 | 0.001 | 0.016 | 0.001 | 0.004 | 0.000 | 0.106 | 0.003 | 6779.151 | 20.685 | 0.000 |
| WILD | LQHP_3 | 0.839 | 0.821 | 0.001 | 0.017 | 0.001 | 0.006 | 0.000 | 0.107 | 0.003 | 6758.466 | 0.000 | 0.965 |
| WILD | L_3.5 | 0.784 | 0.776 | 0.001 | 0.017 | 0.001 | 0.006 | 0.000 | 0.134 | 0.010 | 6890.909 | 132.443 | 0.000 |
| WILD | LQ_3.5 | 0.797 | 0.786 | 0.001 | 0.018 | 0.001 | 0.007 | 0.000 | 0.117 | 0.004 | 6836.132 | 77.666 | 0.000 |
| WILD | LQH_3.5 | 0.828 | 0.811 | 0.001 | 0.016 | 0.001 | 0.004 | 0.000 | 0.109 | 0.003 | 6783.048 | 24.582 | 0.000 |
| WILD | LQHP_3.5 | 0.832 | 0.816 | 0.001 | 0.017 | 0.001 | 0.006 | 0.000 | 0.109 | 0.003 | 6779.521 | 21.055 | 0.000 |
| WILD-SL | L_0.5 | 0.790 | 0.783 | 0.002 | 0.015 | 0.001 | 0.005 | 0.000 | 0.115 | 0.007 | 8328.984 | 133.684 | 0.000 |
| WILD-SL | LQ_0.5 | 0.826 | 0.816 | 0.001 | 0.015 | 0.001 | 0.010 | 0.000 | 0.127 | 0.001 | 8208.886 | 13.587 | 0.001 |
| WILD-SL | LQH_0.5 | 0.859 | 0.831 | 0.000 | 0.032 | 0.000 | 0.013 | 0.000 | 0.133 | 0.001 | 9085.509 | 890.209 | 0.000 |
| WILD-SL | LQHP_0.5 | 0.871 | 0.840 | 0.000 | 0.031 | 0.000 | 0.011 | 0.000 | 0.139 | 0.000 | 8892.194 | 696.895 | 0.000 |

| | | | | | | | | | | | | | |
|---|---|---|---|---|---|---|---|---|---|---|---|---|---|
| WILD-SL | L_1 | 0.789 | 0.783 | 0.002 | 0.015 | 0.001 | 0.005 | 0.000 | 0.118 | 0.007 | 8330.184 | 134.885 | 0.000 |
| WILD-SL | LQ_1 | 0.821 | 0.812 | 0.001 | 0.015 | 0.002 | 0.010 | 0.000 | 0.117 | 0.003 | 8216.933 | 21.633 | 0.000 |
| WILD-SL | LQH_1 | 0.855 | 0.834 | 0.001 | 0.023 | 0.001 | 0.005 | 0.000 | 0.130 | 0.002 | 8375.950 | 180.651 | 0.000 |
| WILD-SL | LQHP_1 | 0.864 | 0.843 | 0.001 | 0.023 | 0.001 | 0.005 | 0.000 | 0.135 | 0.002 | 8280.551 | 85.252 | 0.000 |
| WILD-SL | L_1.5 | 0.789 | 0.783 | 0.002 | 0.015 | 0.001 | 0.005 | 0.000 | 0.115 | 0.007 | 8331.984 | 136.684 | 0.000 |
| WILD-SL | LQ_1.5 | 0.816 | 0.807 | 0.002 | 0.015 | 0.002 | 0.010 | 0.000 | 0.111 | 0.003 | 8236.324 | 41.024 | 0.000 |
| WILD-SL | LQH_1.5 | 0.851 | 0.831 | 0.001 | 0.020 | 0.001 | 0.005 | 0.000 | 0.117 | 0.002 | 8282.726 | 87.427 | 0.000 |
| WILD-SL | LQHP_1.5 | 0.859 | 0.840 | 0.001 | 0.020 | 0.001 | 0.005 | 0.000 | 0.118 | 0.003 | 8217.681 | 22.381 | 0.000 |
| WILD-SL | L_2 | 0.789 | 0.782 | 0.002 | 0.015 | 0.001 | 0.005 | 0.000 | 0.113 | 0.007 | 8334.372 | 139.072 | 0.000 |
| WILD-SL | LQ_2 | 0.810 | 0.802 | 0.002 | 0.015 | 0.002 | 0.008 | 0.000 | 0.113 | 0.004 | 8254.956 | 59.656 | 0.000 |
| WILD-SL | LQH_2 | 0.845 | 0.829 | 0.001 | 0.018 | 0.001 | 0.005 | 0.000 | 0.117 | 0.002 | 8291.053 | 95.753 | 0.000 |
| WILD-SL | LQHP_2 | 0.853 | 0.837 | 0.001 | 0.018 | 0.001 | 0.010 | 0.000 | 0.103 | 0.001 | 8243.466 | 48.167 | 0.000 |
| WILD-SL | L_2.5 | 0.788 | 0.782 | 0.002 | 0.015 | 0.001 | 0.005 | 0.000 | 0.115 | 0.007 | 8337.221 | 141.921 | 0.000 |
| WILD-SL | LQ_2.5 | 0.804 | 0.796 | 0.002 | 0.015 | 0.002 | 0.008 | 0.000 | 0.109 | 0.004 | 8272.260 | 76.960 | 0.000 |
| WILD-SL | LQH_2.5 | 0.841 | 0.827 | 0.001 | 0.016 | 0.001 | 0.005 | 0.000 | 0.119 | 0.002 | 8247.192 | 51.892 | 0.000 |
| WILD-SL | LQHP_2.5 | 0.848 | 0.833 | 0.001 | 0.017 | 0.001 | 0.010 | 0.000 | 0.108 | 0.001 | 8218.637 | 23.338 | 0.000 |
| WILD-SL | L_3 | 0.787 | 0.781 | 0.002 | 0.014 | 0.001 | 0.005 | 0.000 | 0.118 | 0.006 | 8338.418 | 143.119 | 0.000 |
| WILD-SL | LQ_3 | 0.801 | 0.793 | 0.002 | 0.015 | 0.002 | 0.008 | 0.000 | 0.106 | 0.003 | 8279.461 | 84.161 | 0.000 |
| WILD-SL | LQH_3 | 0.837 | 0.824 | 0.001 | 0.016 | 0.001 | 0.005 | 0.000 | 0.108 | 0.002 | 8242.499 | 47.200 | 0.000 |
| WILD-SL | LQHP_3 | 0.841 | 0.830 | 0.001 | 0.017 | 0.001 | 0.008 | 0.000 | 0.103 | 0.001 | 8195.300 | 0.000 | 0.788 |
| WILD-SL | L_3.5 | 0.786 | 0.781 | 0.001 | 0.014 | 0.001 | 0.005 | 0.000 | 0.124 | 0.008 | 8341.351 | 146.052 | 0.000 |
| WILD-SL | LQ_3.5 | 0.799 | 0.792 | 0.002 | 0.015 | 0.002 | 0.005 | 0.000 | 0.109 | 0.004 | 8288.134 | 92.834 | 0.000 |
| WILD-SL | LQH_3.5 | 0.831 | 0.820 | 0.001 | 0.015 | 0.001 | 0.005 | 0.000 | 0.106 | 0.002 | 8229.115 | 33.815 | 0.000 |
| WILD-SL | LQHP_3.5 | 0.837 | 0.828 | 0.001 | 0.016 | 0.001 | 0.008 | 0.000 | 0.103 | 0.002 | 8197.930 | 2.630 | 0.211 |
| SEMIWILD | L_0.5 | 0.867 | 0.856 | 0.002 | 0.022 | 0.001 | 0.012 | 0.001 | 0.122 | 0.009 | 1892.100 | 17.748 | 0.000 |
| SEMIWILD | LQ_0.5 | 0.885 | 0.867 | 0.001 | 0.022 | 0.000 | 0.012 | 0.001 | 0.214 | 0.004 | 1882.770 | 8.418 | 0.007 |
| SEMIWILD | LQH_0.5 | 0.915 | 0.875 | 0.001 | 0.043 | 0.001 | 0.012 | 0.001 | 0.205 | 0.005 | NA | NA | NA |
| SEMIWILD | LQHP_0.5 | 0.918 | 0.872 | 0.000 | 0.048 | 0.000 | 0.012 | 0.001 | 0.270 | 0.021 | NA | NA | NA |
| SEMIWILD | L_1 | 0.867 | 0.857 | 0.001 | 0.021 | 0.001 | 0.012 | 0.001 | 0.134 | 0.010 | 1892.499 | 18.147 | 0.000 |
| SEMIWILD | LQ_1 | 0.883 | 0.863 | 0.001 | 0.022 | 0.001 | 0.012 | 0.001 | 0.153 | 0.007 | 1878.103 | 3.751 | 0.075 |
| SEMIWILD | LQH_1 | 0.901 | 0.873 | 0.001 | 0.031 | 0.001 | 0.012 | 0.001 | 0.172 | 0.005 | 2099.756 | 225.404 | 0.000 |
| SEMIWILD | LQHP_1 | 0.905 | 0.870 | 0.001 | 0.036 | 0.001 | 0.012 | 0.001 | 0.231 | 0.010 | 2177.596 | 303.244 | 0.000 |
| SEMIWILD | L_1.5 | 0.867 | 0.857 | 0.001 | 0.020 | 0.001 | 0.012 | 0.001 | 0.145 | 0.013 | 1890.598 | 16.246 | 0.000 |
| SEMIWILD | LQ_1.5 | 0.879 | 0.858 | 0.001 | 0.023 | 0.001 | 0.012 | 0.001 | 0.187 | 0.016 | 1879.723 | 5.370 | 0.033 |
| SEMIWILD | LQH_1.5 | 0.893 | 0.867 | 0.001 | 0.026 | 0.001 | 0.012 | 0.001 | 0.182 | 0.008 | 1895.446 | 21.093 | 0.000 |
| SEMIWILD | LQHP_1.5 | 0.895 | 0.869 | 0.001 | 0.026 | 0.001 | 0.012 | 0.001 | 0.167 | 0.007 | 1914.355 | 40.002 | 0.000 |
| SEMIWILD | L_2 | 0.867 | 0.856 | 0.001 | 0.019 | 0.001 | 0.012 | 0.001 | 0.155 | 0.010 | 1888.836 | 14.484 | 0.000 |
| SEMIWILD | LQ_2 | 0.876 | 0.856 | 0.001 | 0.022 | 0.001 | 0.012 | 0.001 | 0.164 | 0.007 | 1881.393 | 7.041 | 0.014 |
| SEMIWILD | LQH_2 | 0.888 | 0.863 | 0.001 | 0.025 | 0.001 | 0.012 | 0.001 | 0.182 | 0.008 | 1876.237 | 1.885 | 0.190 |

| | | | | | | | | | | | | | |
|---|---|---|---|---|---|---|---|---|---|---|---|---|---|
| SEMIWILD | LQHP_2 | 0.892 | 0.865 | 0.001 | 0.022 | 0.001 | 0.012 | 0.001 | 0.141 | 0.004 | 1880.355 | 6.002 | 0.024 |
| SEMIWILD | L_2.5 | 0.867 | 0.855 | 0.001 | 0.018 | 0.001 | 0.012 | 0.001 | 0.140 | 0.012 | 1889.514 | 15.162 | 0.000 |
| SEMIWILD | LQ_2.5 | 0.874 | 0.854 | 0.001 | 0.022 | 0.001 | 0.012 | 0.001 | 0.176 | 0.011 | 1885.186 | 10.834 | 0.002 |
| SEMIWILD | LQH_2.5 | 0.884 | 0.860 | 0.001 | 0.023 | 0.001 | 0.012 | 0.001 | 0.171 | 0.006 | 1874.352 | 0.000 | 0.487 |
| SEMIWILD | LQHP_2.5 | 0.888 | 0.863 | 0.001 | 0.022 | 0.001 | 0.012 | 0.001 | 0.141 | 0.004 | 1878.592 | 4.240 | 0.058 |
| SEMIWILD | L_3 | 0.866 | 0.853 | 0.001 | 0.017 | 0.001 | 0.012 | 0.001 | 0.137 | 0.021 | 1890.337 | 15.985 | 0.000 |
| SEMIWILD | LQ_3 | 0.870 | 0.851 | 0.001 | 0.021 | 0.001 | 0.012 | 0.001 | 0.160 | 0.034 | 1889.640 | 15.288 | 0.000 |
| SEMIWILD | LQH_3 | 0.882 | 0.858 | 0.001 | 0.023 | 0.001 | 0.012 | 0.001 | 0.194 | 0.007 | 1883.291 | 8.939 | 0.006 |
| SEMIWILD | LQHP_3 | 0.886 | 0.859 | 0.001 | 0.022 | 0.001 | 0.012 | 0.001 | 0.153 | 0.005 | 1881.673 | 7.321 | 0.013 |
| SEMIWILD | L_3.5 | 0.865 | 0.850 | 0.001 | 0.017 | 0.001 | 0.012 | 0.001 | 0.137 | 0.021 | 1891.279 | 16.927 | 0.000 |
| SEMIWILD | LQ_3.5 | 0.869 | 0.850 | 0.001 | 0.021 | 0.001 | 0.012 | 0.001 | 0.160 | 0.034 | 1889.077 | 14.725 | 0.000 |
| SEMIWILD | LQH_3.5 | 0.879 | 0.857 | 0.001 | 0.022 | 0.001 | 0.012 | 0.001 | 0.150 | 0.007 | 1881.501 | 7.149 | 0.014 |
| SEMIWILD | LQHP_3.5 | 0.883 | 0.857 | 0.001 | 0.022 | 0.001 | 0.012 | 0.001 | 0.143 | 0.007 | 1878.097 | 3.744 | 0.075 |
| LANDRACE | L_0.5 | 0.894 | 0.892 | 0.000 | 0.006 | 0.000 | 0.006 | 0.000 | 0.100 | 0.003 | 5926.407 | 44.975 | 0.000 |
| LANDRACE | LQ_0.5 | 0.904 | 0.900 | 0.000 | 0.007 | 0.000 | 0.003 | 0.000 | 0.103 | 0.004 | 5881.432 | 0.000 | 0.546 |
| LANDRACE | LQH_0.5 | 0.922 | 0.903 | 0.000 | 0.021 | 0.000 | 0.006 | 0.000 | 0.140 | 0.003 | 6666.099 | 784.667 | 0.000 |
| LANDRACE | LQHP_0.5 | 0.923 | 0.904 | 0.000 | 0.022 | 0.000 | 0.006 | 0.000 | 0.170 | 0.001 | 6553.486 | 672.054 | 0.000 |
| LANDRACE | L_1 | 0.894 | 0.892 | 0.000 | 0.006 | 0.000 | 0.006 | 0.000 | 0.100 | 0.003 | 5928.181 | 46.749 | 0.000 |
| LANDRACE | LQ_1 | 0.897 | 0.895 | 0.000 | 0.008 | 0.000 | 0.006 | 0.000 | 0.103 | 0.003 | 5917.663 | 36.231 | 0.000 |
| LANDRACE | LQH_1 | 0.916 | 0.905 | 0.000 | 0.012 | 0.000 | 0.003 | 0.000 | 0.132 | 0.005 | 5990.235 | 108.803 | 0.000 |
| LANDRACE | LQHP_1 | 0.920 | 0.910 | 0.000 | 0.012 | 0.000 | 0.003 | 0.000 | 0.134 | 0.003 | 6070.363 | 188.931 | 0.000 |
| LANDRACE | L_1.5 | 0.893 | 0.892 | 0.000 | 0.006 | 0.000 | 0.006 | 0.000 | 0.100 | 0.003 | 5930.964 | 49.532 | 0.000 |
| LANDRACE | LQ_1.5 | 0.895 | 0.894 | 0.000 | 0.007 | 0.000 | 0.006 | 0.000 | 0.103 | 0.003 | 5922.540 | 41.108 | 0.000 |
| LANDRACE | LQH_1.5 | 0.914 | 0.905 | 0.000 | 0.010 | 0.000 | 0.003 | 0.000 | 0.119 | 0.004 | 5955.917 | 74.485 | 0.000 |
| LANDRACE | LQHP_1.5 | 0.917 | 0.908 | 0.000 | 0.012 | 0.000 | 0.003 | 0.000 | 0.122 | 0.003 | 5972.954 | 91.522 | 0.000 |
| LANDRACE | L_2 | 0.893 | 0.892 | 0.000 | 0.006 | 0.000 | 0.006 | 0.000 | 0.103 | 0.003 | 5934.553 | 53.121 | 0.000 |
| LANDRACE | LQ_2 | 0.894 | 0.893 | 0.000 | 0.006 | 0.000 | 0.006 | 0.000 | 0.107 | 0.003 | 5922.644 | 41.212 | 0.000 |
| LANDRACE | LQH_2 | 0.912 | 0.904 | 0.000 | 0.009 | 0.000 | 0.003 | 0.000 | 0.119 | 0.003 | 5929.626 | 48.194 | 0.000 |
| LANDRACE | LQHP_2 | 0.915 | 0.907 | 0.000 | 0.011 | 0.000 | 0.003 | 0.000 | 0.113 | 0.003 | 5924.217 | 42.785 | 0.000 |
| LANDRACE | L_2.5 | 0.892 | 0.892 | 0.000 | 0.006 | 0.000 | 0.006 | 0.000 | 0.106 | 0.002 | 5936.264 | 54.832 | 0.000 |
| LANDRACE | LQ_2.5 | 0.894 | 0.893 | 0.000 | 0.006 | 0.000 | 0.006 | 0.000 | 0.103 | 0.004 | 5927.843 | 46.411 | 0.000 |
| LANDRACE | LQH_2.5 | 0.911 | 0.902 | 0.000 | 0.009 | 0.000 | 0.003 | 0.000 | 0.114 | 0.004 | 5936.385 | 54.953 | 0.000 |
| LANDRACE | LQHP_2.5 | 0.913 | 0.904 | 0.001 | 0.011 | 0.000 | 0.003 | 0.000 | 0.113 | 0.003 | 5896.256 | 14.824 | 0.000 |
| LANDRACE | L_3 | 0.891 | 0.891 | 0.000 | 0.006 | 0.000 | 0.003 | 0.000 | 0.110 | 0.003 | 5940.476 | 59.044 | 0.000 |
| LANDRACE | LQ_3 | 0.893 | 0.893 | 0.000 | 0.006 | 0.000 | 0.006 | 0.000 | 0.113 | 0.003 | 5929.731 | 48.299 | 0.000 |
| LANDRACE | LQH_3 | 0.909 | 0.901 | 0.000 | 0.009 | 0.000 | 0.003 | 0.000 | 0.117 | 0.004 | 5941.494 | 60.062 | 0.000 |
| LANDRACE | LQHP_3 | 0.911 | 0.902 | 0.001 | 0.011 | 0.000 | 0.003 | 0.000 | 0.113 | 0.003 | 5881.820 | 0.388 | 0.450 |
| LANDRACE | L_3.5 | 0.891 | 0.891 | 0.000 | 0.006 | 0.000 | 0.003 | 0.000 | 0.110 | 0.003 | 5941.857 | 60.425 | 0.000 |
| LANDRACE | LQ_3.5 | 0.893 | 0.892 | 0.000 | 0.006 | 0.000 | 0.006 | 0.000 | 0.113 | 0.003 | 5933.547 | 52.115 | 0.000 |

| | | | | | | | | | | | | | |
|---|---|---|---|---|---|---|---|---|---|---|---|---|---|
| LANDRACE | LQH_3.5 | 0.908 | 0.898 | 0.000 | 0.009 | 0.000 | 0.003 | 0.000 | 0.119 | 0.004 | 5949.067 | 67.635 | 0.000 |
| LANDRACE | LQHP_3.5 | 0.908 | 0.901 | 0.001 | 0.011 | 0.000 | 0.003 | 0.000 | 0.113 | 0.003 | 5891.034 | 9.602 | 0.004 |
| COMMERCIAL | L_0.5 | 0.771 | 0.770 | 0.000 | 0.004 | 0.000 | 0.000 | 0.000 | 0.100 | 0.001 | 50085.992 | 1076.928 | 0.000 |
| COMMERCIAL | LQ_0.5 | 0.782 | 0.780 | 0.000 | 0.005 | 0.000 | 0.001 | 0.000 | 0.105 | 0.001 | 49684.414 | 675.351 | 0.000 |
| COMMERCIAL | LQH_0.5 | 0.827 | 0.820 | 0.000 | 0.009 | 0.000 | 0.001 | 0.000 | 0.110 | 0.000 | 49108.903 | 99.839 | 0.000 |
| COMMERCIAL | LQHP_0.5 | 0.830 | 0.824 | 0.000 | 0.009 | 0.000 | 0.000 | 0.000 | 0.108 | 0.000 | 49009.064 | 0.000 | 1.000 |
| COMMERCIAL | L_1 | 0.770 | 0.769 | 0.000 | 0.004 | 0.000 | 0.000 | 0.000 | 0.099 | 0.001 | 50101.880 | 1092.817 | 0.000 |
| COMMERCIAL | LQ_1 | 0.780 | 0.779 | 0.000 | 0.005 | 0.000 | 0.001 | 0.000 | 0.103 | 0.001 | 49747.271 | 738.208 | 0.000 |
| COMMERCIAL | LQH_1 | 0.822 | 0.817 | 0.000 | 0.008 | 0.000 | 0.001 | 0.000 | 0.102 | 0.000 | 49121.548 | 112.484 | 0.000 |
| COMMERCIAL | LQHP_1 | 0.823 | 0.818 | 0.000 | 0.008 | 0.000 | 0.001 | 0.000 | 0.105 | 0.000 | 49044.365 | 35.301 | 0.000 |
| COMMERCIAL | L_1.5 | 0.769 | 0.768 | 0.000 | 0.003 | 0.000 | 0.000 | 0.000 | 0.101 | 0.001 | 50123.529 | 1114.465 | 0.000 |
| COMMERCIAL | LQ_1.5 | 0.778 | 0.776 | 0.000 | 0.004 | 0.000 | 0.000 | 0.000 | 0.103 | 0.000 | 49821.465 | 812.402 | 0.000 |
| COMMERCIAL | LQH_1.5 | 0.819 | 0.814 | 0.000 | 0.008 | 0.000 | 0.001 | 0.000 | 0.105 | 0.000 | 49124.876 | 115.812 | 0.000 |
| COMMERCIAL | LQHP_1.5 | 0.820 | 0.816 | 0.000 | 0.007 | 0.000 | 0.000 | 0.000 | 0.105 | 0.000 | 49071.384 | 62.320 | 0.000 |
| COMMERCIAL | L_2 | 0.768 | 0.767 | 0.000 | 0.003 | 0.000 | 0.000 | 0.000 | 0.098 | 0.001 | 50140.491 | 1131.427 | 0.000 |
| COMMERCIAL | LQ_2 | 0.775 | 0.774 | 0.000 | 0.004 | 0.000 | 0.000 | 0.000 | 0.100 | 0.000 | 49913.374 | 904.310 | 0.000 |
| COMMERCIAL | LQH_2 | 0.817 | 0.812 | 0.000 | 0.007 | 0.000 | 0.001 | 0.000 | 0.104 | 0.000 | 49183.643 | 174.579 | 0.000 |
| COMMERCIAL | LQHP_2 | 0.818 | 0.813 | 0.000 | 0.007 | 0.000 | 0.000 | 0.000 | 0.106 | 0.000 | 49116.457 | 107.394 | 0.000 |
| COMMERCIAL | L_2.5 | 0.766 | 0.765 | 0.000 | 0.003 | 0.000 | 0.000 | 0.000 | 0.099 | 0.001 | 50159.857 | 1150.794 | 0.000 |
| COMMERCIAL | LQ_2.5 | 0.772 | 0.770 | 0.000 | 0.004 | 0.000 | 0.000 | 0.000 | 0.102 | 0.000 | 50000.066 | 991.002 | 0.000 |
| COMMERCIAL | LQH_2.5 | 0.814 | 0.809 | 0.000 | 0.007 | 0.000 | 0.000 | 0.000 | 0.105 | 0.000 | 49224.245 | 215.181 | 0.000 |
| COMMERCIAL | LQHP_2.5 | 0.814 | 0.810 | 0.000 | 0.007 | 0.000 | 0.000 | 0.000 | 0.104 | 0.000 | 49167.172 | 158.109 | 0.000 |
| COMMERCIAL | L_3 | 0.764 | 0.763 | 0.000 | 0.003 | 0.000 | 0.000 | 0.000 | 0.103 | 0.001 | 50182.798 | 1173.735 | 0.000 |
| COMMERCIAL | LQ_3 | 0.770 | 0.768 | 0.000 | 0.003 | 0.000 | 0.000 | 0.000 | 0.102 | 0.000 | 50048.730 | 1039.666 | 0.000 |
| COMMERCIAL | LQH_3 | 0.810 | 0.806 | 0.000 | 0.006 | 0.000 | 0.001 | 0.000 | 0.106 | 0.000 | 49291.248 | 282.184 | 0.000 |
| COMMERCIAL | LQHP_3 | 0.811 | 0.807 | 0.000 | 0.006 | 0.000 | 0.000 | 0.000 | 0.106 | 0.000 | 49238.984 | 229.920 | 0.000 |
| COMMERCIAL | L_3.5 | 0.762 | 0.761 | 0.000 | 0.003 | 0.000 | 0.000 | 0.000 | 0.097 | 0.001 | 50209.331 | 1200.268 | 0.000 |
| COMMERCIAL | LQ_3.5 | 0.767 | 0.766 | 0.000 | 0.003 | 0.000 | 0.000 | 0.000 | 0.102 | 0.000 | 50099.652 | 1090.589 | 0.000 |
| COMMERCIAL | LQH_3.5 | 0.807 | 0.803 | 0.000 | 0.006 | 0.000 | 0.001 | 0.000 | 0.106 | 0.000 | 49367.756 | 358.693 | 0.000 |
| COMMERCIAL | LQHP_3.5 | 0.808 | 0.805 | 0.000 | 0.006 | 0.000 | 0.000 | 0.000 | 0.105 | 0.000 | 49277.559 | 268.496 | 0.000 |
| CULTIVATED | L_0.5 | 0.755 | 0.753 | 0.000 | 0.003 | 0.000 | 0.000 | 0.000 | 0.099 | 0.001 | 57084.903 | 1201.896 | 0.000 |
| CULTIVATED | LQ_0.5 | 0.766 | 0.764 | 0.000 | 0.003 | 0.000 | 0.000 | 0.000 | 0.102 | 0.000 | 56676.801 | 793.794 | 0.000 |
| CULTIVATED | LQH_0.5 | 0.812 | 0.805 | 0.000 | 0.007 | 0.000 | 0.001 | 0.000 | 0.106 | 0.000 | 55993.480 | 110.474 | 0.000 |
| CULTIVATED | LQHP_0.5 | 0.815 | 0.809 | 0.000 | 0.007 | 0.000 | 0.001 | 0.000 | 0.107 | 0.000 | 55883.006 | 0.000 | 1.000 |
| CULTIVATED | L_1 | 0.754 | 0.752 | 0.000 | 0.003 | 0.000 | 0.000 | 0.000 | 0.102 | 0.000 | 57104.049 | 1221.043 | 0.000 |
| CULTIVATED | LQ_1 | 0.764 | 0.763 | 0.000 | 0.003 | 0.000 | 0.000 | 0.000 | 0.103 | 0.000 | 56725.868 | 842.862 | 0.000 |
| CULTIVATED | LQH_1 | 0.808 | 0.802 | 0.000 | 0.006 | 0.000 | 0.001 | 0.000 | 0.105 | 0.000 | 56044.408 | 161.402 | 0.000 |
| CULTIVATED | LQHP_1 | 0.811 | 0.805 | 0.000 | 0.006 | 0.000 | 0.001 | 0.000 | 0.109 | 0.000 | 55909.639 | 26.633 | 0.000 |

| | | | | | | | | | | | | | |
|---|---|---|---|---|---|---|---|---|---|---|---|---|---|
| CULTIVATED | L_1.5 | 0.753 | 0.751 | 0.000 | 0.003 | 0.000 | 0.000 | 0.000 | 0.101 | 0.000 | 57121.810 | 1238.803 | 0.000 |
| CULTIVATED | LQ_1.5 | 0.763 | 0.761 | 0.000 | 0.003 | 0.000 | 0.000 | 0.000 | 0.104 | 0.000 | 56797.604 | 914.598 | 0.000 |
| CULTIVATED | LQH_1.5 | 0.805 | 0.800 | 0.000 | 0.006 | 0.000 | 0.002 | 0.000 | 0.107 | 0.000 | 56090.547 | 207.540 | 0.000 |
| CULTIVATED | LQHP_1.5 | 0.807 | 0.803 | 0.000 | 0.005 | 0.000 | 0.000 | 0.000 | 0.104 | 0.000 | 56019.681 | 136.674 | 0.000 |
| CULTIVATED | L_2 | 0.751 | 0.749 | 0.000 | 0.002 | 0.000 | 0.000 | 0.000 | 0.102 | 0.000 | 57144.222 | 1261.216 | 0.000 |
| CULTIVATED | LQ_2 | 0.760 | 0.758 | 0.000 | 0.002 | 0.000 | 0.000 | 0.000 | 0.106 | 0.000 | 56874.193 | 991.186 | 0.000 |
| CULTIVATED | LQH_2 | 0.803 | 0.798 | 0.000 | 0.005 | 0.000 | 0.001 | 0.000 | 0.105 | 0.000 | 56130.841 | 247.834 | 0.000 |
| CULTIVATED | LQHP_2 | 0.805 | 0.800 | 0.000 | 0.005 | 0.000 | 0.000 | 0.000 | 0.103 | 0.000 | 56069.798 | 186.792 | 0.000 |
| CULTIVATED | L_2.5 | 0.748 | 0.747 | 0.000 | 0.002 | 0.000 | 0.000 | 0.000 | 0.102 | 0.000 | 57173.899 | 1290.893 | 0.000 |
| CULTIVATED | LQ_2.5 | 0.756 | 0.755 | 0.000 | 0.002 | 0.000 | 0.000 | 0.000 | 0.103 | 0.000 | 56972.379 | 1089.373 | 0.000 |
| CULTIVATED | LQH_2.5 | 0.800 | 0.796 | 0.000 | 0.004 | 0.000 | 0.000 | 0.000 | 0.105 | 0.000 | 56159.273 | 276.266 | 0.000 |
| CULTIVATED | LQHP_2.5 | 0.802 | 0.797 | 0.000 | 0.004 | 0.000 | 0.000 | 0.000 | 0.103 | 0.000 | 56105.924 | 222.918 | 0.000 |
| CULTIVATED | L_3 | 0.746 | 0.745 | 0.000 | 0.002 | 0.000 | 0.000 | 0.000 | 0.102 | 0.000 | 57197.263 | 1314.256 | 0.000 |
| CULTIVATED | LQ_3 | 0.753 | 0.752 | 0.000 | 0.002 | 0.000 | 0.000 | 0.000 | 0.103 | 0.000 | 57037.682 | 1154.676 | 0.000 |
| CULTIVATED | LQH_3 | 0.797 | 0.792 | 0.000 | 0.004 | 0.000 | 0.000 | 0.000 | 0.104 | 0.000 | 56251.813 | 368.807 | 0.000 |
| CULTIVATED | LQHP_3 | 0.799 | 0.794 | 0.000 | 0.004 | 0.000 | 0.000 | 0.000 | 0.103 | 0.000 | 56205.463 | 322.456 | 0.000 |
| CULTIVATED | L_3.5 | 0.745 | 0.744 | 0.000 | 0.002 | 0.000 | 0.000 | 0.000 | 0.103 | 0.000 | 57225.408 | 1342.402 | 0.000 |
| CULTIVATED | LQ_3.5 | 0.751 | 0.750 | 0.000 | 0.002 | 0.000 | 0.000 | 0.000 | 0.100 | 0.000 | 57075.357 | 1192.350 | 0.000 |
| CULTIVATED | LQH_3.5 | 0.793 | 0.789 | 0.000 | 0.003 | 0.000 | 0.000 | 0.000 | 0.103 | 0.000 | 56301.450 | 418.444 | 0.000 |
| CULTIVATED | LQHP_3.5 | 0.796 | 0.791 | 0.000 | 0.004 | 0.000 | 0.000 | 0.000 | 0.101 | 0.000 | 56229.289 | 346.283 | 0.000 |

**Supplementary table 5**

### Single GCM models

| year | ssp | GCM | Dom | none | pixLost | pixNew | pixKeep | percent |
|---|---|---|---|---|---|---|---|---|
| 2050 | 45 | BCC-CSM2-MR | SEMIWILD | 330066 | 891 | 14900 | 71252 | 90.02 |
| 2050 | 45 | CNRM-CM6-1 | SEMIWILD | 328664 | 1015 | 16302 | 71128 | 89.15 |
| 2050 | 45 | CNRM-ESM2-1 | SEMIWILD | 331587 | 2257 | 13379 | 69886 | 89.94 |
| 2050 | 45 | CanESM5 | SEMIWILD | 319371 | 1634 | 25595 | 70509 | 83.82 |
| 2050 | 45 | IPSL-CM6A-LR | SEMIWILD | 328026 | 2110 | 16940 | 70033 | 88.03 |
| 2050 | 45 | MIROC-ES2L | SEMIWILD | 332945 | 4055 | 12021 | 68088 | 89.44 |
| 2050 | 45 | MIROC6 | SEMIWILD | 328663 | 4926 | 16303 | 67217 | 86.36 |
| 2050 | 45 | MRI-ESM2-0 | SEMIWILD | 327540 | 3498 | 17426 | 68645 | 86.77 |
| 2050 | 85 | BCC-CSM2-MR | SEMIWILD | 324766 | 185 | 20200 | 71958 | 87.59 |
| 2050 | 85 | CNRM-CM6-1 | SEMIWILD | 329759 | 1341 | 15207 | 70802 | 89.54 |
| 2050 | 85 | CNRM-ESM2-1 | SEMIWILD | 329014 | 1658 | 15952 | 70485 | 88.9 |
| 2050 | 85 | CanESM5 | SEMIWILD | 312962 | 1706 | 32004 | 70437 | 80.69 |
| 2050 | 85 | IPSL-CM6A-LR | SEMIWILD | 324962 | 1572 | 20004 | 70571 | 86.74 |
| 2050 | 85 | MIROC-ES2L | SEMIWILD | 331775 | 7772 | 13191 | 64371 | 86 |
| 2050 | 85 | MIROC6 | SEMIWILD | 325530 | 4778 | 19436 | 67365 | 84.77 |
| 2050 | 85 | MRI-ESM2-0 | SEMIWILD | 329832 | 4622 | 15134 | 67521 | 87.24 |
| 2070 | 45 | BCC-CSM2-MR | SEMIWILD | 324791 | 252 | 20175 | 71891 | 87.56 |
| 2070 | 45 | CNRM-CM6-1 | SEMIWILD | 327726 | 1400 | 17240 | 70743 | 88.36 |
| 2070 | 45 | CNRM-ESM2-1 | SEMIWILD | 328896 | 2274 | 16070 | 69869 | 88.4 |
| 2070 | 45 | CanESM5 | SEMIWILD | 314680 | 1652 | 30286 | 70491 | 81.53 |
| 2070 | 45 | IPSL-CM6A-LR | SEMIWILD | 326584 | 1751 | 18382 | 70392 | 87.49 |
| 2070 | 45 | MIROC-ES2L | SEMIWILD | 329507 | 6881 | 15459 | 65262 | 85.39 |
| 2070 | 45 | MIROC6 | SEMIWILD | 325101 | 5312 | 19865 | 66831 | 84.15 |
| 2070 | 45 | MRI-ESM2-0 | SEMIWILD | 326559 | 2555 | 18407 | 69588 | 86.91 |
| 2070 | 85 | BCC-CSM2-MR | SEMIWILD | 316336 | 283 | 28630 | 71860 | 83.25 |
| 2070 | 85 | CNRM-CM6-1 | SEMIWILD | 324151 | 1089 | 20815 | 71054 | 86.64 |
| 2070 | 85 | CNRM-ESM2-1 | SEMIWILD | 325394 | 2328 | 19572 | 69815 | 86.44 |
| 2070 | 85 | CanESM5 | SEMIWILD | 301767 | 1435 | 43199 | 70708 | 76.01 |
| 2070 | 85 | IPSL-CM6A-LR | SEMIWILD | 316719 | 1547 | 28247 | 70596 | 82.58 |
| 2070 | 85 | MIROC-ES2L | SEMIWILD | 325127 | 10243 | 19839 | 61900 | 80.45 |
| 2070 | 85 | MIROC6 | SEMIWILD | 313037 | 3863 | 31929 | 68280 | 79.23 |
| 2070 | 85 | MRI-ESM2-0 | SEMIWILD | 320785 | 4205 | 24181 | 67938 | 82.72 |
| 2090 | 45 | BCC-CSM2-MR | SEMIWILD | 326445 | 704 | 18521 | 71439 | 88.14 |
| 2090 | 45 | CNRM-CM6-1 | SEMIWILD | 326663 | 1100 | 18303 | 71043 | 87.98 |
| 2090 | 45 | CNRM-ESM2-1 | SEMIWILD | 327396 | 1872 | 17570 | 70271 | 87.85 |
| 2090 | 45 | CanESM5 | SEMIWILD | 313060 | 1732 | 31906 | 70411 | 80.72 |
| 2090 | 45 | IPSL-CM6A-LR | SEMIWILD | 320864 | 1456 | 24102 | 70687 | 84.69 |
| 2090 | 45 | MIROC-ES2L | SEMIWILD | 331569 | 6083 | 13397 | 66060 | 87.15 |

**Supplementary table 5**

| | | | | | | | | |
|---|---|---|---|---|---|---|---|---|
| 2090 | 45 | MIROC6 | SEMIWILD | 324226 | 4219 | 20740 | 67924 | 84.48 |
| 2090 | 45 | MRI-ESM2-0 | SEMIWILD | 325957 | 3306 | 19009 | 68837 | 86.05 |
| 2090 | 85 | BCC-CSM2-MR | SEMIWILD | 312458 | 109 | 32508 | 72034 | 81.54 |
| 2090 | 85 | CNRM-CM6-1 | SEMIWILD | 320024 | 1975 | 24942 | 70168 | 83.91 |
| 2090 | 85 | CNRM-ESM2-1 | SEMIWILD | 322222 | 2431 | 22744 | 69712 | 84.71 |
| 2090 | 85 | CanESM5 | SEMIWILD | 293230 | 1318 | 51736 | 70825 | 72.75 |
| 2090 | 85 | IPSL-CM6A-LR | SEMIWILD | 312588 | 3299 | 32378 | 68844 | 79.42 |
| 2090 | 85 | MIROC-ES2L | SEMIWILD | 314962 | 8672 | 30004 | 63471 | 76.65 |
| 2090 | 85 | MIROC6 | SEMIWILD | 303708 | 4462 | 41258 | 67681 | 74.75 |
| 2090 | 85 | MRI-ESM2-0 | SEMIWILD | 316752 | 4999 | 28214 | 67144 | 80.17 |
| 2050 | 45 | BCC-CSM2-MR | COMMERCIAL | 330038 | 846 | 21703 | 64522 | 85.13 |
| 2050 | 45 | CNRM-CM6-1 | COMMERCIAL | 332763 | 1197 | 18978 | 64171 | 86.42 |
| 2050 | 45 | CNRM-ESM2-1 | COMMERCIAL | 327051 | 856 | 24690 | 64512 | 83.47 |
| 2050 | 45 | CanESM5 | COMMERCIAL | 326128 | 893 | 25613 | 64475 | 82.95 |
| 2050 | 45 | IPSL-CM6A-LR | COMMERCIAL | 331851 | 908 | 19890 | 64460 | 86.11 |
| 2050 | 45 | MIROC-ES2L | COMMERCIAL | 325691 | 401 | 26050 | 64967 | 83.09 |
| 2050 | 45 | MIROC6 | COMMERCIAL | 322699 | 335 | 29042 | 65033 | 81.58 |
| 2050 | 45 | MRI-ESM2-0 | COMMERCIAL | 331661 | 1052 | 20080 | 64316 | 85.89 |
| 2050 | 85 | BCC-CSM2-MR | COMMERCIAL | 335348 | 2273 | 16393 | 63095 | 87.11 |
| 2050 | 85 | CNRM-CM6-1 | COMMERCIAL | 328358 | 1253 | 23383 | 64115 | 83.88 |
| 2050 | 85 | CNRM-ESM2-1 | COMMERCIAL | 327704 | 1324 | 24037 | 64044 | 83.47 |
| 2050 | 85 | CanESM5 | COMMERCIAL | 325412 | 1605 | 26329 | 63763 | 82.03 |
| 2050 | 85 | IPSL-CM6A-LR | COMMERCIAL | 330111 | 1883 | 21630 | 63485 | 84.37 |
| 2050 | 85 | MIROC-ES2L | COMMERCIAL | 323557 | 1493 | 28184 | 63875 | 81.15 |
| 2050 | 85 | MIROC6 | COMMERCIAL | 319380 | 355 | 32361 | 65013 | 79.9 |
| 2050 | 85 | MRI-ESM2-0 | COMMERCIAL | 333618 | 2446 | 18123 | 62922 | 85.95 |
| 2070 | 45 | BCC-CSM2-MR | COMMERCIAL | 328909 | 2283 | 22832 | 63085 | 83.4 |
| 2070 | 45 | CNRM-CM6-1 | COMMERCIAL | 327593 | 1373 | 24148 | 63995 | 83.38 |
| 2070 | 45 | CNRM-ESM2-1 | COMMERCIAL | 326410 | 1533 | 25331 | 63835 | 82.62 |
| 2070 | 45 | CanESM5 | COMMERCIAL | 323659 | 1272 | 28082 | 64096 | 81.37 |
| 2070 | 45 | IPSL-CM6A-LR | COMMERCIAL | 332070 | 2280 | 19671 | 63088 | 85.18 |
| 2070 | 45 | MIROC-ES2L | COMMERCIAL | 326783 | 1695 | 24958 | 63673 | 82.69 |
| 2070 | 45 | MIROC6 | COMMERCIAL | 314803 | 1011 | 36938 | 64357 | 77.23 |
| 2070 | 45 | MRI-ESM2-0 | COMMERCIAL | 335127 | 2118 | 16614 | 63250 | 87.1 |
| 2070 | 85 | BCC-CSM2-MR | COMMERCIAL | 330681 | 4173 | 21060 | 61195 | 82.91 |
| 2070 | 85 | CNRM-CM6-1 | COMMERCIAL | 325442 | 2514 | 26299 | 62854 | 81.35 |
| 2070 | 85 | CNRM-ESM2-1 | COMMERCIAL | 325890 | 2632 | 25851 | 62736 | 81.5 |
| 2070 | 85 | CanESM5 | COMMERCIAL | 324011 | 3103 | 27730 | 62265 | 80.15 |
| 2070 | 85 | IPSL-CM6A-LR | COMMERCIAL | 326549 | 4325 | 25192 | 61043 | 80.53 |
| 2070 | 85 | MIROC-ES2L | COMMERCIAL | 317631 | 2135 | 34110 | 63233 | 77.72 |

**Supplementary table 5**

| | | | | | | | | |
|---|---|---|---|---|---|---|---|---|
| 2070 | 85 | MIROC6 | COMMERCIAL | 310303 | 2105 | 41438 | 63263 | 74.4 |
| 2070 | 85 | MRI-ESM2-0 | COMMERCIAL | 321306 | 1223 | 30435 | 64145 | 80.21 |
| 2090 | 45 | BCC-CSM2-MR | COMMERCIAL | 330878 | 1438 | 20863 | 63930 | 85.15 |
| 2090 | 45 | CNRM-CM6-1 | COMMERCIAL | 330607 | 2126 | 21134 | 63242 | 84.47 |
| 2090 | 45 | CNRM-ESM2-1 | COMMERCIAL | 323753 | 1552 | 27988 | 63816 | 81.21 |
| 2090 | 45 | CanESM5 | COMMERCIAL | 324338 | 1762 | 27403 | 63606 | 81.35 |
| 2090 | 45 | IPSL-CM6A-LR | COMMERCIAL | 327643 | 2260 | 24098 | 63108 | 82.72 |
| 2090 | 45 | MIROC-ES2L | COMMERCIAL | 323607 | 1626 | 28134 | 63742 | 81.07 |
| 2090 | 45 | MIROC6 | COMMERCIAL | 317269 | 710 | 34472 | 64658 | 78.61 |
| 2090 | 45 | MRI-ESM2-0 | COMMERCIAL | 331428 | 3156 | 20313 | 62212 | 84.13 |
| 2090 | 85 | BCC-CSM2-MR | COMMERCIAL | 328492 | 4594 | 23249 | 60774 | 81.36 |
| 2090 | 85 | CNRM-CM6-1 | COMMERCIAL | 325028 | 4240 | 26713 | 61128 | 79.8 |
| 2090 | 85 | CNRM-ESM2-1 | COMMERCIAL | 325075 | 4641 | 26666 | 60727 | 79.51 |
| 2090 | 85 | CanESM5 | COMMERCIAL | 324774 | 5173 | 26967 | 60195 | 78.93 |
| 2090 | 85 | IPSL-CM6A-LR | COMMERCIAL | 327197 | 6354 | 24544 | 59014 | 79.25 |
| 2090 | 85 | MIROC-ES2L | COMMERCIAL | 313060 | 2850 | 38681 | 62518 | 75.07 |
| 2090 | 85 | MIROC6 | COMMERCIAL | 304844 | 2387 | 46897 | 62981 | 71.88 |
| 2090 | 85 | MRI-ESM2-0 | COMMERCIAL | 316816 | 2038 | 34925 | 63330 | 77.41 |
| 2050 | 45 | BCC-CSM2-MR | LANDRACE | 357356 | 1224 | 12657 | 45872 | 86.86 |
| 2050 | 45 | CNRM-CM6-1 | LANDRACE | 355770 | 958 | 14243 | 46138 | 85.86 |
| 2050 | 45 | CNRM-ESM2-1 | LANDRACE | 357206 | 1198 | 12807 | 45898 | 86.76 |
| 2050 | 45 | CanESM5 | LANDRACE | 349176 | 494 | 20837 | 46602 | 81.38 |
| 2050 | 45 | IPSL-CM6A-LR | LANDRACE | 356487 | 1623 | 13526 | 45473 | 85.72 |
| 2050 | 45 | MIROC-ES2L | LANDRACE | 359888 | 4549 | 10125 | 42547 | 85.29 |
| 2050 | 45 | MIROC6 | LANDRACE | 357272 | 6070 | 12741 | 41026 | 81.35 |
| 2050 | 45 | MRI-ESM2-0 | LANDRACE | 354474 | 3720 | 15539 | 43376 | 81.83 |
| 2050 | 85 | BCC-CSM2-MR | LANDRACE | 355245 | 1031 | 14768 | 46065 | 85.36 |
| 2050 | 85 | CNRM-CM6-1 | LANDRACE | 356521 | 1191 | 13492 | 45905 | 86.21 |
| 2050 | 85 | CNRM-ESM2-1 | LANDRACE | 354651 | 823 | 15362 | 46273 | 85.11 |
| 2050 | 85 | CanESM5 | LANDRACE | 346363 | 670 | 23650 | 46426 | 79.24 |
| 2050 | 85 | IPSL-CM6A-LR | LANDRACE | 353592 | 2649 | 16421 | 44447 | 82.34 |
| 2050 | 85 | MIROC-ES2L | LANDRACE | 359256 | 6960 | 10757 | 40136 | 81.92 |
| 2050 | 85 | MIROC6 | LANDRACE | 356574 | 5873 | 13439 | 41223 | 81.02 |
| 2050 | 85 | MRI-ESM2-0 | LANDRACE | 357496 | 4136 | 12517 | 42960 | 83.76 |
| 2070 | 45 | BCC-CSM2-MR | LANDRACE | 353870 | 790 | 16143 | 46306 | 84.54 |
| 2070 | 45 | CNRM-CM6-1 | LANDRACE | 355445 | 1619 | 14568 | 45477 | 84.89 |
| 2070 | 45 | CNRM-ESM2-1 | LANDRACE | 355930 | 1576 | 14083 | 45520 | 85.32 |
| 2070 | 45 | CanESM5 | LANDRACE | 347259 | 633 | 22754 | 46463 | 79.89 |
| 2070 | 45 | IPSL-CM6A-LR | LANDRACE | 356271 | 2801 | 13742 | 44295 | 84.26 |
| 2070 | 45 | MIROC-ES2L | LANDRACE | 356780 | 7877 | 13233 | 39219 | 78.79 |

**Supplementary table 5**

| | | | | | | | | |
|---|---|---|---|---|---|---|---|---|
| 2070 | 45 | MIROC6 | LANDRACE | 357200 | 7488 | 12813 | 39608 | 79.6 |
| 2070 | 45 | MRI-ESM2-0 | LANDRACE | 353040 | 1706 | 16973 | 45390 | 82.94 |
| 2070 | 85 | BCC-CSM2-MR | LANDRACE | 350896 | 1397 | 19117 | 45699 | 81.67 |
| 2070 | 85 | CNRM-CM6-1 | LANDRACE | 352625 | 1495 | 17388 | 45601 | 82.85 |
| 2070 | 85 | CNRM-ESM2-1 | LANDRACE | 352174 | 1591 | 17839 | 45505 | 82.41 |
| 2070 | 85 | CanESM5 | LANDRACE | 343933 | 2216 | 26080 | 44880 | 76.03 |
| 2070 | 85 | IPSL-CM6A-LR | LANDRACE | 350574 | 2813 | 19439 | 44283 | 79.92 |
| 2070 | 85 | MIROC-ES2L | LANDRACE | 355638 | 8386 | 14375 | 38710 | 77.28 |
| 2070 | 85 | MIROC6 | LANDRACE | 350305 | 7806 | 19708 | 39290 | 74.07 |
| 2070 | 85 | MRI-ESM2-0 | LANDRACE | 351562 | 2530 | 18451 | 44566 | 80.95 |
| 2090 | 45 | BCC-CSM2-MR | LANDRACE | 355766 | 1739 | 14247 | 45357 | 85.02 |
| 2090 | 45 | CNRM-CM6-1 | LANDRACE | 356289 | 1462 | 13724 | 45634 | 85.73 |
| 2090 | 45 | CNRM-ESM2-1 | LANDRACE | 354244 | 1277 | 15769 | 45819 | 84.32 |
| 2090 | 45 | CanESM5 | LANDRACE | 346828 | 770 | 23185 | 46326 | 79.46 |
| 2090 | 45 | IPSL-CM6A-LR | LANDRACE | 354536 | 2583 | 15477 | 44513 | 83.14 |
| 2090 | 45 | MIROC-ES2L | LANDRACE | 357832 | 6132 | 12181 | 40964 | 81.73 |
| 2090 | 45 | MIROC6 | LANDRACE | 355669 | 5993 | 14344 | 41103 | 80.17 |
| 2090 | 45 | MRI-ESM2-0 | LANDRACE | 351232 | 1985 | 18781 | 45111 | 81.29 |
| 2090 | 85 | BCC-CSM2-MR | LANDRACE | 347878 | 2044 | 22135 | 45052 | 78.84 |
| 2090 | 85 | CNRM-CM6-1 | LANDRACE | 350693 | 2646 | 19320 | 44450 | 80.19 |
| 2090 | 85 | CNRM-ESM2-1 | LANDRACE | 350958 | 2270 | 19055 | 44826 | 80.78 |
| 2090 | 85 | CanESM5 | LANDRACE | 340647 | 4717 | 29366 | 42379 | 71.32 |
| 2090 | 85 | IPSL-CM6A-LR | LANDRACE | 349567 | 5223 | 20446 | 41873 | 76.54 |
| 2090 | 85 | MIROC-ES2L | LANDRACE | 352606 | 7782 | 17407 | 39314 | 75.74 |
| 2090 | 85 | MIROC6 | LANDRACE | 350538 | 8815 | 19475 | 38281 | 73.02 |
| 2090 | 85 | MRI-ESM2-0 | LANDRACE | 351899 | 4491 | 18114 | 42605 | 79.03 |
| 2050 | 45 | BCC-CSM2-MR | WILD | 327003 | 633 | 16875 | 72598 | 89.24 |
| 2050 | 45 | CNRM-CM6-1 | WILD | 326857 | 1623 | 17021 | 71608 | 88.48 |
| 2050 | 45 | CNRM-ESM2-1 | WILD | 325835 | 2350 | 18043 | 70881 | 87.42 |
| 2050 | 45 | CanESM5 | WILD | 309023 | 1683 | 34855 | 71548 | 79.66 |
| 2050 | 45 | IPSL-CM6A-LR | WILD | 325172 | 3768 | 18706 | 69463 | 86.08 |
| 2050 | 45 | MIROC-ES2L | WILD | 326200 | 3638 | 17678 | 69593 | 86.72 |
| 2050 | 45 | MIROC6 | WILD | 323345 | 3394 | 20533 | 69837 | 85.37 |
| 2050 | 45 | MRI-ESM2-0 | WILD | 318045 | 4448 | 25833 | 68783 | 81.96 |
| 2050 | 85 | BCC-CSM2-MR | WILD | 324188 | 302 | 19690 | 72929 | 87.95 |
| 2050 | 85 | CNRM-CM6-1 | WILD | 325675 | 1821 | 18203 | 71410 | 87.7 |
| 2050 | 85 | CNRM-ESM2-1 | WILD | 325143 | 2053 | 18735 | 71178 | 87.26 |
| 2050 | 85 | CanESM5 | WILD | 302572 | 2186 | 41306 | 71045 | 76.56 |
| 2050 | 85 | IPSL-CM6A-LR | WILD | 321680 | 3629 | 22198 | 69602 | 84.35 |
| 2050 | 85 | MIROC-ES2L | WILD | 322789 | 4175 | 21089 | 69056 | 84.54 |

**Supplementary table 5**

| | | | | | | | | |
|---|---|---|---|---|---|---|---|---|
| 2050 | 85 | MIROC6 | WILD | 321140 | 4868 | 22738 | 68363 | 83.2 |
| 2050 | 85 | MRI-ESM2-0 | WILD | 322085 | 3081 | 21793 | 70150 | 84.94 |
| 2070 | 45 | BCC-CSM2-MR | WILD | 324618 | 1114 | 19260 | 72117 | 87.62 |
| 2070 | 45 | CNRM-CM6-1 | WILD | 325937 | 2914 | 17941 | 70317 | 87.09 |
| 2070 | 45 | CNRM-ESM2-1 | WILD | 321212 | 2687 | 22666 | 70544 | 84.77 |
| 2070 | 45 | CanESM5 | WILD | 304033 | 2149 | 39845 | 71082 | 77.2 |
| 2070 | 45 | IPSL-CM6A-LR | WILD | 322421 | 3164 | 21457 | 70067 | 85.06 |
| 2070 | 45 | MIROC-ES2L | WILD | 322553 | 4448 | 21325 | 68783 | 84.22 |
| 2070 | 45 | MIROC6 | WILD | 318709 | 6074 | 25169 | 67157 | 81.13 |
| 2070 | 45 | MRI-ESM2-0 | WILD | 316465 | 2233 | 27413 | 70998 | 82.73 |
| 2070 | 85 | BCC-CSM2-MR | WILD | 313559 | 885 | 30319 | 72346 | 82.26 |
| 2070 | 85 | CNRM-CM6-1 | WILD | 317491 | 2504 | 26387 | 70727 | 83.04 |
| 2070 | 85 | CNRM-ESM2-1 | WILD | 316264 | 2489 | 27614 | 70742 | 82.46 |
| 2070 | 85 | CanESM5 | WILD | 292043 | 4147 | 51835 | 69084 | 71.17 |
| 2070 | 85 | IPSL-CM6A-LR | WILD | 310358 | 4080 | 33520 | 69151 | 78.62 |
| 2070 | 85 | MIROC-ES2L | WILD | 308128 | 6777 | 35750 | 66454 | 75.76 |
| 2070 | 85 | MIROC6 | WILD | 307117 | 5655 | 36761 | 67576 | 76.11 |
| 2070 | 85 | MRI-ESM2-0 | WILD | 308478 | 4230 | 35400 | 69001 | 77.69 |
| 2090 | 45 | BCC-CSM2-MR | WILD | 324074 | 1202 | 19804 | 72029 | 87.27 |
| 2090 | 45 | CNRM-CM6-1 | WILD | 323482 | 3003 | 20396 | 70228 | 85.72 |
| 2090 | 45 | CNRM-ESM2-1 | WILD | 318610 | 2428 | 25268 | 70803 | 83.64 |
| 2090 | 45 | CanESM5 | WILD | 301930 | 1938 | 41948 | 71293 | 76.47 |
| 2090 | 45 | IPSL-CM6A-LR | WILD | 316442 | 3217 | 27436 | 70014 | 82.04 |
| 2090 | 45 | MIROC-ES2L | WILD | 322002 | 5511 | 21876 | 67720 | 83.18 |
| 2090 | 45 | MIROC6 | WILD | 318631 | 3451 | 25247 | 69780 | 82.94 |
| 2090 | 45 | MRI-ESM2-0 | WILD | 318472 | 3333 | 25406 | 69898 | 82.95 |
| 2090 | 85 | BCC-CSM2-MR | WILD | 310235 | 1549 | 33643 | 71682 | 80.29 |
| 2090 | 85 | CNRM-CM6-1 | WILD | 307906 | 4013 | 35972 | 69218 | 77.59 |
| 2090 | 85 | CNRM-ESM2-1 | WILD | 308681 | 3191 | 35197 | 70040 | 78.49 |
| 2090 | 85 | CanESM5 | WILD | 284775 | 7764 | 59103 | 65467 | 66.19 |
| 2090 | 85 | IPSL-CM6A-LR | WILD | 302631 | 5462 | 41247 | 67769 | 74.37 |
| 2090 | 85 | MIROC-ES2L | WILD | 293934 | 4281 | 49944 | 68950 | 71.78 |
| 2090 | 85 | MIROC6 | WILD | 287872 | 6935 | 56006 | 66296 | 67.81 |
| 2090 | 85 | MRI-ESM2-0 | WILD | 299067 | 5336 | 44811 | 67895 | 73.03 |
| 2050 | 45 | BCC-CSM2-MR | CULTIVATED | 323250 | 762 | 22449 | 70648 | 85.89 |
| 2050 | 45 | CNRM-CM6-1 | CULTIVATED | 323360 | 751 | 22339 | 70659 | 85.96 |
| 2050 | 45 | CNRM-ESM2-1 | CULTIVATED | 318460 | 520 | 27239 | 70890 | 83.63 |
| 2050 | 45 | CanESM5 | CULTIVATED | 317207 | 431 | 28492 | 70979 | 83.07 |
| 2050 | 45 | IPSL-CM6A-LR | CULTIVATED | 323177 | 743 | 22522 | 70667 | 85.87 |
| 2050 | 45 | MIROC-ES2L | CULTIVATED | 316677 | 289 | 29022 | 71121 | 82.91 |

**Supplementary table 5**

| | | | | | | | | |
|---|---|---|---|---|---|---|---|---|
| 2050 | 45 | MIROC6 | CULTIVATED | 315317 | 378 | 30382 | 71032 | 82.2 |
| 2050 | 45 | MRI-ESM2-0 | CULTIVATED | 323030 | 874 | 22669 | 70536 | 85.7 |
| 2050 | 85 | BCC-CSM2-MR | CULTIVATED | 326476 | 2155 | 19223 | 69255 | 86.63 |
| 2050 | 85 | CNRM-CM6-1 | CULTIVATED | 319593 | 807 | 26106 | 70603 | 83.99 |
| 2050 | 85 | CNRM-ESM2-1 | CULTIVATED | 317832 | 816 | 27867 | 70594 | 83.11 |
| 2050 | 85 | CanESM5 | CULTIVATED | 315680 | 690 | 30019 | 70720 | 82.16 |
| 2050 | 85 | IPSL-CM6A-LR | CULTIVATED | 320909 | 1127 | 24790 | 70283 | 84.43 |
| 2050 | 85 | MIROC-ES2L | CULTIVATED | 314693 | 809 | 31006 | 70601 | 81.61 |
| 2050 | 85 | MIROC6 | CULTIVATED | 312145 | 416 | 33554 | 70994 | 80.69 |
| 2050 | 85 | MRI-ESM2-0 | CULTIVATED | 324746 | 2214 | 20953 | 69196 | 85.66 |
| 2070 | 45 | BCC-CSM2-MR | CULTIVATED | 321881 | 3178 | 23818 | 68232 | 83.48 |
| 2070 | 45 | CNRM-CM6-1 | CULTIVATED | 318412 | 1014 | 27287 | 70396 | 83.26 |
| 2070 | 45 | CNRM-ESM2-1 | CULTIVATED | 316753 | 957 | 28946 | 70453 | 82.49 |
| 2070 | 45 | CanESM5 | CULTIVATED | 314301 | 422 | 31398 | 70988 | 81.69 |
| 2070 | 45 | IPSL-CM6A-LR | CULTIVATED | 323747 | 2203 | 21952 | 69207 | 85.14 |
| 2070 | 45 | MIROC-ES2L | CULTIVATED | 316644 | 899 | 29055 | 70511 | 82.48 |
| 2070 | 45 | MIROC6 | CULTIVATED | 307082 | 533 | 38617 | 70877 | 78.36 |
| 2070 | 45 | MRI-ESM2-0 | CULTIVATED | 325354 | 1621 | 20345 | 69789 | 86.4 |
| 2070 | 85 | BCC-CSM2-MR | CULTIVATED | 322340 | 4060 | 23359 | 67350 | 83.09 |
| 2070 | 85 | CNRM-CM6-1 | CULTIVATED | 314161 | 2069 | 31538 | 69341 | 80.49 |
| 2070 | 85 | CNRM-ESM2-1 | CULTIVATED | 315389 | 2101 | 30310 | 69309 | 81.05 |
| 2070 | 85 | CanESM5 | CULTIVATED | 313883 | 2087 | 31816 | 69323 | 80.35 |
| 2070 | 85 | IPSL-CM6A-LR | CULTIVATED | 318010 | 3974 | 27689 | 67436 | 80.99 |
| 2070 | 85 | MIROC-ES2L | CULTIVATED | 307568 | 1270 | 38131 | 70140 | 78.07 |
| 2070 | 85 | MIROC6 | CULTIVATED | 302725 | 1490 | 42974 | 69920 | 75.87 |
| 2070 | 85 | MRI-ESM2-0 | CULTIVATED | 312780 | 477 | 32919 | 70933 | 80.95 |
| 2090 | 45 | BCC-CSM2-MR | CULTIVATED | 323078 | 2342 | 22621 | 69068 | 84.69 |
| 2090 | 45 | CNRM-CM6-1 | CULTIVATED | 321075 | 1619 | 24624 | 69791 | 84.17 |
| 2090 | 45 | CNRM-ESM2-1 | CULTIVATED | 313493 | 940 | 32206 | 70470 | 80.96 |
| 2090 | 45 | CanESM5 | CULTIVATED | 315003 | 716 | 30696 | 70694 | 81.82 |
| 2090 | 45 | IPSL-CM6A-LR | CULTIVATED | 319853 | 1653 | 25846 | 69757 | 83.53 |
| 2090 | 45 | MIROC-ES2L | CULTIVATED | 316648 | 1325 | 29051 | 70085 | 82.19 |
| 2090 | 45 | MIROC6 | CULTIVATED | 309379 | 391 | 36320 | 71019 | 79.46 |
| 2090 | 45 | MRI-ESM2-0 | CULTIVATED | 322144 | 2372 | 23555 | 69038 | 84.19 |
| 2090 | 85 | BCC-CSM2-MR | CULTIVATED | 319898 | 5030 | 25801 | 66380 | 81.15 |
| 2090 | 85 | CNRM-CM6-1 | CULTIVATED | 313307 | 4008 | 32392 | 67402 | 78.74 |
| 2090 | 85 | CNRM-ESM2-1 | CULTIVATED | 313434 | 4056 | 32265 | 67354 | 78.76 |
| 2090 | 85 | CanESM5 | CULTIVATED | 313381 | 4575 | 32318 | 66835 | 78.37 |
| 2090 | 85 | IPSL-CM6A-LR | CULTIVATED | 316801 | 6335 | 28898 | 65075 | 78.7 |
| 2090 | 85 | MIROC-ES2L | CULTIVATED | 304764 | 1322 | 40935 | 70088 | 76.84 |

**Supplementary table 5**

| | | | | | | | | |
|---|---|---|---|---|---|---|---|---|
| 2090 | 85 | MIROC6 | CULTIVATED | 295694 | 1440 | 50005 | 69970 | 73.12 |
| 2090 | 85 | MRI-ESM2-0 | CULTIVATED | 308178 | 1772 | 37521 | 69638 | 78 |
| 2050 | 45 | BCC-CSM2-MR | WILDsl | 326345 | 1023 | 17400 | 72341 | 88.7 |
| 2050 | 45 | CNRM-CM6-1 | WILDsl | 325472 | 1446 | 18273 | 71918 | 87.94 |
| 2050 | 45 | CNRM-ESM2-1 | WILDsl | 326017 | 2504 | 17728 | 70860 | 87.51 |
| 2050 | 45 | CanESM5 | WILDsl | 309819 | 1862 | 33926 | 71502 | 79.98 |
| 2050 | 45 | IPSL-CM6A-LR | WILDsl | 323599 | 3911 | 20146 | 69453 | 85.24 |
| 2050 | 45 | MIROC-ES2L | WILDsl | 325969 | 3517 | 17776 | 69847 | 86.77 |
| 2050 | 45 | MIROC6 | WILDsl | 324228 | 3327 | 19517 | 70037 | 85.98 |
| 2050 | 45 | MRI-ESM2-0 | WILDsl | 318885 | 5060 | 24860 | 68304 | 82.03 |
| 2050 | 85 | BCC-CSM2-MR | WILDsl | 321613 | 270 | 22132 | 73094 | 86.71 |
| 2050 | 85 | CNRM-CM6-1 | WILDsl | 324406 | 1807 | 19339 | 71557 | 87.13 |
| 2050 | 85 | CNRM-ESM2-1 | WILDsl | 324470 | 2070 | 19275 | 71294 | 86.98 |
| 2050 | 85 | CanESM5 | WILDsl | 303764 | 2397 | 39981 | 70967 | 77.01 |
| 2050 | 85 | IPSL-CM6A-LR | WILDsl | 319162 | 3260 | 24583 | 70104 | 83.43 |
| 2050 | 85 | MIROC-ES2L | WILDsl | 322684 | 4509 | 21061 | 68855 | 84.34 |
| 2050 | 85 | MIROC6 | WILDsl | 320659 | 5210 | 23086 | 68154 | 82.81 |
| 2050 | 85 | MRI-ESM2-0 | WILDsl | 322619 | 3464 | 21126 | 69900 | 85.04 |
| 2070 | 45 | BCC-CSM2-MR | WILDsl | 322962 | 1724 | 20783 | 71640 | 86.42 |
| 2070 | 45 | CNRM-CM6-1 | WILDsl | 324599 | 2760 | 19146 | 70604 | 86.57 |
| 2070 | 45 | CNRM-ESM2-1 | WILDsl | 320975 | 2893 | 22770 | 70471 | 84.6 |
| 2070 | 45 | CanESM5 | WILDsl | 305232 | 2377 | 38513 | 70987 | 77.64 |
| 2070 | 45 | IPSL-CM6A-LR | WILDsl | 321341 | 2604 | 22404 | 70760 | 84.98 |
| 2070 | 45 | MIROC-ES2L | WILDsl | 323422 | 4940 | 20323 | 68424 | 84.42 |
| 2070 | 45 | MIROC6 | WILDsl | 319352 | 6032 | 24393 | 67332 | 81.57 |
| 2070 | 45 | MRI-ESM2-0 | WILDsl | 317757 | 2572 | 25988 | 70792 | 83.21 |
| 2070 | 85 | BCC-CSM2-MR | WILDsl | 309566 | 1201 | 34179 | 72163 | 80.31 |
| 2070 | 85 | CNRM-CM6-1 | WILDsl | 318056 | 2664 | 25689 | 70700 | 83.3 |
| 2070 | 85 | CNRM-ESM2-1 | WILDsl | 316335 | 2673 | 27410 | 70691 | 82.46 |
| 2070 | 85 | CanESM5 | WILDsl | 292203 | 3371 | 51542 | 69993 | 71.82 |
| 2070 | 85 | IPSL-CM6A-LR | WILDsl | 310306 | 3593 | 33439 | 69771 | 79.03 |
| 2070 | 85 | MIROC-ES2L | WILDsl | 312297 | 7935 | 31448 | 65429 | 76.87 |
| 2070 | 85 | MIROC6 | WILDsl | 307203 | 6499 | 36542 | 66865 | 75.65 |
| 2070 | 85 | MRI-ESM2-0 | WILDsl | 309764 | 4848 | 33981 | 68516 | 77.92 |
| 2090 | 45 | BCC-CSM2-MR | WILDsl | 321087 | 1511 | 22658 | 71853 | 85.6 |
| 2090 | 45 | CNRM-CM6-1 | WILDsl | 322080 | 2860 | 21665 | 70504 | 85.18 |
| 2090 | 45 | CNRM-ESM2-1 | WILDsl | 318714 | 2704 | 25031 | 70660 | 83.59 |
| 2090 | 45 | CanESM5 | WILDsl | 303451 | 2188 | 40294 | 71176 | 77.02 |
| 2090 | 45 | IPSL-CM6A-LR | WILDsl | 315461 | 2735 | 28284 | 70629 | 81.99 |
| 2090 | 45 | MIROC-ES2L | WILDsl | 323595 | 5656 | 20150 | 67708 | 83.99 |

**Supplementary table 5**

| | | | | | | | | |
|---|---|---|---|---|---|---|---|---|
| 2090 | 45 | MIROC6 | WILDsl | 317802 | 3767 | 25943 | 69597 | 82.41 |
| 2090 | 45 | MRI-ESM2-0 | WILDsl | 318249 | 3392 | 25496 | 69972 | 82.89 |
| 2090 | 85 | BCC-CSM2-MR | WILDsl | 305489 | 1573 | 38256 | 71791 | 78.28 |
| 2090 | 85 | CNRM-CM6-1 | WILDsl | 311270 | 4073 | 32475 | 69291 | 79.13 |
| 2090 | 85 | CNRM-ESM2-1 | WILDsl | 311251 | 3603 | 32494 | 69761 | 79.45 |
| 2090 | 85 | CanESM5 | WILDsl | 285499 | 5298 | 58246 | 68066 | 68.18 |
| 2090 | 85 | IPSL-CM6A-LR | WILDsl | 306255 | 5458 | 37490 | 67906 | 75.97 |
| 2090 | 85 | MIROC-ES2L | WILDsl | 298376 | 5555 | 45369 | 67809 | 72.7 |
| 2090 | 85 | MIROC6 | WILDsl | 290236 | 8022 | 53509 | 65342 | 67.99 |
| 2090 | 85 | MRI-ESM2-0 | WILDsl | 303956 | 5963 | 39789 | 67401 | 74.66 |

### GCM median

| year | ssp | GCM | Dom | none | pixLost | pixNew | pixKeep | percent |
|---|---|---|---|---|---|---|---|---|
| 2050 | 45 | Median | SEMIWILD | 329674 | 1227 | 15292 | 70916 | 89.57 |
| 2050 | 85 | Median | SEMIWILD | 328349 | 980 | 16617 | 71163 | 89 |
| 2070 | 45 | Median | SEMIWILD | 327492 | 1241 | 17474 | 70902 | 88.34 |
| 2070 | 85 | Median | SEMIWILD | 320410 | 799 | 24556 | 71344 | 84.91 |
| 2090 | 45 | Median | SEMIWILD | 327123 | 874 | 17843 | 71269 | 88.39 |
| 2090 | 85 | Median | SEMIWILD | 313900 | 1166 | 31066 | 70977 | 81.5 |
| 2050 | 45 | Median | COMMERCIAL | 329286 | 372 | 22455 | 64996 | 85.06 |
| 2050 | 85 | Median | COMMERCIAL | 328719 | 1121 | 23022 | 64247 | 84.18 |
| 2070 | 45 | Median | COMMERCIAL | 328327 | 1325 | 23414 | 64043 | 83.81 |
| 2070 | 85 | Median | COMMERCIAL | 324787 | 2259 | 26954 | 63109 | 81.21 |
| 2090 | 45 | Median | COMMERCIAL | 327051 | 1397 | 24690 | 63971 | 83.06 |
| 2090 | 85 | Median | COMMERCIAL | 323286 | 2869 | 28455 | 62499 | 79.96 |
| 2050 | 45 | Median | LANDRACE | 357115 | 1055 | 12898 | 46041 | 86.84 |
| 2050 | 85 | Median | LANDRACE | 356499 | 1315 | 13514 | 45781 | 86.06 |
| 2070 | 45 | Median | LANDRACE | 355963 | 1160 | 14050 | 45936 | 85.8 |
| 2070 | 85 | Median | LANDRACE | 353484 | 1122 | 16529 | 45974 | 83.89 |
| 2090 | 45 | Median | LANDRACE | 356260 | 1135 | 13753 | 45961 | 86.06 |
| 2090 | 85 | Median | LANDRACE | 352102 | 1924 | 17911 | 45172 | 82 |
| 2050 | 45 | Median | WILD | 324800 | 1398 | 19078 | 71833 | 87.53 |
| 2050 | 85 | Median | WILD | 323222 | 1421 | 20656 | 71810 | 86.68 |
| 2070 | 45 | Median | WILD | 322962 | 1978 | 20916 | 71253 | 86.16 |
| 2070 | 85 | Median | WILD | 314002 | 1825 | 29876 | 71406 | 81.83 |
| 2090 | 45 | Median | WILD | 321168 | 1639 | 22710 | 71592 | 85.47 |
| 2090 | 85 | Median | WILD | 304283 | 2065 | 39595 | 71166 | 77.36 |
| 2050 | 45 | Median | CULTIVATED | 320583 | 125 | 25116 | 71285 | 84.96 |
| 2050 | 85 | Median | CULTIVATED | 319648 | 489 | 26051 | 70921 | 84.24 |
| 2070 | 45 | Median | CULTIVATED | 319182 | 709 | 26517 | 70701 | 83.85 |

**Supplementary table 5**

| | | | | | | | | |
|---|---|---|---|---|---|---|---|---|
| 2070 | 85 | Median | CULTIVATED | 315294 | 1468 | 30405 | 69942 | 81.44 |
| 2090 | 45 | Median | CULTIVATED | 318725 | 754 | 26974 | 70656 | 83.6 |
| 2090 | 85 | Median | CULTIVATED | 312817 | 2304 | 32882 | 69106 | 79.71 |
| 2050 | 45 | Median | WILDsl | 324102 | 1415 | 19643 | 71949 | 87.23 |
| 2050 | 85 | Median | WILDsl | 321692 | 1446 | 22053 | 71918 | 85.96 |
| 2070 | 45 | Median | WILDsl | 321947 | 2007 | 21798 | 71357 | 85.7 |
| 2070 | 85 | Median | WILDsl | 312706 | 2086 | 31039 | 71278 | 81.14 |
| 2090 | 45 | Median | WILDsl | 319582 | 1716 | 24163 | 71648 | 84.7 |
| 2090 | 85 | Median | WILDsl | 306030 | 2500 | 37715 | 70864 | 77.9 |

### GCM Intersecion

| year | ssp | GCM | Dom | none | pixLost | pixNew | pixKeep | percent |
|---|---|---|---|---|---|---|---|---|
| 2050 | 45 | Interseciton | SEMIWILD | 338278 | 9078 | 6688 | 63065 | 88.89 |
| 2050 | 85 | Interseciton | SEMIWILD | 337680 | 11017 | 7286 | 61126 | 86.98 |
| 2070 | 45 | Interseciton | SEMIWILD | 337511 | 10449 | 7455 | 61694 | 87.33 |
| 2070 | 85 | Interseciton | SEMIWILD | 335865 | 13431 | 9101 | 58712 | 83.9 |
| 2090 | 45 | Interseciton | SEMIWILD | 337137 | 9721 | 7829 | 62422 | 87.68 |
| 2090 | 85 | Interseciton | SEMIWILD | 334195 | 13425 | 10771 | 58718 | 82.92 |
| 2050 | 45 | Interseciton | COMMERCIAL | 342337 | 3289 | 9404 | 62079 | 90.72 |
| 2050 | 85 | Interseciton | COMMERCIAL | 342483 | 4632 | 9258 | 60736 | 89.74 |
| 2070 | 45 | Interseciton | COMMERCIAL | 341895 | 4878 | 9846 | 60490 | 89.15 |
| 2070 | 85 | Interseciton | COMMERCIAL | 340523 | 7284 | 11218 | 58084 | 86.26 |
| 2090 | 45 | Interseciton | COMMERCIAL | 341923 | 5817 | 9818 | 59551 | 88.4 |
| 2090 | 85 | Interseciton | COMMERCIAL | 339595 | 10419 | 12146 | 54949 | 82.97 |
| 2050 | 45 | Interseciton | LANDRACE | 365736 | 9403 | 4277 | 37693 | 84.64 |
| 2050 | 85 | Interseciton | LANDRACE | 365240 | 10131 | 4773 | 36965 | 83.22 |
| 2070 | 45 | Interseciton | LANDRACE | 365883 | 11071 | 4130 | 36025 | 82.58 |
| 2070 | 85 | Interseciton | LANDRACE | 365415 | 12689 | 4598 | 34407 | 79.92 |
| 2090 | 45 | Interseciton | LANDRACE | 365423 | 9684 | 4590 | 37412 | 83.98 |
| 2090 | 85 | Interseciton | LANDRACE | 364449 | 16887 | 5564 | 30209 | 72.91 |
| 2050 | 45 | Interseciton | WILD | 338013 | 8275 | 5865 | 64956 | 90.18 |
| 2050 | 85 | Interseciton | WILD | 337440 | 9587 | 6438 | 63644 | 88.82 |
| 2070 | 45 | Interseciton | WILD | 338179 | 9906 | 5699 | 63325 | 89.03 |
| 2070 | 85 | Interseciton | WILD | 335206 | 13760 | 8672 | 59471 | 84.13 |
| 2090 | 45 | Interseciton | WILD | 337170 | 9844 | 6708 | 63387 | 88.45 |
| 2090 | 85 | Interseciton | WILD | 333646 | 18814 | 10232 | 54417 | 78.93 |
| 2050 | 45 | Interseciton | CULTIVATED | 334765 | 2948 | 10934 | 68462 | 90.79 |
| 2050 | 85 | Interseciton | CULTIVATED | 335648 | 4384 | 10051 | 67026 | 90.28 |
| 2070 | 45 | Interseciton | CULTIVATED | 334878 | 5027 | 10821 | 66383 | 89.34 |
| 2070 | 85 | Interseciton | CULTIVATED | 334290 | 7121 | 11409 | 64289 | 87.4 |

**Supplementary table 5**

| | | | | | | | | |
|---|---|---|---|---|---|---|---|---|
| 2090 | 45 | Interseciton | CULTIVATED | 334024 | 5237 | 11675 | 66173 | 88.67 |
| 2090 | 85 | Interseciton | CULTIVATED | 333070 | 10247 | 12629 | 61163 | 84.25 |
| 2050 | 45 | Interseciton | WILDsl | 336917 | 8926 | 6828 | 64438 | 89.11 |
| 2050 | 85 | Interseciton | WILDsl | 336763 | 9520 | 6982 | 63844 | 88.56 |
| 2070 | 45 | Interseciton | WILDsl | 337202 | 10569 | 6543 | 62795 | 88.01 |
| 2070 | 85 | Interseciton | WILDsl | 334754 | 13749 | 8991 | 59615 | 83.98 |
| 2090 | 45 | Interseciton | WILDsl | 335949 | 9930 | 7796 | 63434 | 87.74 |
| 2090 | 85 | Interseciton | WILDsl | 333484 | 17329 | 10261 | 56035 | 80.24 |

### Supplementary table 6

| year | ssp | FirstInPair | SecondInPair | none | only_FirstInPair | only_SecondInPair | overlap | percent |
|---|---|---|---|---|---|---|---|---|
| 2050 | 45 | ARVENSE | COMMERCIAL | 306379 | 39247 | 40977 | 30506 | 43.20 |
| 2050 | 85 | ARVENSE | COMMERCIAL | 308610 | 38505 | 40087 | 29907 | 43.22 |
| 2070 | 45 | ARVENSE | COMMERCIAL | 308709 | 38064 | 39251 | 31085 | 44.57 |
| 2070 | 85 | ARVENSE | COMMERCIAL | 310142 | 37665 | 39154 | 30148 | 43.97 |
| 2090 | 45 | ARVENSE | COMMERCIAL | 307880 | 39860 | 38978 | 30391 | 43.53 |
| 2090 | 85 | ARVENSE | COMMERCIAL | 310403 | 39611 | 37217 | 29878 | 43.75 |
| 2050 | 45 | ARVENSE | LANDRACE | 346047 | 29092 | 1309 | 40661 | 72.79 |
| 2050 | 85 | ARVENSE | LANDRACE | 347507 | 27864 | 1190 | 40548 | 73.62 |
| 2070 | 45 | ARVENSE | LANDRACE | 346895 | 30059 | 1065 | 39090 | 71.53 |
| 2070 | 85 | ARVENSE | LANDRACE | 348752 | 29352 | 544 | 38461 | 72.01 |
| 2090 | 45 | ARVENSE | LANDRACE | 346055 | 29052 | 803 | 41199 | 73.40 |
| 2090 | 85 | ARVENSE | LANDRACE | 347277 | 34059 | 343 | 35430 | 67.32 |
| 2050 | 45 | ARVENSE | WILD | 328060 | 18228 | 19296 | 51525 | 73.31 |
| 2050 | 85 | ARVENSE | WILD | 328557 | 18470 | 20140 | 49942 | 72.12 |
| 2070 | 45 | ARVENSE | WILD | 329096 | 18989 | 18864 | 50160 | 72.60 |
| 2070 | 85 | ARVENSE | WILD | 329101 | 19865 | 20195 | 47948 | 70.53 |
| 2090 | 45 | ARVENSE | WILD | 328873 | 18141 | 17985 | 52110 | 74.26 |
| 2090 | 85 | ARVENSE | WILD | 326961 | 25499 | 20659 | 43990 | 65.59 |
| 2050 | 45 | COMMERCIAL | LANDRACE | 325798 | 49341 | 19828 | 22142 | 39.03 |
| 2050 | 85 | COMMERCIAL | LANDRACE | 327055 | 48316 | 20060 | 21678 | 38.80 |
| 2070 | 45 | COMMERCIAL | LANDRACE | 329076 | 47878 | 17697 | 22458 | 40.65 |
| 2070 | 85 | COMMERCIAL | LANDRACE | 327948 | 50156 | 19859 | 19146 | 35.36 |
| 2090 | 45 | COMMERCIAL | LANDRACE | 327087 | 48020 | 20653 | 21349 | 38.34 |
| 2090 | 85 | COMMERCIAL | LANDRACE | 331312 | 50024 | 18702 | 17071 | 33.19 |
| 2050 | 45 | COMMERCIAL | WILD | 308427 | 37861 | 37199 | 33622 | 47.25 |
| 2050 | 85 | COMMERCIAL | WILD | 310128 | 36899 | 36987 | 33095 | 47.25 |
| 2070 | 45 | COMMERCIAL | WILD | 311088 | 36997 | 35685 | 33339 | 47.85 |
| 2070 | 85 | COMMERCIAL | WILD | 312342 | 36624 | 35465 | 32678 | 47.55 |
| 2090 | 45 | COMMERCIAL | WILD | 311494 | 35520 | 36246 | 33849 | 48.54 |
| 2090 | 85 | COMMERCIAL | WILD | 316540 | 35920 | 33474 | 31175 | 47.33 |
| 2050 | 45 | LANDRACE | WILD | 339952 | 6336 | 35187 | 35634 | 63.19 |
| 2050 | 85 | LANDRACE | WILD | 340211 | 6816 | 35160 | 34922 | 62.46 |
| 2070 | 45 | LANDRACE | WILD | 341298 | 6787 | 35656 | 33368 | 61.13 |
| 2070 | 85 | LANDRACE | WILD | 342964 | 6002 | 35140 | 33003 | 61.60 |
| 2090 | 45 | LANDRACE | WILD | 341582 | 5432 | 33525 | 36570 | 65.25 |
| 2090 | 85 | LANDRACE | WILD | 346364 | 6096 | 34972 | 29677 | 59.10 |
| 2050 | 45 | CULTIVATED | WILDsl | 306051 | 39792 | 31662 | 39604 | 52.57 |
| 2050 | 85 | CULTIVATED | WILDsl | 308168 | 38115 | 31864 | 38962 | 52.69 |
| 2070 | 45 | CULTIVATED | WILDsl | 308650 | 39121 | 31255 | 38083 | 51.98 |
| 2070 | 85 | CULTIVATED | WILDsl | 309870 | 38633 | 31541 | 37065 | 51.37 |
| 2090 | 45 | CULTIVATED | WILDsl | 308387 | 37492 | 30874 | 40356 | 54.14 |
| 2090 | 85 | CULTIVATED | WILDsl | 312699 | 38114 | 30618 | 35678 | 50.94 |

**Supplementary table 7**

| comparisonData1 | comparisonData2 | D | I | rank.cor |
|---|---|---|---|---|
| SEMIWILD | COMMERCIAL | 0.526 | 0.797 | 0.755 |
| SEMIWILD | LANDRACE | 0.793 | 0.951 | 0.874 |
| SEMIWILD | WILD | 0.718 | 0.926 | 0.928 |
| SEMIWILD | CULTIVATED | 0.570 | 0.829 | 0.785 |
| SEMIWILD | WILDsl | 0.752 | 0.939 | 0.939 |
| COMMERCIAL | LANDRACE | 0.487 | 0.746 | 0.491 |
| COMMERCIAL | WILD | 0.611 | 0.855 | 0.760 |
| COMMERCIAL | CULTIVATED | 0.933 | 0.995 | 0.995 |
| COMMERCIAL | WILDsl | 0.603 | 0.851 | 0.759 |
| LANDRACE | WILD | 0.683 | 0.892 | 0.800 |
| LANDRACE | CULTIVATED | 0.526 | 0.775 | 0.526 |
| LANDRACE | WILDsl | 0.714 | 0.908 | 0.829 |
| WILD | CULTIVATED | 0.660 | 0.885 | 0.792 |
| WILD | WILDsl | 0.946 | 0.997 | 0.990 |

### Supplementary table A2

| category | pixels |
|---|---|
| **SEMIWILD-COMMERCIAL** | |
| none | 305216 |
| only_SEMIWILD | 46525 |
| only_COMMERCIAL | 39750 |
| overlap | 25618 |
| percent | 37.26 |
| **WILD-LANDRACE** | |
| none | 337510 |
| only_LANDRACE | 6368 |
| only_WILD | 32503 |
| overlap | 40728 |
| percent | 67.70 |
| **WILD_SL-CULTIVATED** | |
| none | 310630 |
| only_CULTIVATED | 33115 |
| only_WILDsl | 35069 |
| overlap | 38295 |
| percent | 52.90 |
| **WILD-COMMERCIAL** | |
| none | 311218 |
| only_COMMERCIAL | 32660 |
| only_WILD | 40523 |
| overlap | 32708 |
| percent | 47.20 |
| **LANDRACE-COMMERCIAL** | |
| none | 324775 |
| only_COMMERCIAL | 45238 |
| only_LANDRACE | 26966 |
| overlap | 20130 |
| percent | 35.80 |
| **WILD-SEMIWILD** | |
| none | 325209 |
| only_SEMIWILD | 18669 |
| only_WILD | 19757 |
| overlap | 53474 |
| percent | 73.57 |
| **SEMIWILD-LANDRACE** | |
| none | 342193 |
| only_SEMIWILD | 27820 |
| only_LANDRACE | 2773 |
| overlap | 44323 |
| percent | 74.34 |
