## Appendix_1 for "Capturing the distribution as it shifts: chile pepper (*Capsicum annuum* L.) domestication gradient meets geography"

(Methods details)

### PCA

#### Data imputation

Exceptionally, values were imputed for some point-variable combinations that had missing data, using the multiple imputation with principal components analyses missMDA v.1.17 (Josse & Husson, 2016), using MIPCA function with 5 components and visually inspecting the uncertainty associated to each variable on the corresponding factor map (Supp Fig NEW A1).

#### Variable selection

Since one of our primary goals was to obtain future projections of the niches, and we do not know of future soil layers, we performed an exploratory PCA with FactoMineR v.2.3 (Husson et al., 2016) and assessed the relative contribution of soil variables and their influence on the classes' clustering. Low soil variable contribution (exempting soil texture fraction clay -CLYPPT for wilds, Supp Table NEW 2) allowed us to continue our downstream analyses on climatic variables only. We conducted a correlation analysis with corrplot v. 0.84 (Wei & Simko, 2017) to select variable combinations with low correlations, and selected biologically relevant variables according to expert knowledge as recommended by (Rödder et al., 2009) (Supp Fig NEW A2). This decision was made to keep the models relatively simple because our priority was to compare general patterns among classes and not conflate differences among datasets due to differential input variables.

We selected nine variables: BIO2 = Mean Diurnal Range (Mean of monthly (max temp - min temp)), BIO3 = Isothermality (BIO2/(BIO7 = Temperature Annual Range), BIO4 = Temperature Seasonality (standard deviation x 100), BIO5 = Max Temperature of Warmest Month, BIO9 = Mean Temperature of Driest Quarter, BIO14 = Precipitation of Driest Month, BIO15 = Precipitation Seasonality (Coefficient of Variation), BIO18 = Precipitation of Warmest Quarter, and BIO19 = Precipitation of Coldest Quarter.

#### Outlier occurrences

With these variables, we proceeded to analyze our whole set of data points by scanning the resulting PCA for Mahalanobis outliers with FactoInvestigate v.1.7 (Thuleau & Husson, 2020) and ClassDiscovery function from the oompaBase v.3.2.9 (Coombes, 2019). Our climatic PCA results on the nine retained variables included none or few Mahalanobis outliers (FactoInvestigate v.1.7 and ClassDiscovery packages, respectively), which were carefully examined and found to be trusty presence data points.

### Maxent

#### Layer preparation

Prior to modeling, each domestication category dataset was spatially thinned five times at 5km with spThin v.0.2.0 (Aiello-Lammens et al., 2015) and the resulting mapped occurrences were compared. The nearest neighbor distance was selected to avoid spatial sorting bias within the dataset (Hijmans, 2012). We downloaded the previously selected environmental variable layers that averaged climatic data for the years 1970-2000 at 2.5 arc-min resolution from WorldClim (Fick & Hijmans, 2017). We used this resolution because the future layers were not available at finer grain at when we performed our analyses. We recognize that an ecophysiologically incomplete set of predictors may alter fine-scale predictions, but we were primarily interested in the comparison among domestication categories, including their climatic niches and general ranges under climate change, thus justifying our subset of climatic variables (Mod et al., 2016).

In addition, these present projection layers were cropped to a custom box (-123.0417, -80.91667, 8.875, 38.70833) chosen based on our knowledge of geographic scope of chile pepper in its native range (*C. annuum*). The afore-mentioned area box included all of Mexico, plus southern United States, Central America and northern tip of South America.

Future layer GCMs corresponded to different consortiums and countries climate assessments: one Beijing Climate Center Climate System Model (BCC-CSM2-MR), two Centre National de Recherches Meteorologiques (CNRM-CM6-1 and CNRM-ESM2-1), one Canadian Earth System Model -version 5 (CanESM5), one Institut Pierre-Simon Laplace (IPSL-CM6A-LR), two Model for Interdisciplinary Research on Climate – version6 (MIROC6) and Earth System version 2 (MIROC-ES2L), and one Meteorological Research Institute – Earth System Model version2 (MMRI-ESM2-0).

### Maxent tunning

MaxEnt model tuning is crucial to avoid overfitting and to gain realistic predictions of spatial range (Merow et al., 2013; Radosavljevic & Anderson, 2014). Inadequately simple or complex models will tend to perform poorly when inferring suitable habitats and consequently exhibit reduced transferability in time (Warren & Seifert, 2011). Model overfitting to spatially biased sampling should be avoided in presence-only modelling and can be reduced by the identification of optimal levels of model complexity for the species under study (Anderson & Gonzalez, 2011). We used ENMeval v.0.3.1 (Muscarella et al., 2014) to tune our MaxEnt modeling, as coded by the Wallace Ecological Modeling App (Kass et al., 2018). This allowed us to select the best combination of Feature Class and Regularization multiplier values *per* data set. Data was spatially partitioned using checkerboard2 (which includes fine and coarse-grain aggregation). Accounting for spatial partitioning is important to recover realistic distributions (Wisz et al., 2013). Feature Class combinations tested were L, LQ, LQH and LQHP (standing for L=linear, Q=quadratic, H=hinge and P=product, respectively). Regularization multiplier values tested were 0.5, 1.0, 1.5, 2.0, 2.5, 3.0 and 3.5. Varying regularization multiplier values allows us to relax models suspected of favoring a particular geographical area due to sampling bias, and thus may vary depending on the species under study (Anderson & Gonzalez, 2011) or domestication category in our case. Tuning results were plotted and compared within and among occurrence datasets (Supp Table NEW 4, Supp Fig NEW A3). The optimal combination for each dataset considered a compromise between low values for the corrected Akaike Information Criterion (AICc) and high average test Area Under the Curve (AUC). We refrained from selecting exclusively lowest AICc, because intermediately complex models may perform better when projecting across time (Moreno-Amat et al., 2015) and AIC seems to not correlate with predictive accuracy of the spatial distribution whereas AUC and TSS do (Velasco & González-Salazar, 2019). Medium to low omission rates (OR) were considered acceptable.

### Background tests

To run our background tests, we compared wild-landrace and wild-commercial pairs (adding a few Central America occurrences – Supp Table NEW A1). We first produced a mask for each category by intersecting the observed data points with Mexican physiographic regions using QGIS2.2 (QGIS Association, n.d.). For a given category, the area contained in its mask was then used to simulate its random distribution. This simulated distribution was then contrasted with other category's data points and D and I were calculated using ENMTools v.1.0.5 (Warren et al., 2010). The procedure was repeated 100 times for each pair in both directions. Results were plotted as histograms with D and I for each category pair marked to show whether the observed overlap is greater than 95% of the simulated values (Supp Fig NEW 8).

**Binary maps plotting**

For a given domestication category, the upper 10th quantile value of the distribution of training values was registered and averaged among all ten replicate runs. Binary maps were then plotted for each domestication category (Supp Fig NEW 5b) as well as pairwise overlap maps among categories (Fig 3, Supp Fig NEW 5c, Supp Table NEW A2).

In order to better follow the expected impact of time and mitigation strategies on niche suitability projections, we opted to intersect all eight GCM models per year-SSP combination (coinciding and sum of GCMs: Supp Fig NEW A4) and visualize the resulting map's overlap with present-time projections (Fig 5 and Supp Fig NEW A5) as well as pairwise overlap among domestication categories (Fig 6 and Supp Fig NEW A6). For each of these composite overlaps, pixel loss, gain and shifts were also calculated from raster files through custom R commands. (Supp Table NEW 5).

**MESS analyses**

We used dismo v.1.3.3 (Hijmans et al., 2020) to calculate MESS for each dataset projection -including present occurrence points values plus background points values- for each future scenario, to assess the extent of environmental differences involved in the projection (Supp Fig NEW 9). We also used terminal commands to perform MESS analyses for each future scenario *per se*, obtaining a geographical view of novel environments estimates and their novel limiting variables (those that are most novel at each pixel) (Supp Fig NEW 10). We were specifically interested in ensuring those areas predicted to shift for our domestication categories were not strongly climatically dissimilar in the future projections, in order to make more confident interpretations.
